# Defining Vaginal Community Dynamics: daily microbiome transitions, the role of menstruation, bacteriophages and bacterial genes

**DOI:** 10.1101/2023.06.02.543379

**Authors:** Luisa W. Hugerth, Maria Christine Krog, Kilian Vomstein, Juan Du, Zahra Bashir, Vilde Kaldhusdal, Emma Fransson, Lars Engstrand, Henriette Svarre Nielsen, Ina Schuppe-Koistinen

## Abstract

**Background:** The composition of the vaginal microbiota during the menstrual cycle is dynamic, with some women remaining eu- or dysbiotic and others transitioning between these states. What defines these dynamics, and whether these differences are microbiome-intrinsic or mostly driven by the host is unknown. To address this, we characterized 49 healthy, young women by metagenomic sequencing of daily vaginal swabs during a menstrual cycle. We classified the dynamics of the vaginal microbiome and assessed the impact of host behavior as well as microbiome differences at the species, strain, gene and phage levels.

**Results:** Based on the daily shifts in community state types (CSTs) during a menstrual cycle the vaginal microbiome was classified into four Vaginal Community Dynamics (VCDs) and reported in a classification tool, named VALODY: constant eubiotic, constant dysbiotic, menses-related and unstable dysbiotic. The abundance of bacteria, phages, and bacterial gene content was compared between the four VCDs. Women with different VCDs showed significant differences in relative phage abundance and bacterial composition even when assigned to the same CST. Women with unstable VCDs had higher phage counts and were more likely dominated by *L. iners*. Their *Gardnerella spp.* strains were also more likely to harbour bacteriocin-coding genes.

**Conclusions:** The VCDs present a novel time series classification which highlights the complexity of varying degrees of vaginal dysbiosis. Knowing the differences in phage levels and the genomic strains present allows a deeper understanding of the initiation and maintenance of permanent dysbiosis. Applying the VCD’s to further characterize the different types of microbiome dynamics qualifies the investigation of disease and enables comparisons at individual and population levels. Based on our data, to be able to classify a dysbiotic sample into the accurate VCD, clinicians would need two-three mid-cyclical samples and two samples during menses. In the future, it will be important to address whether transient VCDs pose a similar risk profile to persistent dysbiosis with similar clinical outcomes. This framework may aid interdisciplinary translational teams in deciphering the role of the vaginal microbiome in women’s health and reproduction.

## Background

The vaginal microbiota guards the entry of the reproductive tract. In concert with immune cells and the vaginal mucosa, it provides a physical and (bio-)chemical barrier against pathogens, preventing gynaecological infections. A healthy vaginal microbiome is dominated by non-pathogenic *Lactobacillus spp.,* producing lactic acid, hydrogen peroxide and bacteriocins, providing an acidic environment and hindering the growth of other bacteria^1^. In some women, the vaginal microbial composition can change suddenly, with a loss of *Lactobacillus* and the growth of other bacteria, often defined as vaginal dysbiosis^2^. A dysbiotic vaginal microbiome is considered ‘unhealthy’ as previous studies have associated it with poor reproductive outcomes, such as prolonged unexplained infertility, preterm birth, sexually transmitted infections (STI) and even gynaecological cancers ^3–8^. Understanding the dynamics of the intricate interplay between the vaginal microbiota and its environment is crucial for understanding how to maintain or improve women’s fertility and reproductive health.

Despite the epidemiological evidence, many unaccounted factors exist in defining a ‘healthy’ vaginal microbiome. A key issue is that this definition often lacks the temporal aspect: it remains to be determined whether there is a difference in the reproductive health between women with constant vaginal dysbiosis and women with fluctuations between *Lactobacillus* dominance and dysbiosis. *Lactobacillus* dominance can disappear abruptly, with high diversity as a result, but can sometimes quickly restore. So far, menstruation and sexual activity have been identified as primary drivers of these temporal changes^2^. The pattern of transitions in the vaginal microbiome over time in any individual woman is a complex interaction between three main determinants: inherent causes (genetic, immune system, hormone levels), lifestyle/clinical drivers (sexual intercourse, bleeding, hygiene habits) and microbiome determinants (for example interactions between species or strain-level differences)^9,10^. How readily a microbiome recovers from dysbiosis may depend on which of these determinants initiates the dysbiosis.

Whether a sudden lack of *Lactobacillus* dominance is due to a loss of *Lactobacillus spp.* that favours the growth of other bacteria or whether other bacteria can suppress the *Lactobacillus spp.* dominance is still a pending question^11–13^. A considerable reduction of *Lactobacilli spp.* that occurs before the expansion of anaerobic bacteria typical of bacterial vaginosis (BV) could be caused by bacteriophages. Lysogenic phages reside inside the bacterial host, and this viral strategy is probably favourable when the density of its host bacteria is low. Notably, phages can rapidly switch from a lysogenic to a lytic cycle, quickly killing the host bacteria and releasing thousands of phage virions ^14,15^. However, only a few studies have focused on the viral players of the vaginal ecosystem. Therein, it was shown that bacteriophages in vaginal swabs, like bacteria, cluster into two unique bacteriophage community groups: a high-diversity and a low-diversity group^16^. These two bacteriophage community groups correlated with the *Lactobacillus* dominance (low diversity bacteriophages) and non-*Lactobacillus spp.* dominance (high diversity bacteriophages) bacterial groups. Moreover, the bacteriophage composition may predict clinical BV as efficiently as the bacteriome composition^17^.

The main drivers necessary to induce the overgrowth of certain bacteria, such as *Gardnerella spp.*, *Prevotella spp.* or *Fannyhessea vaginae*, into full-blown bacterial vaginosis are still poorly understood. Contributing factors such as biofilm formation, local inflammation, and endocrine differences in the individual may all contribute^18^. While genus *Gardnerella* has long been associated with BV, its recent delineation into 13 genomic species^19^ has led to a better understanding of its genomic variability and association with disease^20^. Still, little is known about the pangenome of these species, considering that their genetic variability and accessory genes could be significant in understanding the development of bacterial vaginosis and its role as a risk factor for poor reproductive outcomes and wome’s health.

In most studies, determining whether a vaginal microbiome is ‘unhealthy’ is based on its composition on the arbitrary day the sample was taken. Therefore, the definition of an unhealthy vaginal microbiome in women of reproductive age calls for more precise terminology, both in terms of specific patterns of species abundance and in the timing and duration of the dysbiosis in relation to the menstrual cycle or pregnancy. There is a great need for a thorough investigation of the daily transitions in the vaginal microbiome with detailed information on both lifestyle and microbiome features to support future research by defining when and how often a woman needs to be sampled for accurate characterization of her vaginal microbiome.

In the pilot phase of this study 15 women were followed with daily swabs for 42 days. Then, the daily changes in the vaginal microbiome were investigated by shotgun sequencing in 49 young, healthy women during a menstrual cycle to identify potential drivers of sudden transitions in microbiome composition. Based on these data, we aimed to classify the dynamic patterns of the vaginal microbiome composition during a menstrual cycle into Vaginal Community Dynamics (VCDs). The metagenomic approach made it possible to analyze the presence of co-occurring bacteriophages and, by metagenomic assembly, investigate the different bacterial genomic strains that are connected to dysbiosis.

## Results

### Daily vaginal swabs reveal both rapid and cyclic changes in the microbiome

In the pilot stage of this study, the daily vaginal microbiome of 15 women for up to 42 days was sequenced using V3-V4 tags from the 16S rRNA gene. These women were using three different contraceptive regimens (non-hormonal contraceptives, NHC; combined oral contraceptives, COC; levonorgestrel intra-uterine system, IUS; n = 5 in each group). The samples generated 839 ASV, corresponding to 154 species from 130 genera. The full dataset is presented in **Supplementary table 1**. Some women, such as ID 15, 71, 150, present significant fluctuations daily, including rapid changes in alpha-diversity (**fig. 1**). Conversely, participant ID 11 or 139 have essentially constant diversity, and 91 has stable profiles throughout the entire cycle. Despite these fluctuations, we observed the effect of menses in women with regular menstrual bleeding. Menstruation in participant ID 11, 15, 29 is repeatedly accompanied by an increase in *Lactobacillus iners, Prevotella spp* and *Sneathia spp,* respectively. Participant 46 is an excellent example of a reversible change during menses, with a rapid expansion in *Gardnerella spp*, quickly followed by dominance of *Lactobacillus spp.* Conversely, participant 51 shows how changes triggered by menses can become permanent. A significant expansion of *Sneathia amnii* in parallel with loss of *Lactobacillus iners* results in a *Gardnerella vaginalis* dominated microbiome until the end of the follow-up.

**Figure 1:**
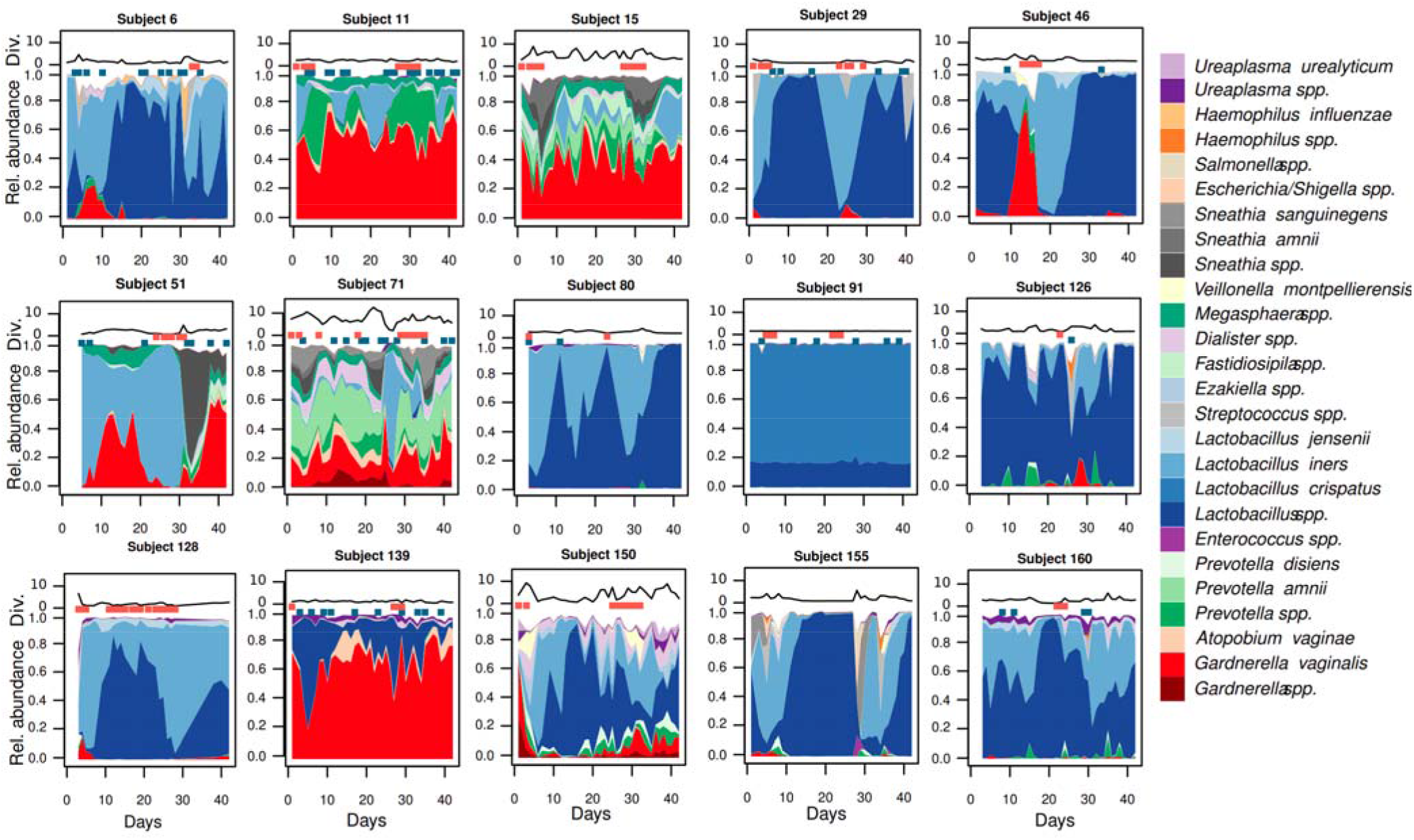
The vaginal microbiome can be remarkably stable over 6 weeks, but also experience both cyclical and rapid shifts. Area plots represent bacterial composition inferred from 16S rRNA gene amplicons, with relative abundance on the Y-axis and days in the X-axis. Red dots above the area chart represent days with menstrual bleeding or spotting, and blue dots represent days with vaginal intercourse. The black line above each profile shows their alpha-diversity (inverted Simpson’s index).

Since the pilot stage showed the need for daily samples to capture the whole dynamics of the menstrual cycle, additional 49 women were included with daily vaginal swabs from day 4 to day 32 of their menstrual cycle. Metagenomic sequencing was used in the larger sample set to improve taxonomic and functional resolution and extend our analysis to viruses.

### Characteristics of the main study population

For the main study, samples from forty-nine women were included using metagenomic shotgun sequencing. An average of 25.9 samples were successfully sequenced, starting from cycle day four with daily samples for another 28 days. The women were generally young, lean and healthy, and the majority had had a single sexual partner in the previous month. Detailed health and demographic information are presented in **table 1**, divided into the proposed vaginal time-series microbiome classification presented below.

**Table 1:**
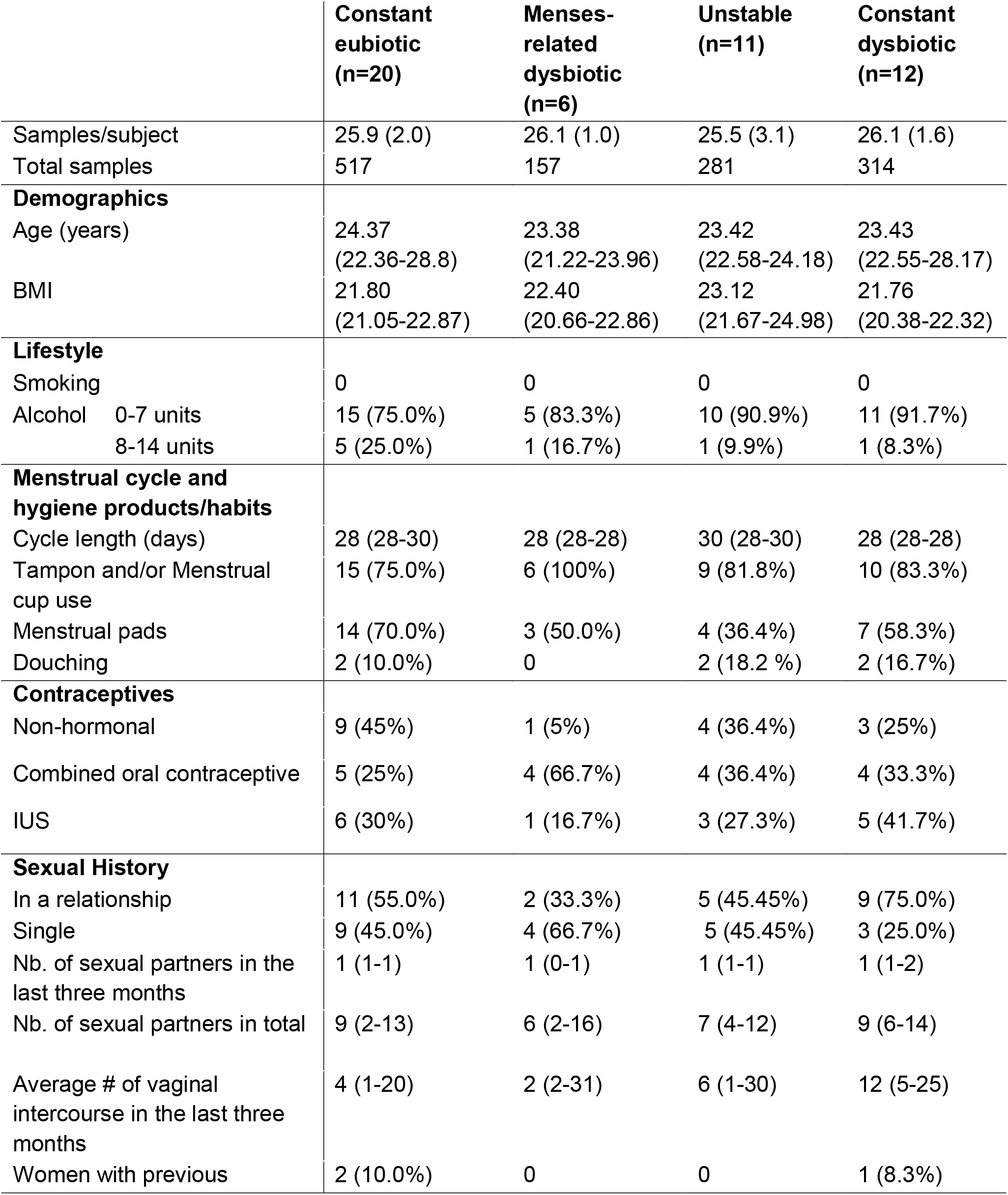
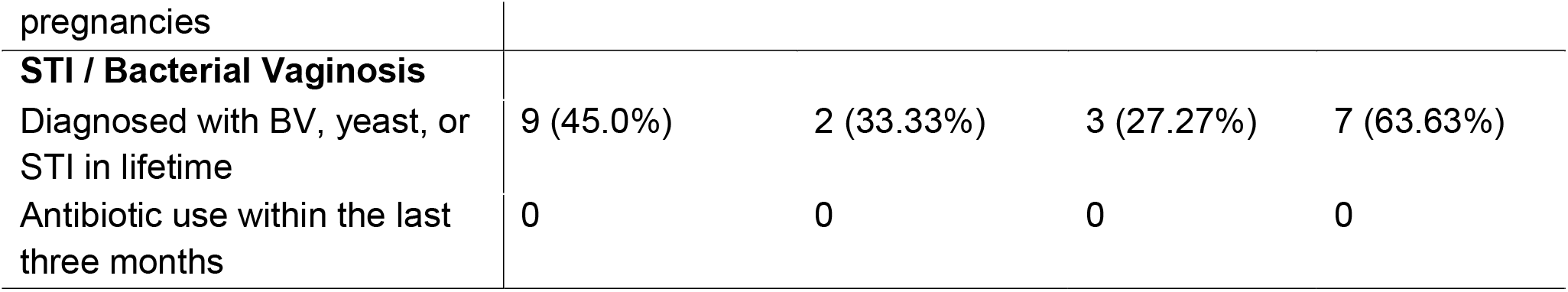
Clinical characteristics of the participants. Prevalence is given as n (%) and other values as means (SD)

### Unprotected intercourse and menstrual bleeding negatively affect the stability of the vaginal microbiome

The 1269 samples sequenced by shotgun were, like the 532 metabarcoding samples, dominated by *Lactobacillus crispatus, Lactobacillus iners, Gardnerella spp.* and *Prevotella spp*. The total beta-diversity during the time series fluctuated considerably in the individual woman. Four representative samples are shown in **fig. 2:** The vaginal microbiome of participant 156 is *L. crispatus-*dominated and shows no response to either intercourse or bleeding. Participant 48 has no *Lactobacillus spp,* and is instead dominated by *Gardnerella spp, Prevotella spp* and *Peptoniphilus lacrimalis.* Her vaginal microbiome is changing rapidly throughout the study period. Participant 115 is dominated by *Lactobacillus crispatus* in most samples, but during menses, becomes dominated by *Gardnerella spp.* Finally, participant 34 is *Lactobacillus crispatus* dominated but presents rapid shifts in response to intercourse. All participants are shown in **supplementary figure 1 and supplementary table 2.** Interestingly, except for subject 120, *Sneathia spp.* is only observed in close proximity to menstrual bleeding.

**Figure 2:**
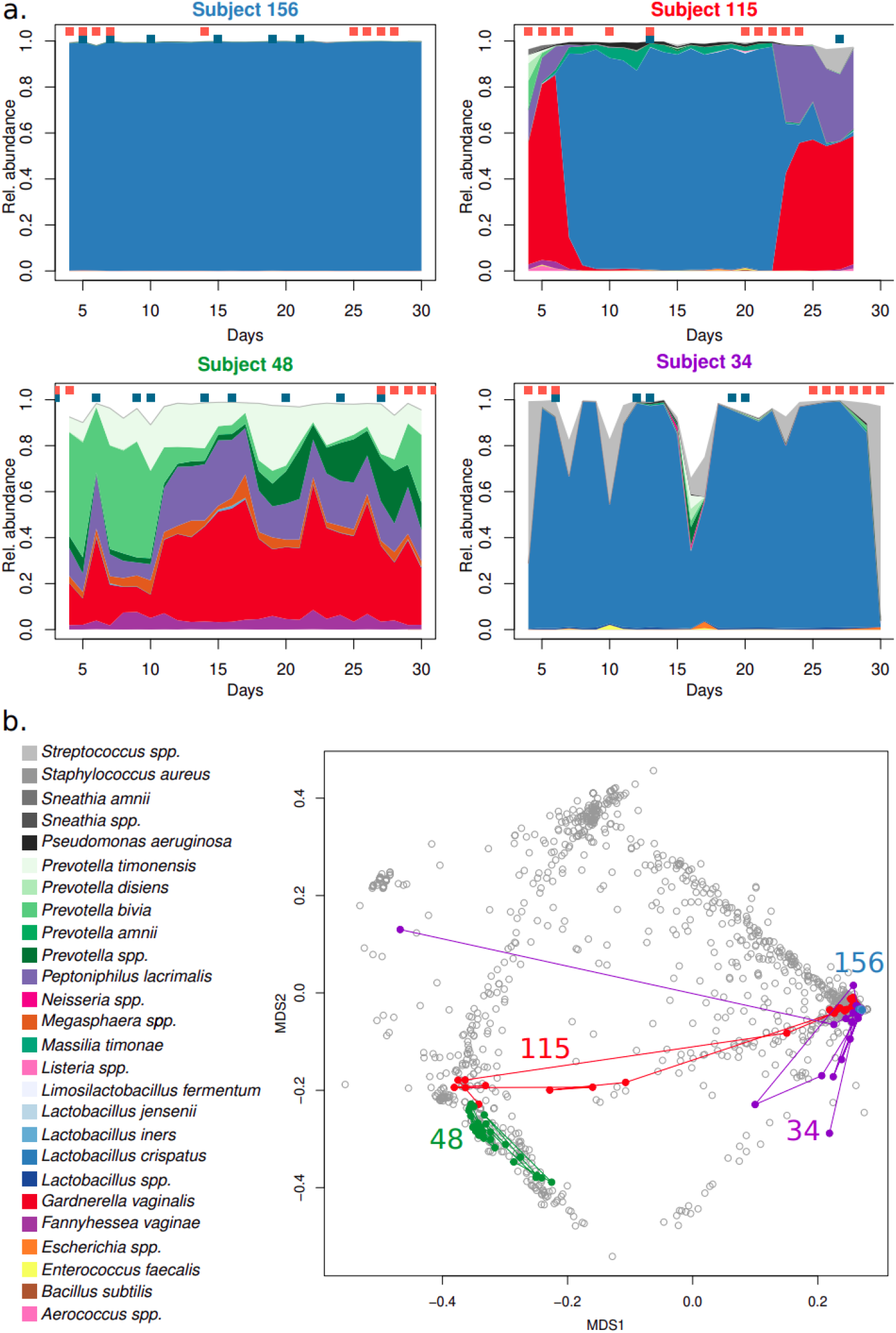
Vaginal samples are dominated by either *Lactobacillus spp, Gardnerella spp.* Or *Prevotella spp.,* and can rapidly or cyclically switch between types. a. Four representative individuals’ vaginal microbiome are shown during a menstrual cycle, starting from cycle day 4. Women can be stably high *Lactobacillus spp,* stably *Lactobacillus* spp depleted, high *Lactobacillus spp* except during their menses or high *Lactobacillus spp* but with relative abundances falling as a response to unprotected sexual intercourse. b. a non-metric multidimensional scaling based on Bray-Curtis dissimilarity of all shotgun metagenomics samples in this study. The same individuals are highlighted, showing their trajectory during the follow-up.

The stability of the vaginal microbiome may result from features unique to the microbiome or host. We, therefore, started assessing the correlation between total beta-diversity and host-specific factors. Changes in microbiome composition can be divided into two main categories, qualitative (the introduction or removal of species) and quantitative (alterations in the relative abundance of species present). To assess these key components, the total beta-diversity observed in a time series, normalized to the number of samples successfully sequenced, was calculated as either the qualitative Jaccard’s or the predominantly quantitative Aitchinson’s distance. Contraceptive usage did not affect normalized beta-diversity with the Jaccard distance (all p > 0.3). However, Aitchinson’s distance was higher in the non-hormonal contraceptive group than in the combined oral contraceptive or progestin-only intra-uterine system (**fig 3a**). Aitchinson’s and Jaccard’s distances were weakly correlated to the total bleeding and spotting days (**fig 3b**). However, only Aitchinson’s was strongly correlated with the total number of intercourses, indicating a (temporary) quantitative change in beta-diversity (**fig 3c**). Conversely, only a qualitative change in beta-diversity as measured by Jaccard’s was correlated to days with menstrual bleedings -after the removal of participants with an IUS (**fig 3d**). No significant differences in total beta-diversity were found in relation to menstrual hygiene products or douching.

**Figure 3:**
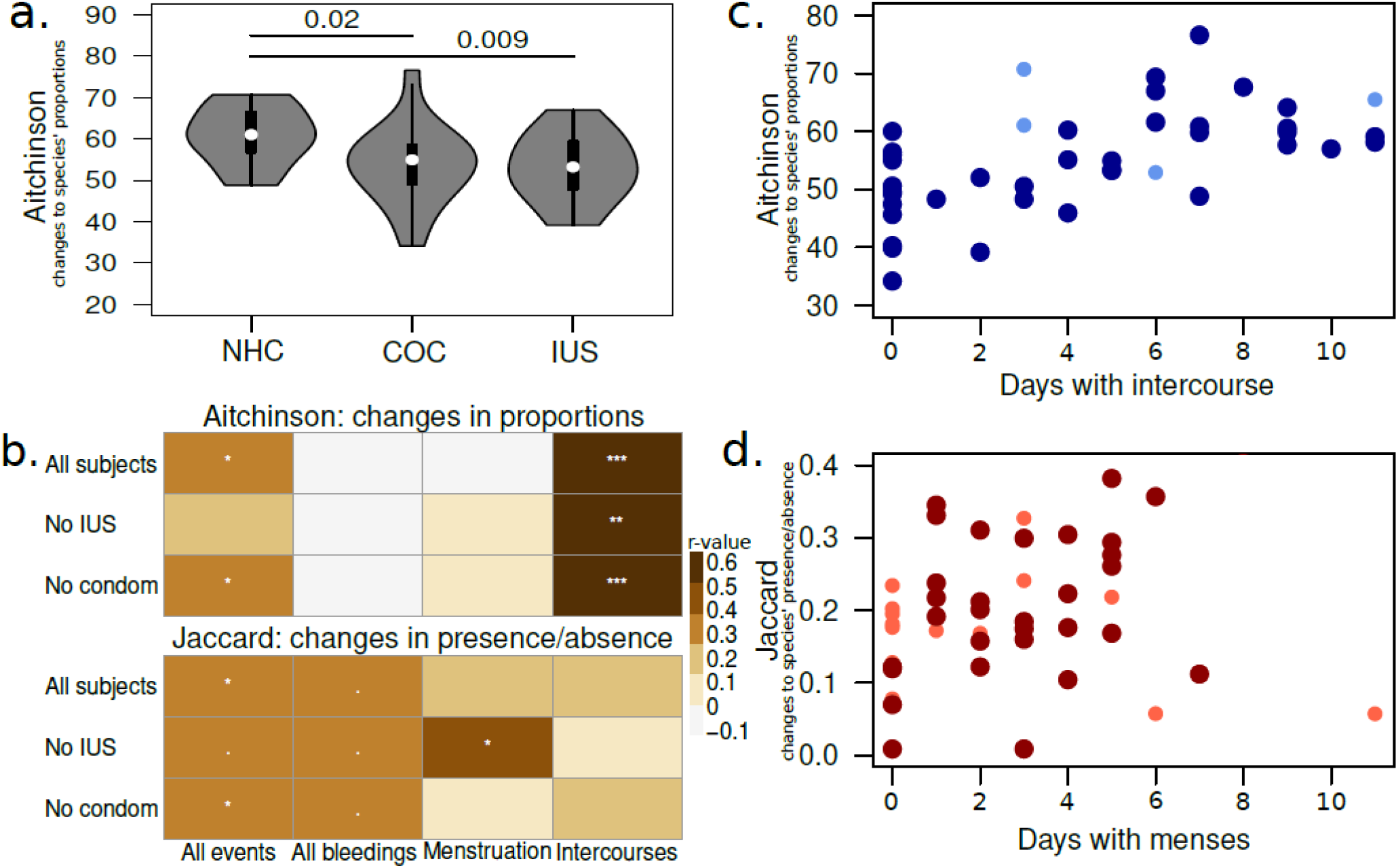
Contraceptive usage and intercourse frequency affect relative abundance of bacteria, while days with menses affects the influx of new bacteria. a.Women not using hormonal contraceptives have higher total beta-diversity over the sampling period than women on combined oral contraceptives or with an IUS. b. Pearson’s correlations between life events (bleedings, intercourses) and total beta-diversity, Aitchinson’s or Jaccard’s, per subject in different groups c. Number of days with unprotected sexual intercourse is directly correlated to total Aitchinson’s distance. Dark blue dots: women with unprotected sexual intercourse. Light blue dots: women with sexual intercourse with condoms. d. Number of days with menstrual bleeding is directly correlated to total Jaccard dissimilarity. Dark red dots: women not using hormonal contraceptives or on combined oral contraceptives. Light red dots: women with an IUS, with typically very light bleeding.

In the same way that community state types (CST) have become a standard for the research community, we propose a standard for classifying the dynamics of a vaginal microbiome. We classify vaginal samples during a complete menstrual cycle based on CST into four Vaginal Community Dynamics (VCD). The two first categories consist of “constant eubiotic” and “constant dysbiotic” if the woman is either eubiotic or dysbiotic in more than 80% of the daily samples during a complete menstrual cycle. Then for the two intermediate VCDs, we assess the remaining samples’ dynamics when samples are not under the influence of menses. If a woman is having >80 % eubiotic samples only during the cycle days 9-25 her vaginal microbiome is classified as "menses-related dysbiotic” and, otherwise, as “unstable” (**fig.4**). In this study, we have defined eubiosis as CST-I and CST-V and dysbiosis as CST-III and CST-IV. CST-II was not observed in this cohort (**fig. 4)**. These parameters can be adjusted for the specific population studied and the experimental set-up. Code is available from www.github.com/ctmrbio/valody. Using this scheme, we find that in 49 women: 20 are constant eubiotic, six have menses-related dysbiosis, 11 are unstable, and 12 are constant dysbiotic (**fig.4)**. The CSTs and VCDs for the entire investigated population in this study are shown in **supplementary fig.2** (16S pilot) and **supplementary fig.3** (main study).

**Figure 4:**
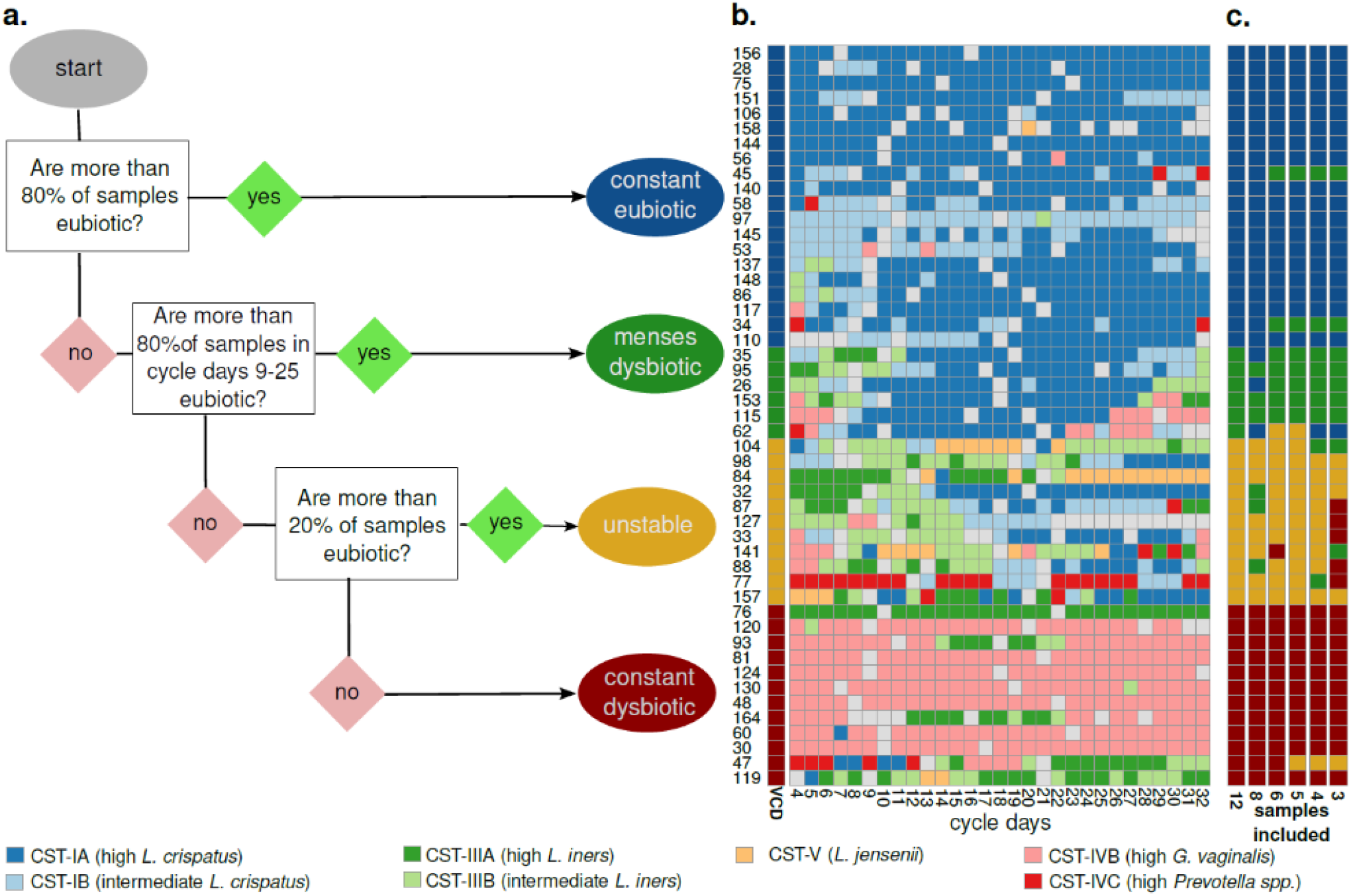
Vaginal time-series can be classified into four categories (Vaginal Community Dynamics) according to their proportion of eubiotic samples. A decision tree can separate a time-series of samples into dynamic groups, based on the community state-types. Input from the user is which CST are considered eubiotic (here: I, II and V) and which days are to be considered free from the influence of menses (here: cycle day 9 to cycle day 25). Time-series with >=80% eubiotic samples are considered constant eubiotic; conversely, those with >80% dysbiotic samples are considered constant dysbiotic. For those in the 20-80% range, a second assessment is done on the days free of menses: if they’re >80% eubiotic, the time-series is considered menses-related dysbiotic, and otherwise unstable (changing from eubiosis to dysbiosis without a clear temporal pattern). b. A color map with one individual per row and one day per column. The color of each intersection depicts CST. Colored bars on the left side show the vaginal community dynamics of each woman. c. Additional color bars show the inferred vaginal community dynamics of each subject when using fewer samples for classification.

To increase the applicability of the VCDs, we also assessed how many samples during a menstrual cycle are necessary to assign them to the correct VCD accurately. Sampling every other day still gives perfect accuracy. However, separating menses-related dysbiosis from an unstable vaginal microbiome requires at least five samples, two during menses and three outside. Meanwhile, constant eubiotic dynamics can be detected with two eubiotic samples, at least one during menses, and constant dysbiosis requires 2-3 mid-cycle dysbiotic samples **(fig.4)**.

### Samples from different Vaginal Dynamic Types show large differences in bacterial content despite having the same CST

One factor that can affect microbiome communities’ stability is bacterial species that perform critical ecological services despite being in low abundance. To assess whether this was likely the case for our samples, we used ANCOM-BC2 to assess whether samples from different VCDs differed in their bacterial composition. Because the VCDs are based on CST proportions, we only compared samples within the same CST subtype. Additionally, in each comparison, we only included dynamic groups with at least ten samples from at least three different women to minimize the effect of individual outliers. This way, we could compare the menses-dysbiotic and unstable groups to constant eubiotic in CST-IA and CST-IB and constant dysbiotic in CST-IIIA, CST-IIIB and CST-IVB.

Regarding the highly eubiotic CST-IA, samples from menses-dysbiotic and unstable subjects have more *L. iners, Gardnerella spp.* and *Ureaplasma urealyticum* than constant eubiotic individuals. The unstable individuals also had an increased abundance of *Campylobacter spp, Corynebacterium spp* and *Streptococcus spp.* Interestingly, the eubiotic samples had two-fold higher *E. coli* than both the unstable and the menses-dysbiotic samples. In CST-IB, the results were similar, albeit the non-stable time series had an even lower abundance of certain BV-associated bacteria. Specifically, samples from menses-related dysbiotic women had a lower abundance of *P. amnii, P. bivia and P. disiens,* while the unstable time series had a lower abundance of *Fannyhessea vaginae* and *Corynebacterium spp.*, in addition to decreased *E. coli.* The top 30 most extreme fold-changes are shown in **fig.5a** (for the complete results, see **suppl. Fig.4 and suppl. Tables 3 and 4)**

Focusing on CST-III, the unstable and menses-dysbiotic had a higher relative abundance of several *Lactobacillus* species than the constant dysbiotic, most notably *L. crispatus* and *L. gasseri*, *L. vaginalis, L. reuteri and Limosilactobacillus spp.* Additionally, in CST-IIIB, the menses-associated individuals had decreased abundance of several *Prevotella spp, Veillonela spp, Megasphaera spp, and Mobiluncus spp* and the BV-associated Clostridiales KA00067. The top 30 extreme fold-changes are shown in **fig 5b** (complete in **suppl. Fig. 4 and suppl. Tables 5 and 6).** For comparison, too few menses-dysbiotic samples was assigned CST-IIIA. No significant results were found comparing samples in CST-IVB.

**Figure 5:**
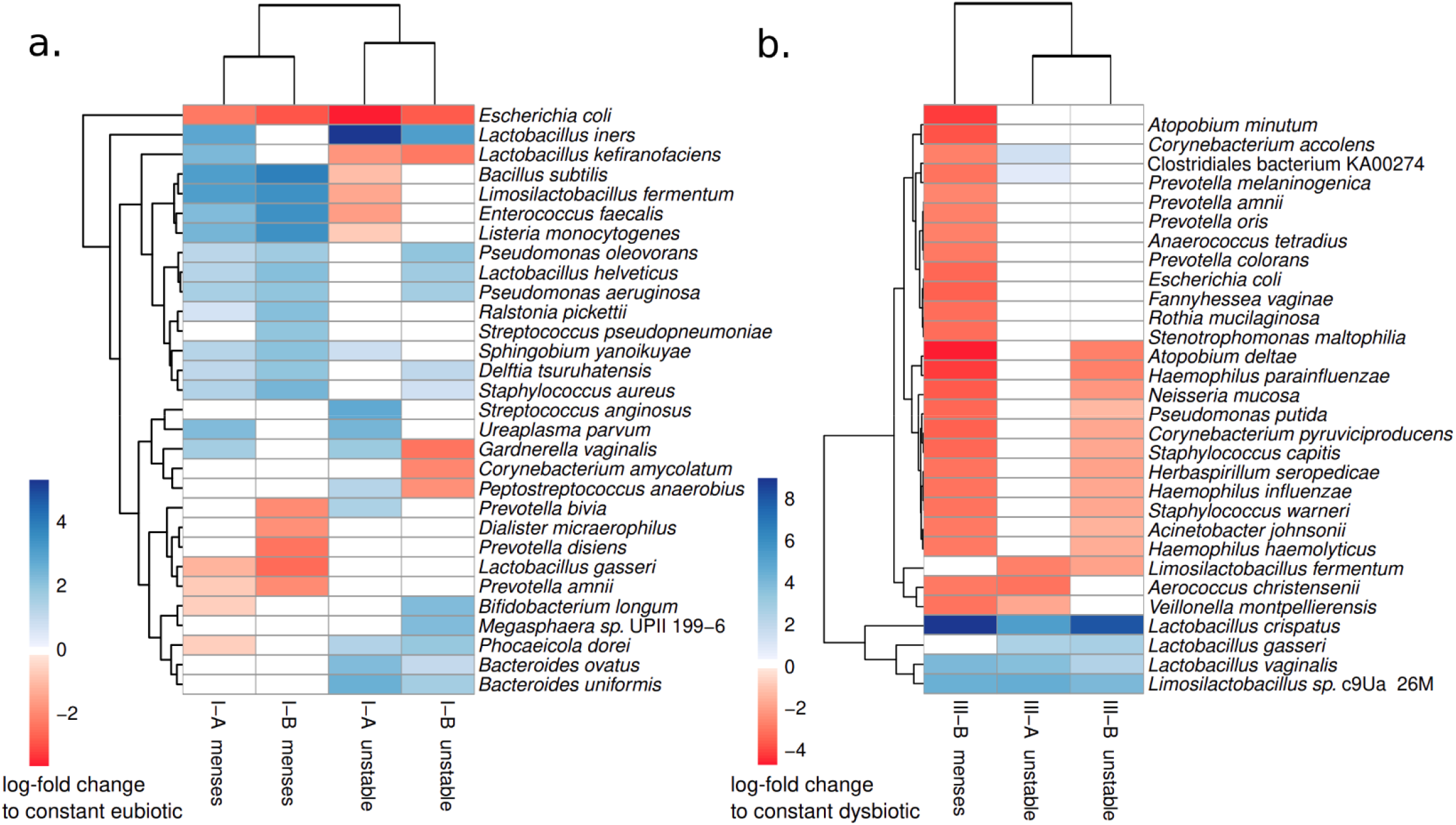
Samples belonging to the same CST, but deriving from different dynamic groups, have changes in the relative abundance of several bacterial species. a. Samples in CST-IA and CST-IB from menses-related dysbiotic or unstable individuals were compared to those from stable eubiotic individuals. b. Samples in CST-IIIA and CST-IIIB from menses-related dysbiotic or unstable individuals were compared to those from stable dysbiotic individuals. In each panel, the heatmap shows the log-fold change of the top 30 most extreme differences. White fields represent no significant change.

As a complementary analysis to the one above, we also assessed differentially abundant bacteria between dynamic groups not separated by CST, using either the constant eubiotic or the constant dysbiotic as the baseline for comparison. Both menses-dysbiotic and unstable had more differentially abundant bacteria compared to the constant dysbiotic time-series than when compared to eubiotic (menses-dysbiotic: 41 from constant eubiotic, 136 from constant dysbiotic; unstable: 81 from constant eubiotic, 128 from constant dysbiotic). In common, both intermediate groups have more *L. iners,* more *S. agalactiae* and more *Ureaplasma parvum* than the constant eubiotic (**suppl. Fig.6; suppl. Table 7)**. Conversely, compared to the constant dysbiotic, both of the intermediate VCDs have decreased relative abundance of both *Prevotella spp* and BV-associated bacteria, but an increase in *Bifidobacterium spp, Lactobacillus spp, Staphylococcus spp* and *Streptococcus spp (***suppl. Fig.6; suppl. Table 8)**. While roughly confirming what was expected, these analyses were not significant when adjusted for either individual ID or sample CST and must therefore be interpreted with caution.

### Common and abundant species display considerable differences in functional gene content, but no association with microbiome stability

In addition to bacteria in low abundance, it is possible that what determines the dynamics of a community are specific strains and their accessory functional genes. The time-series design of this study also allowed us to build draft genomes (MAGs, metagenome-assembled genomes) for prevalent and abundant bacteria within each participant. At the read-level analysis, all *Gardnerella spp.* reads were classified as *G. vaginalis.* However, genome assembly revealed at least three species in this small cohort, namely *G. leopoldii*, *G.piotii* and *G. vaginalis*, of which the latter is the least common (**fig 6a**).

**Figure 6:**
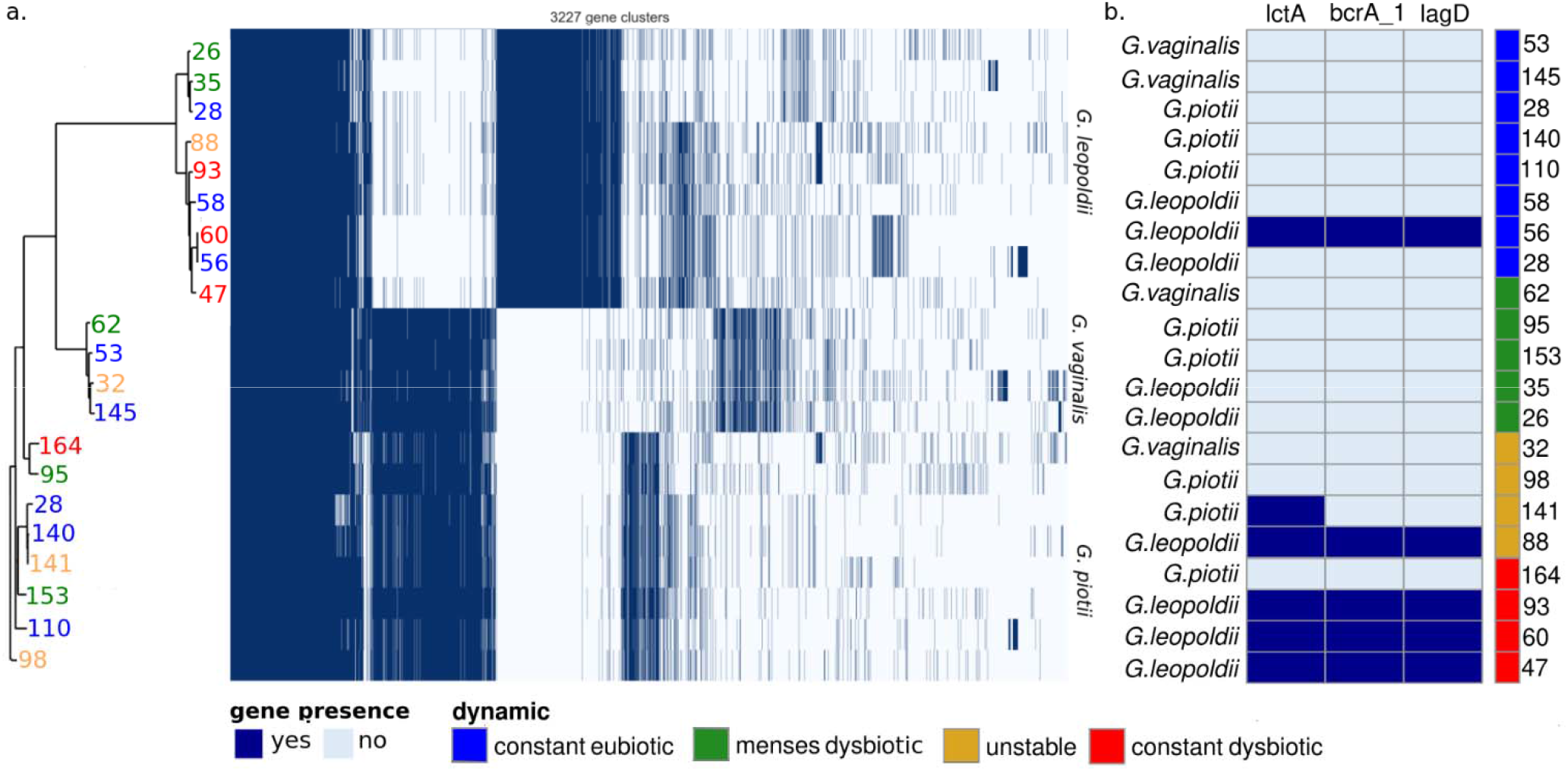
While strains do not segregate by vaginal community dynamics, bacteriocins are associated with instability. a. Phylogenomic analysis of all detected *Gardnerella* species does not find a correlation between the individuals’ vaginal community dynamics and the observed phylogeny. b. Three bacteriocins from *G. leopoldii* are over-represented in unstable and dysbiotic samples. Presence of a gene is represented in dark blue and its absence in light blue. Blue: women who are constant eubiotic. Red: women who are constant dysbiotic. Yellow: women with unstable VCD. Green: women who are menses-related dysbiotic.

To assess strain-level differences, we focused on the genera which yielded the largest number of high-quality MAG, i.e. *Gardnerella spp., Lactobacillus spp.* and *Prevotella spp*. The three *Gardnerella* species had a relatively small pangenome, unlike the large *Prevotella* pangenomes. Among *Lactobacillus spp, L. crispatus* had by far the largest pangenome, while *L. iners* had the smallest (**supplementary fig. 7**). Most gene families were classified as either “core” (present in all or almost all genomes) or “cloud” (present in one or very few genomes), with fewer clusters in the intermediate “shell” category. Because of this distribution pattern, no gene cluster could be significantly associated with a VCD **(figures S8-S9; Tables S9-S11)**.

Still, while not significant past multiple testing corrections, we did find three bacteriocin genes associated with *G. leopoldii* that were over-represented in unstable or constant dysbiotic subjects (**fig 6b).** While the statistical support for this finding is weak, *Gardnerella* bacteriocins are known to affect *Lactobacillus spp. in vitro*.

### Bacteriophages stabilize vaginal microbiomes, both in eubiosis and dysbiosis

Bacteriophages are also known to affect the stability of bacterial communities directly, sometimes very rapidly. In this cohort, the taxonomic profile of phages mostly followed their host bacteria (**fig.7**). This was expected, since samples were not enriched for viral particles, so presumably, most viral DNA originates from internalized or integrated phages. Still, we also detected a variety of phages associated to species more typical of the skin, such as *Streptococcus spp, Staphylococcus spp* and *Propionibacterium spp* (**supplementary fig.1**).

**Figure 7:**
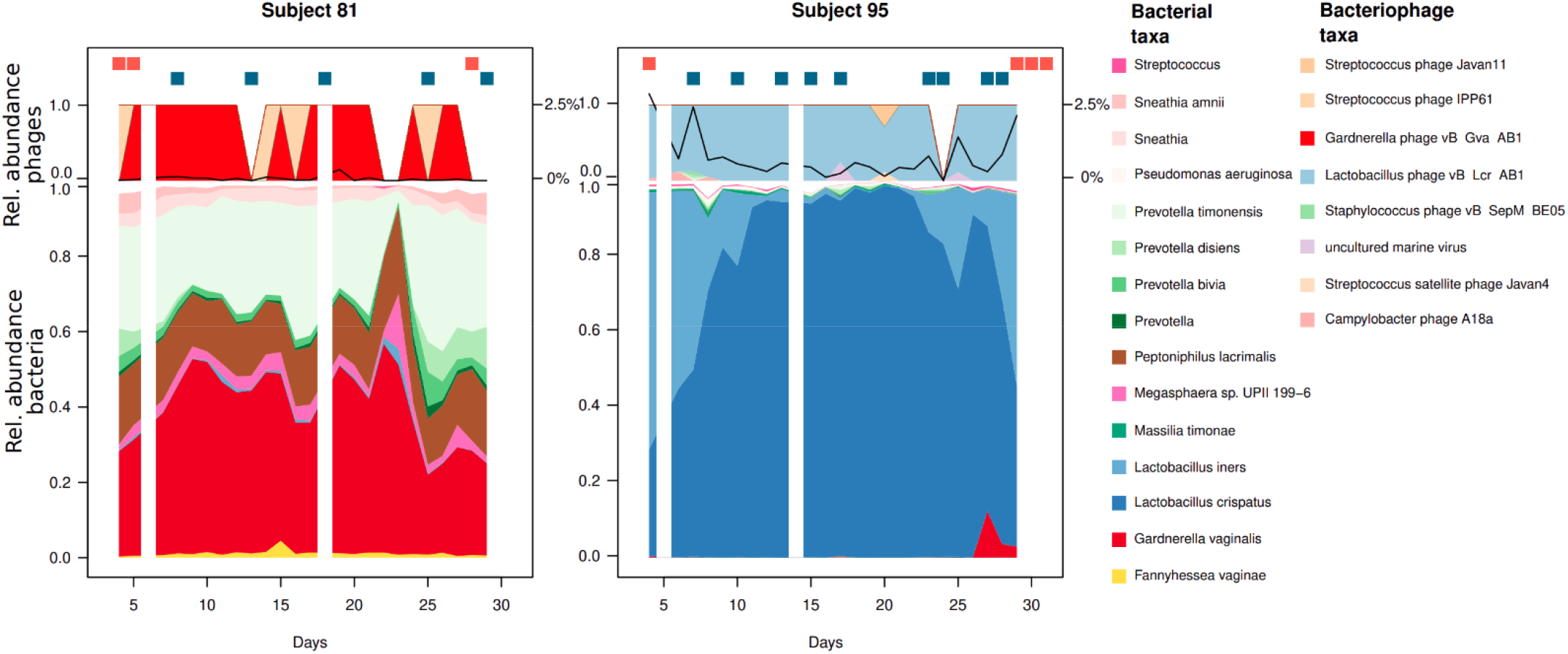
Phage profiles follow roughly the bacterial profiles but can fall below detection limit in samples of lower coverage. Two representative individuals’ vaginal bacteriomes and phageomes are shown during a menstrual cycle, starting from cycle day 4. Red dots above the area chart represent days with menstrual bleeding or spotting, and blue dots represent days with vaginal intercourse. The black line overlapped to the phage profiles represents the ratio between phage reads and bacterial reads. Days with missing data are omitted.

The ratio of phage read counts to bacterial read counts was highly variable, ranging from the detection limit of c. 10E-05 up to 2.7%. Phage counts typically spiked when samples deviated from their most prevalent CST. Samples with high phage ratios had a larger beta-diversity than their previous and following samples, with Jaccard’s and Aitchinson’s distances (Spearman’s rank correlation: Aitchinson’s, rho=0.09, p=0.02; Jaccard, rho=0.13, p=0.0002).

Phage ratios were also significantly different between VCD and, within them, CST. The constant dysbiotic samples had the lowest phage counts overall, c. 10-fold lower than all others. Within CST-I and CST-III, the unstable VCD had the highest phage counts, while menses-related dysbiotic only had a higher phage ratio in CST IV-B (**fig.8)**.

**Figure 8:**
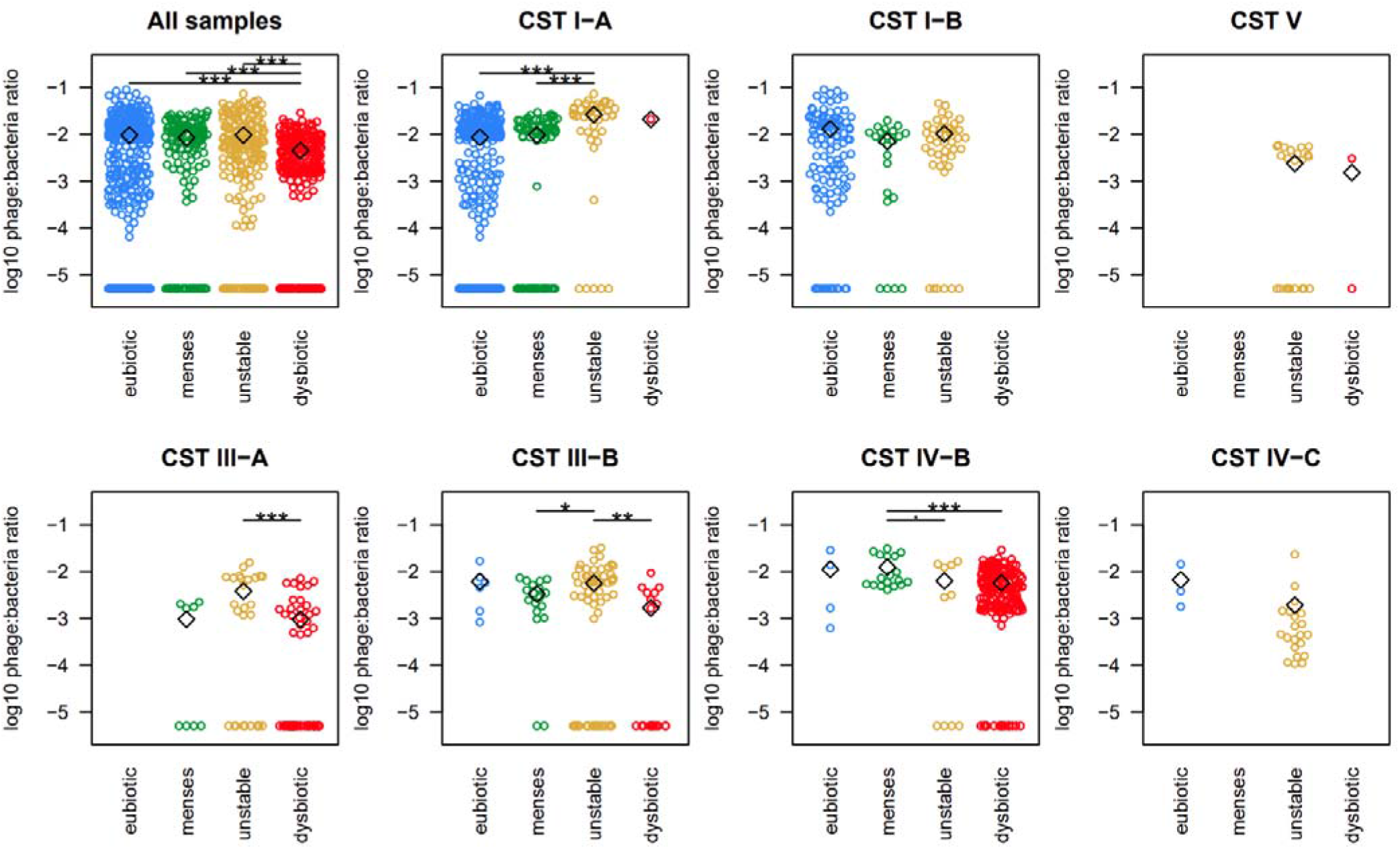
Relative abundance of phages is connected to CST and individuals’ dynamics in ways that promote eubiosis. The Y-axis in each plot represents log10 of the ratio between phage reads and bacterial reads. Each open circle is a sample, open diamonds are means. a. The unstable women have the lower overall phage count over the entire time-series b. For samples with menses-related dysbiosis only, the dysbiotic samples present 100-fold increased phage counts compared to their eubiotic samples. Blue: constant eubiotic Red: constant dysbiotic Yellow: unstable Green: menses-related dysbiotic.

## Discussion

The main aim of this study was to identify and map the daily variations in the vaginal microbiome composition in healthy, young, Caucasian women during an entire menstrual cycle. Several groups have tried to define types of vaginal compositions in women, only to ascertain an extreme variation between individuals. One of the pivotal keys when refining our understanding of the categorization of the vaginal microbiome is to evaluate the composition dynamics during the menstrual cycle in each woman. As shown in this study, the vaginal microbiome composition can change rapidly, from day to day, and with marked alterations primarily driven by external factors such as menstrual bleeding or sexual intercourse, with both exposures also reported in previous studies^21,22^.

In contrast to previous works, we tried to identify the difference between women who constantly remain eubiotic or dysbiotic during an entire menstrual cycle. Since previous work did not fully account for the dynamic patterns in vaginal microbiome composition, it is unknown whether women with transient (menses-related or unstable) dysbiosis present a lower risk of reproductive health complications than women who are constantly eubiotic. The transient dysbiotic phenotype could reflect a stepwise increase in severity, with menses-related and unstable dysbiosis being potential precursors of eventually transiting into unhealthy dysbiosis. This study defined four Vaginal Community Dynamics (VCDs): 1) the constant eubiotic with non-*iners Lactobacillus spp.* dominance throughout the cycle and an apparent resilience against exposures, 2) the menses-related dysbiotic with a sudden drop in *Lactobacillus spp.* dominance during menstruation, 3) the unstable dysbiotic which changes community states for a short while, for example, after sexual intercourse, and reinstate *Lactobacillus spp.* dominance, 4) the constant dysbiotic, characterized by an overgrowth of typical BV-associated bacteria such as *Gardnerella spp*, *Prevotella spp.* and *Fannyhessea vaginae* throughout the menstrual cycle. Women with menses-related dysbiosis present a qualitative change in their microbiome composition during menses, with the introduction and removal of several species. In this period, there is an overgrowth of *Gardnerella spp* or *L. iners, Prevotella spp* and *Sneathi<u>a</u> spp*, with conversion back to *L. crispatus* dominance mid-cyclical. Alterations in the mucosa could be driving this pattern, perhaps caused by decreasing estradiol levels. However, we also see a direct correlation between the number of days with menstrual blood and Jaccard’s distance, suggesting an essential role of menstrual blood providing bacterial nutrients and increasing the vaginal pH value. Another interesting observation in this study is that the expansion of *Sneathia spp*. was almost only observed immediately after menstrual bleeding. *Sneathia spp*. is an opportunistic pathogen that has been associated with adverse pregnancy outcomes such as premature rupture of membranes, preterm birth and chorioamnionitis^23^. It was recently found that *Sneathia amnii* produces an exotoxin that can haemolyze erythrocytes and break down cellular barriers^24^. Additionally, the higher level of phages in the dysbiotic samples (CST-IVB) observed in women classified as menses-related VCD could contribute to the rapid eradication of BV-associated bacteria after menses. Due to methodological constraints, our analysis of phages focused on their abundance rather than their taxonomy. Nevertheless, these findings show that vaginal bacteriophages are linked to the bacterial community’s stability, consistent with previous research on bacteriophages and the vaginal microbiota^17,25^.

Women with unstable VCD may be particularly susceptible to pH changes, allowing a temporary overgrowth of anaerobic bacteria. Alternatively, this could be explained by a direct introduction of non-vaginal species by their sexual partner. Both scenarios would explain a strong correlation between total beta-diversity (measured as Aitchinson’s distance) and the number of intercourses. Moreover, the unstable group showed a higher prevalence of *L. iners*, which is associated with the formation of vaginal biofilms in women with bacterial vaginosis (BV)^26^. This biofilm could make the affected women more prone to an unstable vaginal microbiome, eventually leading to constant vaginal dysbiosis. Women classified as unstable VCD also had higher phage counts in CST-IA and CST-III. The higher phage count could contribute to a kill-the-winner dynamic, where neither *L. crispatus* nor *L. iners* can dominate the microbiome before having their population killed by phages^27^. Further research is needed to explore the role of bacteriophages in the development, maintenance, and recovery from dysbiosis.

The constant dysbiotic VCD is characterized by a highly diverse composition with many aberrant bacteria throughout the menstrual cycle. The critical difference between persisting dysbiosis and the two VCDs with a varying degree of dysbiosis (unstable and menses-related) could be the establishment of a resistant polymicrobial biofilm in the former. The biofilm can consume nutrients, produce metabolic waste products, and create an altered microenvironment that is more permissive to the growth of harmful bacteria. At the same time, it hinders the re-colonization by *Lactobacillus spp*. Indeed, we found that both menses-related dysbiotic and unstable time series had more *Lactobacillus crispatus* and other non-*iners* species in their CST-III and CST-IV samples than the women classified as constant dysbiotic VCD.

We hypothesized that specific genomic strains of species might be more prone to promote either permanent dysbiosis or favour a stable *Lactobacillus spp*. dominance with a special focus on the prevalent and abundant genera *Gardnerella, Prevotella.* and *Lactobacillus.* The metagenomic assembly of the pangenomes revealed that despite all *Gardnerella* sequencing reads present being assigned to be *G. vaginalis,* this species was the least prevalent genomic *Gardnerella spp.*. Some studies have suggested that the production of bacteriocins by *Gardnerella spp.* may contribute to the pathogenesis of BV by inhibiting the growth of *Lactobacilli spp.* and other beneficial bacteria in the vaginal microbiome. *Gardnerella spp*. can produce antagonistic substances against eubiotic indicator strains in BV conditions, such as *Lactobacillus acidophilus* ATCC 3456. In this study, bacteriocin-like genes in *G. leopoldii*, such as lctA and lagD, affect Gram-positive bacteria cytoplasmic membranes and inhibit cell wall synthesis, resulting in a broad action spectrum in the vaginal niche^28,29^. As a result, this could create an environment conductive to the overgrowth of pathogenic bacteria and dysbiosis. Indeed, we identified three bacteriocins produced by *G. leopoldii,* which were almost exclusive to the VCDs constant dysbiotic and unstable, supporting a role for bacteriocins in maintaining dysbiosis.

The new time series classification presented here extends the complexity of vaginal dysbiosis and provides a better characterization of the vaginal ecology system. This framework may aid interdisciplinary translational teams working to improve reproductive outcomes. Further research is needed to identify markers of VCDs that could help reduce the need for consecutive sampling. Based on the data presented here, the clinicians need to sample two-three times mid-cyclical and two times during menses to be able to classify a dysbiotic sample into the accurate VCD.

## Conclusion

Women have been excluded from clinical trials primarily due to the complexity related to the menstrual cycle changes. This approach has left a black box in medicine that will require substantial focus to overcome. Indeed, this study confirms the complexity related to the cycle and based on the data presented here we propose four Vaginal Community Dynamics based on the specific dynamics of the bacterial composition over time. This categorization of the time series of vaginal samples enables comparisons at individual and population levels. It will assist in identifying the drivers behind the different dynamic profiles of the vaginal microbiome by gaining a better ecological understanding of the role of *Lactobacillus spp*. and their interaction with the host and other components of the vaginal microbiota. To further understand why some women are resilient to exposures such as menses and intercourse while others are not, there is a need for detailed research on the bacterial, fungal and viral populations of the four dynamic categories. These findings could develop into both prevention and rescue strategies against bacterial vaginosis. Future research should also assess whether transient dysbiosis, either menses-related or unstable, presents a risk profile similar to either constant eubiosis or dysbiosis.

## Methods

### Participants and samples

The participants of this study were recruited for the MiMens study, aiming to understand the interplay between hormonal contraceptives, the menstrual cycle and the human microbiome^30,31^. The samples analyzed in this study were collected at home by the participants daily for 42 days, starting on the first day of menses. Women without regular menses started their sampling on a random day. Samples were collected with a FLOQSwab (COPAN diagnostics) and preserved in DNA/RNA-shield (Zymo Research) in the participant’s house until the end of the 42 days when they were taken to the clinic and frozen at -80°C. For the first part of this paper, 15 participants were selected, five each on combined oral contraceptives (COC), levonogestrel-intrauterine system (LNG-IUS), or not using hormonal contraceptives (NHC), and all samples were analyzed. For the second part, 49 participants were selected from the MiMens cohort, 15-17 from each contraceptive group. The included participants had collected a minimum of 22 out of 25 samples from cycle day 4 and 28 days onwards (cycle day 32 or early in the next cycle).

### DNA extraction, library preparation and sequencing

DNA was extracted with Quick-DNA Magbead Plus kit, as previously described^33^. Libraries of 16S rRNA gene fragments were based on the V3-V4 region, using 80 ng of input DNA and primers 341f-805r^34^ prepared with a construct containing Illumina adapters and double barcodes: 341f 5’-CAA GCA GAA GAC GGC ATA CGA GAT N8 GTC TCG TGG GCT CGG AGA TGT GTA TAA GAG ACA GGA CTA CHV GGG TAT CTA ATC C-3’ and 805r 5’-AAT GAT ACG GCG ACC ACC GAG ATC N8 TCG TCG GCA GCG TCA GAT GTG TAT AAG AGA CAG CCT ACG GGN GGC WGC AG-3’, where N8 represents an eight bp long barcode. PCR was conducted for 25 cycles of 98°C for denaturation, 53°C for annealing, and 72°C for the extension. Shotgun metagenomic libraries were prepared with the MGI FS DNA library prep kit (MGI, Shenzhen, China) with the alterations described in^33^ and sequenced on a DNBSEQ-G400 sequencer (MGI) using the high-throughput sequencing set (PE150 1000016952; MGI) with DNA libraries loaded onto to the flow cell using the DNB loader MGIDL-200 (MGI).

### Taxonomic annotation

16S rRNA gene amplicons were processed and annotated with the DADA2 pipeline^35^ using standard settings. Taxonomy was based on the SILVA 128 database^36^.

Shotgun libraries were annotated by mapping to the OptiVag DB v2 with kraken2^37^, as previously described^33^. Viruses were annotated with kmcp^38^ against the genbank-viral database with standard parameters.

### Metagenomic assembly, binning, and annotation

Before assembly, metagenomic read libraries were normalized with bbnorm^39^ to discard reads with a coverage <3 and subsample those with a coverage >80. All available samples for each participant were co-assembled using Metaspades with standard parameters^40^. Reads were mapped back to contigs using bbmap^39^. Before mapping, contigs were filtered to retain those with >1kbp and contigs longer than 20 kbp were broken up into 10 kbp segments with a 100 bp overlap. The mapping and composition information was used for metagenomic binning using CONCOCT v1.1.0^41^. Proteins were called and annotated using Prokka^42^. Bins were then analyzed with checkm^43^ and retained if they presented >90% completeness and <2% contamination. Phylogenetic trees were built with FastTree with standard parameters^44^. Phylogenomic and pangenomic analyses were run in Panaroo^45^ with standard settings. Since Panaroo only accepts dichotomous variables, we classified both “constantly eubiotic" and “menses-related dysbiotic” as “mid-cycle eubiosis”, in contrast to the “constant dysbiosis” and “unstable” groups.

### Statistics and figures

All figures were generated in R v4.2.2. Alpha- and Beta-diversity statistics were calculated with Vegan v.2.6.4. Alpha-diversity was calculated as inverted Simpson’s, and beta-diversity as Aitchinson’s distance, unless specified as Jaccard’s. Associations of specific bacteria to the beta-diversity step size to the following sample were calculated in ANCOM-BC v2.0.1 and corrected through the Benjamini-Hochberg procedure^46^. CSTs were assigned with VALENCIA, github commit c41897d^47^. Time series were further classified into four VCDs (constantly eubiotic, menses-related dysbiotic, unstable and constantly dysbiotic) as described in the main text and in www.github.com/ctmrbio/valody. Differences in total phage content between groups were calculated with Kruskal-Wallis tests.

## Declarations

### Ethics approval and consent to participate

The study is approved by The Regional Ethics Committee on Health Research (H-17017580) and the Data Protection Agency in the Capital Region of Denmark (2012-58-0004). All data were collected and hosted using REDCap electronic data capture tools^32^ hosted by the Capital Region of Denmark.

### Consent for publication

All participants gave oral and written consent to participate according to the Helsinki Declaration and were remunerated with 3,000 DKK after completing sample collection.

### Availability of data and material

All sequencing data has been submitted to the European Nucleotide Archive under project PRJEB37731. The 16S samples have accession ID ERS14866734-ERS14867253, and the shotgun samples, after human DNA removal, have ID ERS14864440-ERS14865713. The code is available at https://github.com/ctmrbio/valody/commit/17cb300a4571819260daa54319473f8a5dc9161a.

## Funding

LWH has been partially supported by the SciLifeLab & Wallenberg Data Driven Life Science Program (grant: KAW 2020.0239). The Rigshospitalet Research Fund has supported MCK (E-22614-01 and E-22614-02). JD has been supported by the Swedish Research Council (grant: 2021-01683).

The Centre for Translational Microbiome Research is partly funded by Ferring Pharmaceuticals (LWH, EF, JD, LE, IS-K). An unrestricted research grant from Ferring Pharmaceuticals enabled the clinical infrastructure and sampling (MCK, ZB and HSN). The funder was not involved in the study design, collection, analysis, interpretation of data, the writing of this article or the decision to submit it for publication.

### Author contributions

MCK, HSN – planned and organized study cohort, obtained ethics and data protection approval, collected samples, sample and data management, analysed data and wrote manuscript

ZB – included participants and secured informed consent, collected samples, sample and data management, wrote manuscript

LWH, KV, JD, VK - analysed data, wrote the manuscript

EF – wrote the manuscript

ISK, LE – planned and organized the study cohort, analysed data, wrote the manuscript

All authors have read and approved the final manuscript

## Acknowledgements

We would like to thank the nurses employed at the Recurrent Pregnancy Loss Unit in Copenhagen for their tremendous contribution with including participants: Louise Lunøe, Karen Kirchheiner and Marie Chonovitsch. We appreciate Alexandra Pennhag, Marica Hamsten and Maike Seiferts efforts in DNA extraction and sequencing. We further acknowledge Fredrik Boulund’s continuous support in improving the quality of the Valody code base.

## Supplementary figure and table legends

**Supplementary figure 1: Bacterial and viral profiles for each sample over one menstrual cycle**

Each subject’s bacterial and viral profile are depicted as area plots. Sexual intercourse is overlaid as blue dots and vaginal bleedings as red dots. Log10 of the ratio of viral to bacterial reads is shown as a black line over the viral profiles, for time-series with sufficient data (>5 samples with detectable phages). Missing data is omitted.

Next to each taxonomic profile is an ordination showing all samples in the study as gray circles, and the samples for the relevant subject as numbers, following the days of their menstrual cycle. Days with vaginal bleedings are shown in red, days with sexual intercourse in blue and days with both events in purple.

**Supplementary figure 2: CST distribution and time-series dynamics for the 16S samples**

CSTs are shown as colored dots as per the legend in the second part. The outline of each box depicts the assignment to vaginal community dynamics. Missing samples are omitted. Bleedings are marked as red dots. Blue: constant eubiotic. Green: menses-related dysbiotic. Yellow: unstable. Red: constant dysbiotic.

**Supplementary figure 3: CST distribution and time-series dynamics for the shotgun samples**

CSTs are marked as colored dots above the taxonomic profiles as per the legend. The outline of each box depicts its dynamic group. Bleedings are marked as light red dots. Missing samples are omitted. Blue: constant eubiotic. Green: menses-related dysbiotic. Yellow: Unstable. Red: constant dysbiotic.

**Supplementary figure 4: Log-fold change of bacterial species in samples from CST-I**

Samples in CST-IA and CST-IB from menses-related dysbiotic or unstable individuals were compared to constant eubiotic individuals. The heatmap shows the log-fold change of all significant differences. Gray fields represent no significant change.

**Supplementary figure 5: Log-fold change of bacterial species in samples from CST-III**

Samples in CST-IIIA and CST-IIIB from menses-related dysbiotic or unstable individuals were compared with constant dysbiotic individuals. The heatmap shows the log-fold change of all significant differences. White fields represent no significant change.

**Supplementary figure 6: Volcano plots for the vaginal community dynamics compared to either constant eubiotic or constant dysbiotic**

**Supplementary figure 7: Histograms showing gene cluster prevalence in nine relevant pangenomes**

For each species, the prevalence (number of genomes containing each gene cluster) of each gene cluster is shown as a histogram. Gene clusters present in most or all genomes are considered “core”, while those in one or very few genomes can be considered “cloud”. The “shell” genomes, present in many, but not all genomes, are less frequent in this dataset.

**Supplementary figure 8: Phylogenomic analysis of *Lactobacillus* genomes**

Phylogenomic analysis of all detected *Lactobacillus* species does not find a correlation between the womens’ vaginal community dynamics and the observed phylogeny. The presence of a gene is represented in dark blue and its absence in light blue. Blue: constant eubiotic. Red: constant dysbiotic. Yellow: unstable. Green: menses-related dysbiotic.

**Supplementary figure 9: Phylogenomic analysis of *Prevotella* genomes**

Phylogenomic analysis of all detected *Prevotella* species does not find a correlation between the womens’ vaginal community dynamics and the observed phylogeny. The presence of a gene is represented in dark blue and its absence in light blue. Blue: constant eubiotic. Red: constant dysbiotic. Yellow: unstable. Green: menses-related dysbiotic.

**Supplementary table 1:** Full ASV table for samples sequenced by 16S marker gene sequencing. Each sample is in a column, named by individual ID and cycle day, and each ASV in a row. Taxonomic annotations are in the second-to-last column and centroid sequence in the last

**Supplementary table 2:** Full taxonomic annotation and feature counts for the samples sequenced by shotgun. Each sample is in a column, named by subject and cycle day, and each taxon in a row

**Supplementary table 3:** Differential abundance results for samples in CST-IA from individuals with menses-related dysbiotic or unstable VCD compared to constant eubiotic VCD

**Supplementary table 4:** Differential abundance results for samples in CST-IB from individuals with menses-related dysbiotic or unstable VCD compared to constant eubiotic VCD

**Supplementary table 5:** Differential abundance results for samples in CST-IIIA from individuals with menses-related dysbiotic or unstable VCD compared to constant dysbiotic VCD

**Supplementary table 6:** Differential abundance results for samples in CST-IIIB from individuals with menses-related dysbiotic or unstable VCD compared to constant dysbiotic VCD

**Supplementary table 7:** Differential abundance results for all samples in menses-related dysbiotic, unstable and constant dysbiotic VCD against constant eubiotic

**Supplementary table 8:** Differential abundance results for all samples in menses-related dysbiotic, unstable and constant eubiotic VCD against constant dysbiotic

**Supplementary table 9:** Differential frequency of gene clusters in *Lactobacillus spp.*, contrasting constant eubiotic and menses-related dysbiotic vs. unstable and constant dysbiotic

**Supplementary table 10:** Differential frequency of gene clusters in *Gardnerella spp.*, contrasting constant eubiotic and menses-related dysbiotic vs. unstable and constant dysbiotic

**Supplementary table 11:** Differential frequency of gene clusters in *Prevotella spp.*, contrasting constant eubiotic and menses-related eubiotic vs. unstable and constant dysbiotic

