## Supplementary figures 1-9 for "Defining Vaginal Community Dynamics: daily microbiome transitions, the role of menstruation, bacteriophages and bacterial genes"

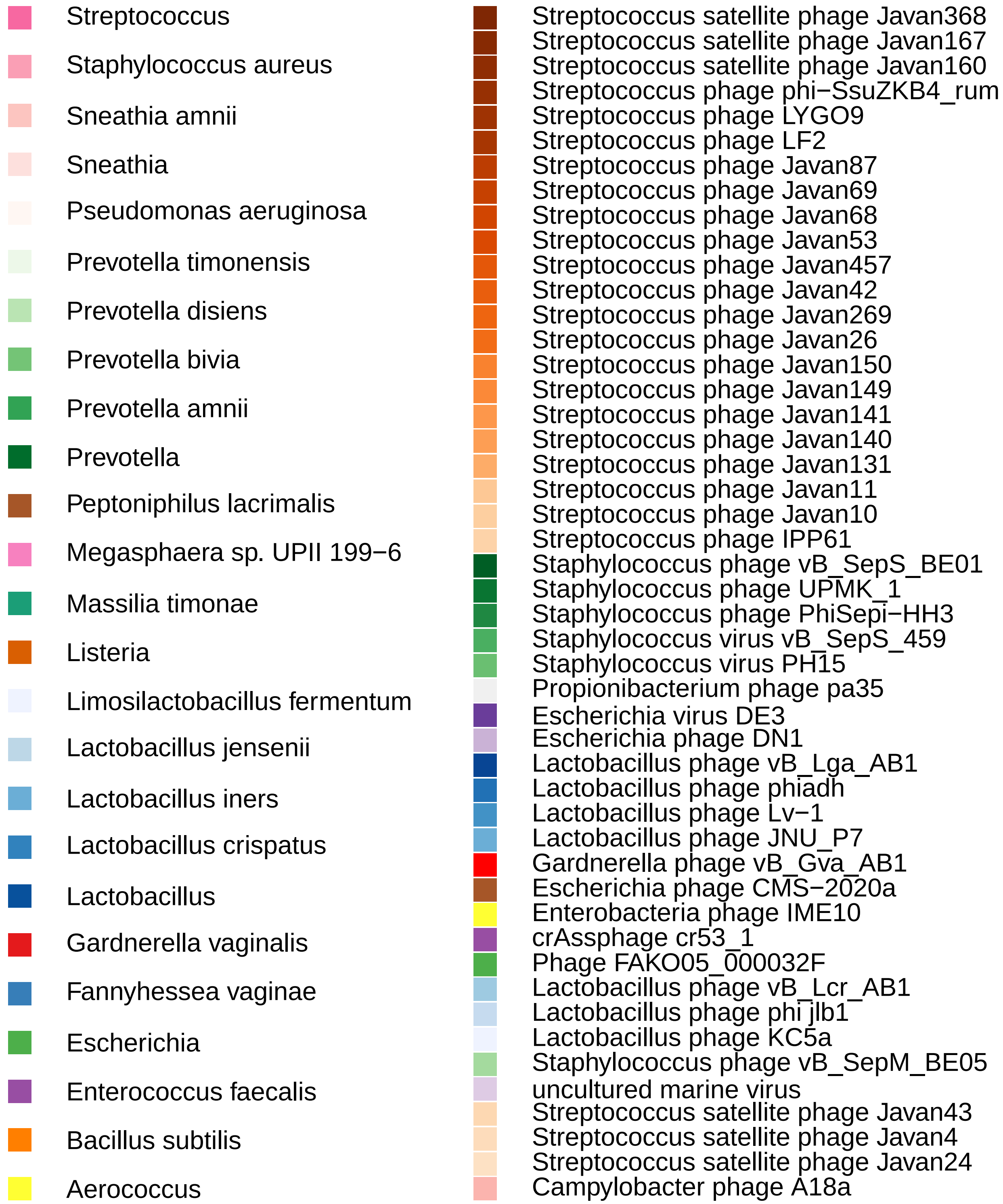
Streptococcus
Staphylococcus aureus
Sneathia amnii
Sneathia
Pseudomonas aeruginosa
Prevotella timonensis
Prevotella disiens
Prevotella bivia
Prevotella amnii
Prevotella
Peptoniphilus lacrimalis
Megasphaera sp. UPII 199−6
Massilia timonae
Listeria
Limosilactobacillus fermentum
Lactobacillus jensenii
Lactobacillus iners
Lactobacillus crispatus
Lactobacillus
Gardnerella vaginalis
Fannyhessea vaginae
Escherichia
Enterococcus faecalis
Bacillus subtilis
Aerococcus
Streptococcus satellite phage Javan368
Streptococcus satellite phage Javan167
Streptococcus satellite phage Javan160
Streptococcus phage phi−SsuZKB4_rum
Streptococcus phage LYGO9
Streptococcus phage LF2
Streptococcus phage Javan87
Streptococcus phage Javan69
Streptococcus phage Javan68
Streptococcus phage Javan53
Streptococcus phage Javan457
Streptococcus phage Javan42
Streptococcus phage Javan269
Streptococcus phage Javan26
Streptococcus phage Javan150
Streptococcus phage Javan149
Streptococcus phage Javan141
Streptococcus phage Javan140
Streptococcus phage Javan131
Streptococcus phage Javan11
Streptococcus phage Javan10
Streptococcus phage IPP61
Staphylococcus phage vB_SepS_BE01
Staphylococcus phage UPMK_1
Staphylococcus phage PhiSepi−HH3
Staphylococcus virus vB_SepS_459
Staphylococcus virus PH15
Propionibacterium phage pa35
Escherichia virus DE3
Escherichia phage DN1
Lactobacillus phage vB_Lga_AB1
Lactobacillus phage phiadh
Lactobacillus phage Lv−1
Lactobacillus phage JNU_P7
Gardnerella phage vB_Gva_AB1
Escherichia phage CMS−2020a
Enterobacteria phage IME10
crAssphage cr53_1
Phage FAKO05_000032F
Lactobacillus phage vB_Lcr_AB1
Lactobacillus phage phi jlb1
Lactobacillus phage KC5a
Staphylococcus phage vB_SepM_BE05
uncultured marine virus
Streptococcus satellite phage Javan43
Streptococcus satellite phage Javan4
Streptococcus satellite phage Javan24
Campylobacter phage A18a

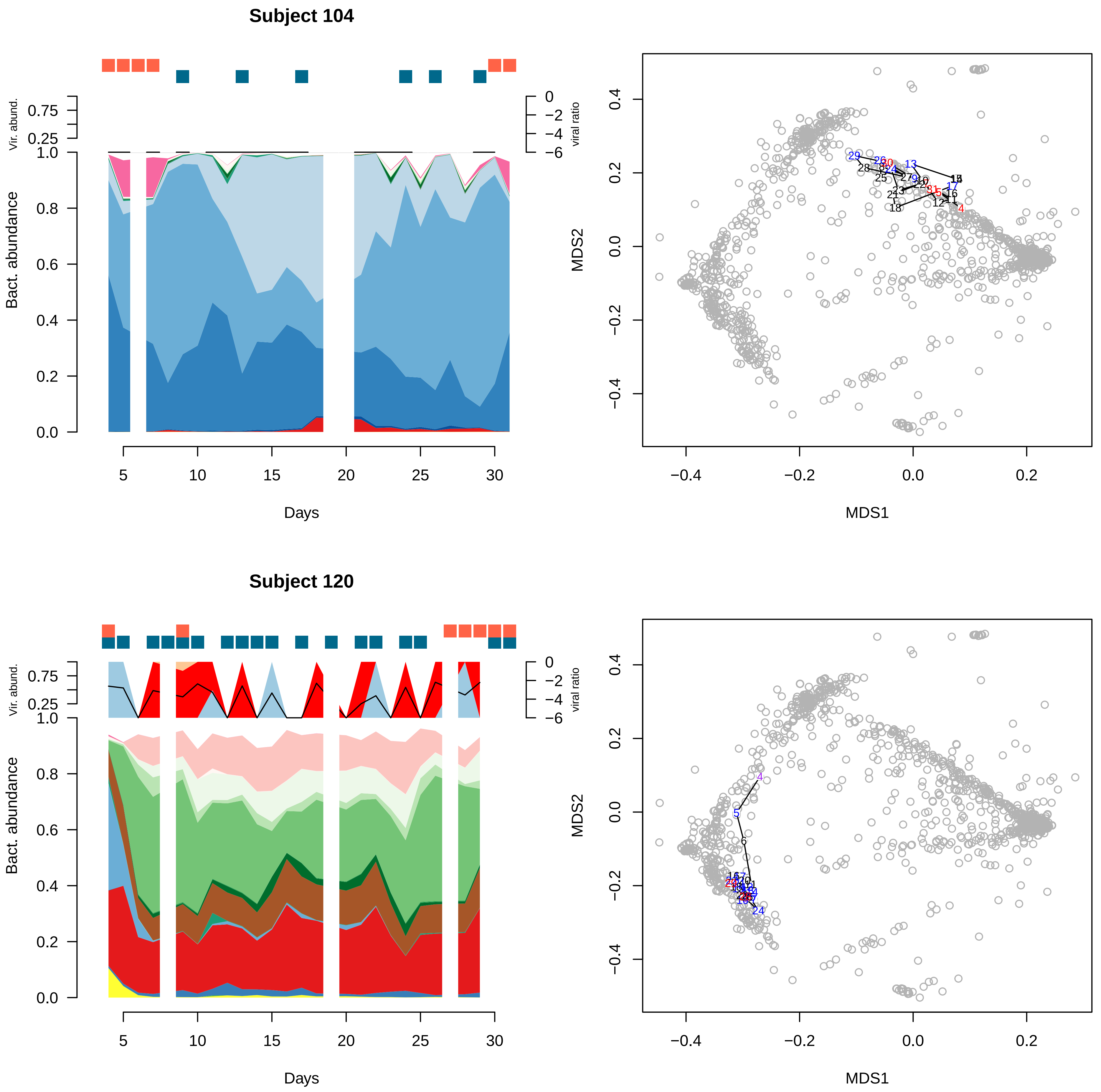
Subject 104
Vir. abund.
viral ratio
Bact. abundance
Days
MDS1
MDS2
Subject 120
Vir. abund.
viral ratio
Bact. abundance
Days
MDS1
MDS2

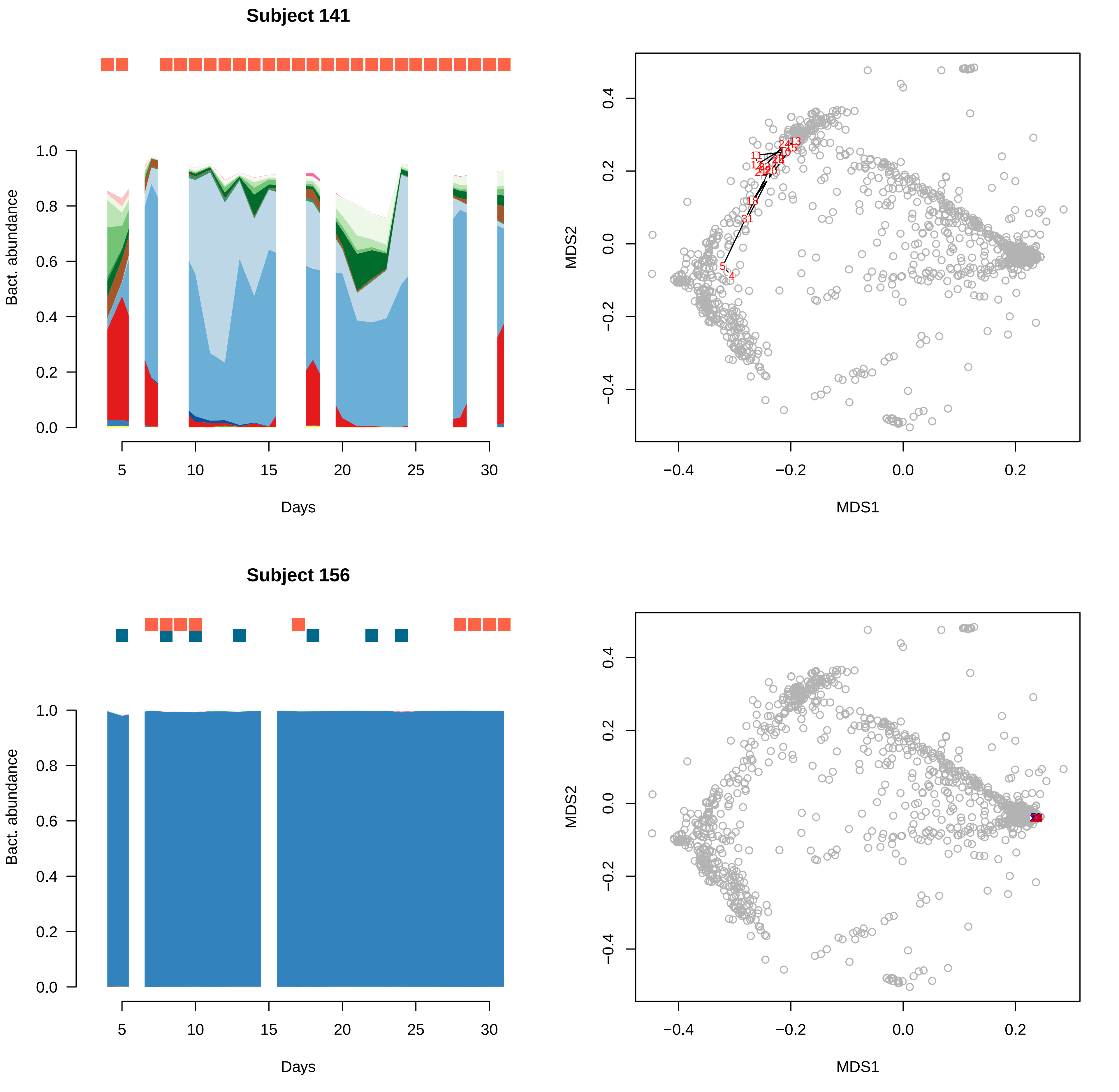

Subject 141
Bact. abundance
Days
MDS1
MDS2
Subject 156
Bact. abundance
Days
MDS1
MDS2

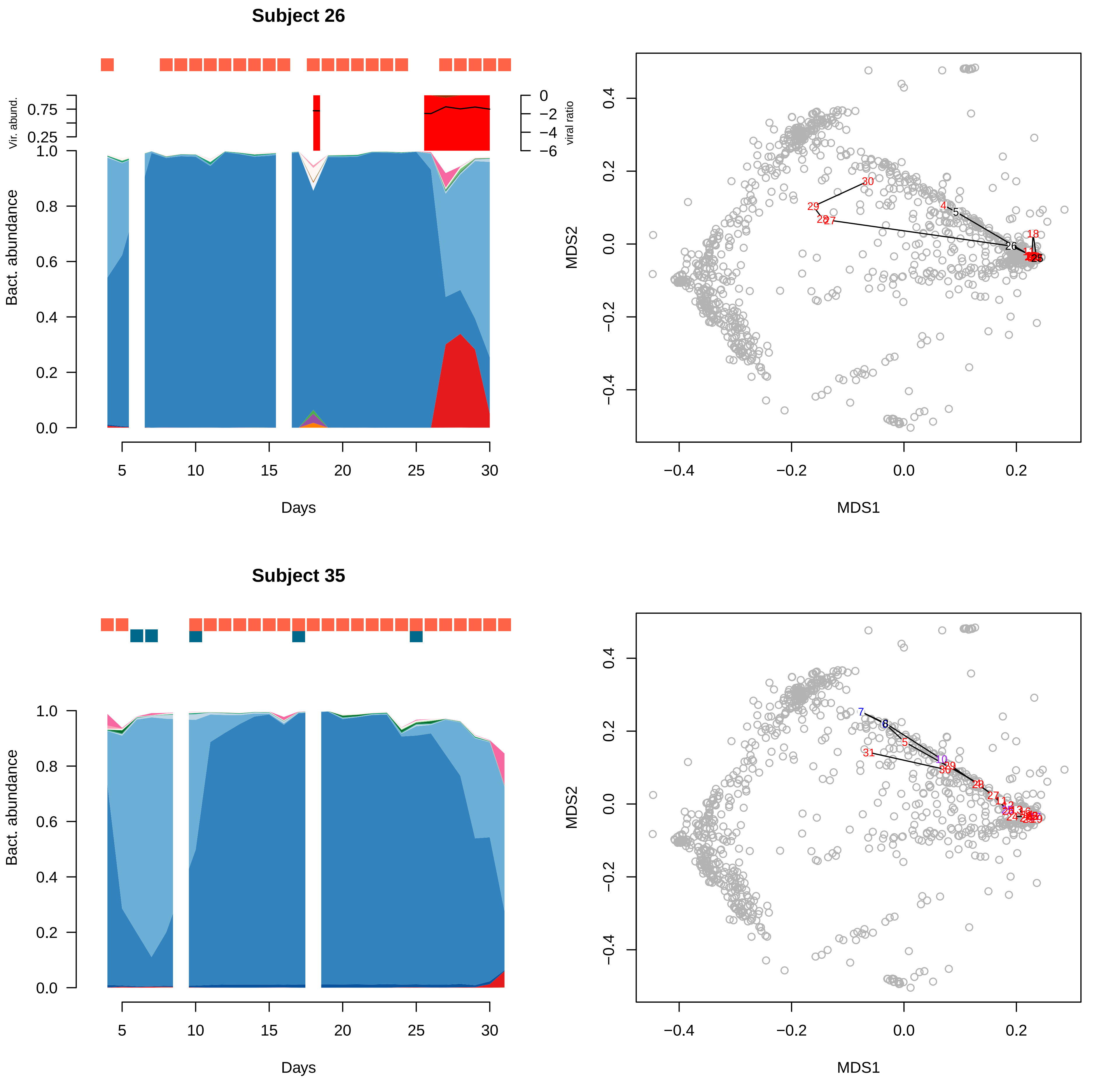

Subject 26
Vir. abund.
0.75
0.25
viral ratio
0
−2
−4
−6
Bact. abundance
1.0
0.8
0.6
0.4
0.2
0.0
5
10
15
20
25
30
Days
MDS2
MDS1
0.4
0.2
0.0
−0.2
−0.4
Subject 35
Bact. abundance
Days
MDS2
MDS1

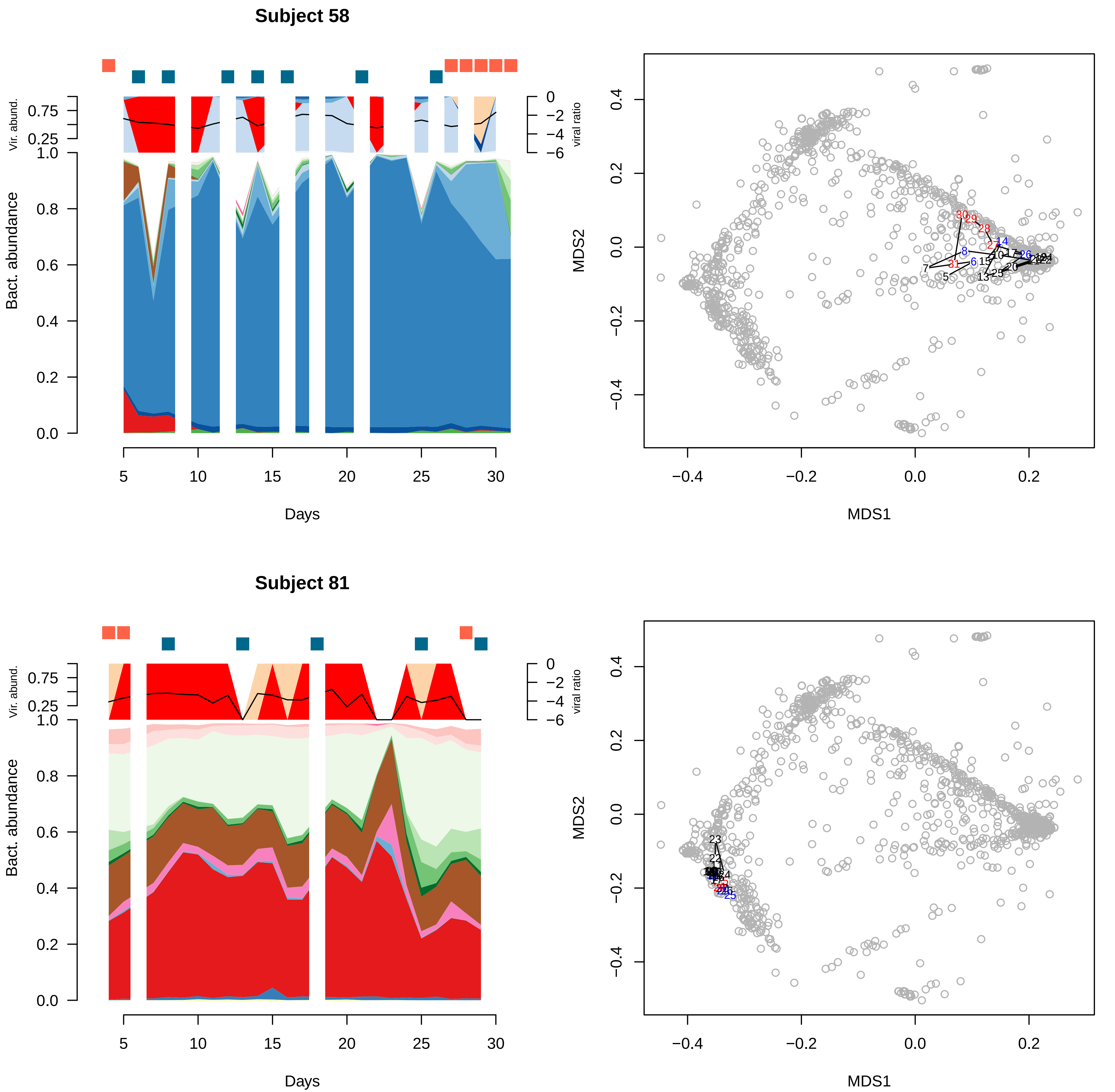
Subject 58
Vir. abund.
Bact. abundance
viral ratio
Days
MDS1
MDS2
Subject 81
Vir. abund.
Bact. abundance
viral ratio
Days
MDS1
MDS2

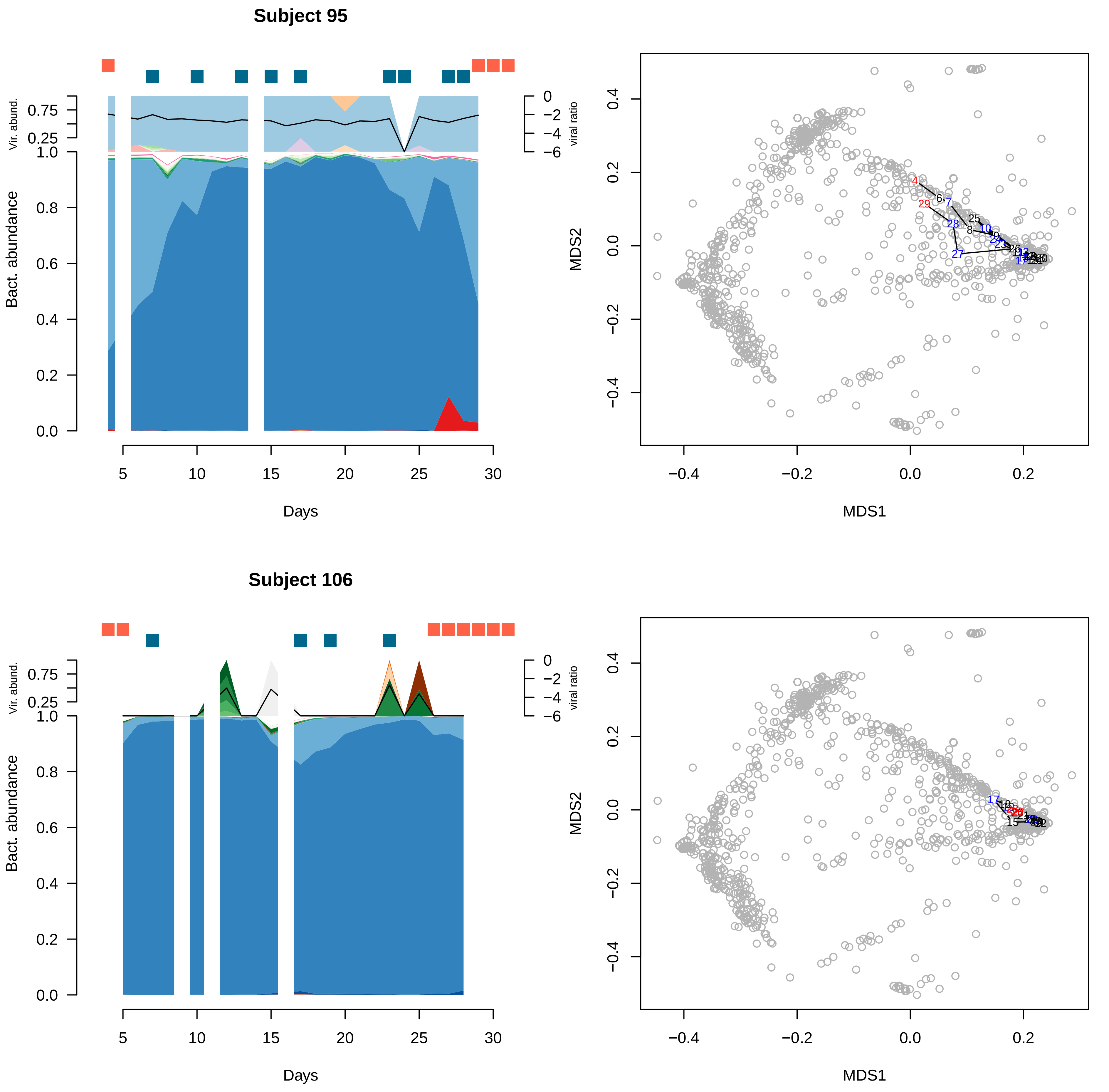

Subject 95
Vir. abund.
0.75
0.25
1.0
Bact. abundance
0.8
0.6
0.4
0.2
0.0
0
−2
−4
−6
viral ratio
5
10
15
20
25
30
Days
MDS2
MDS1
0.4
0.2
0.0
−0.2
−0.4
Subject 106

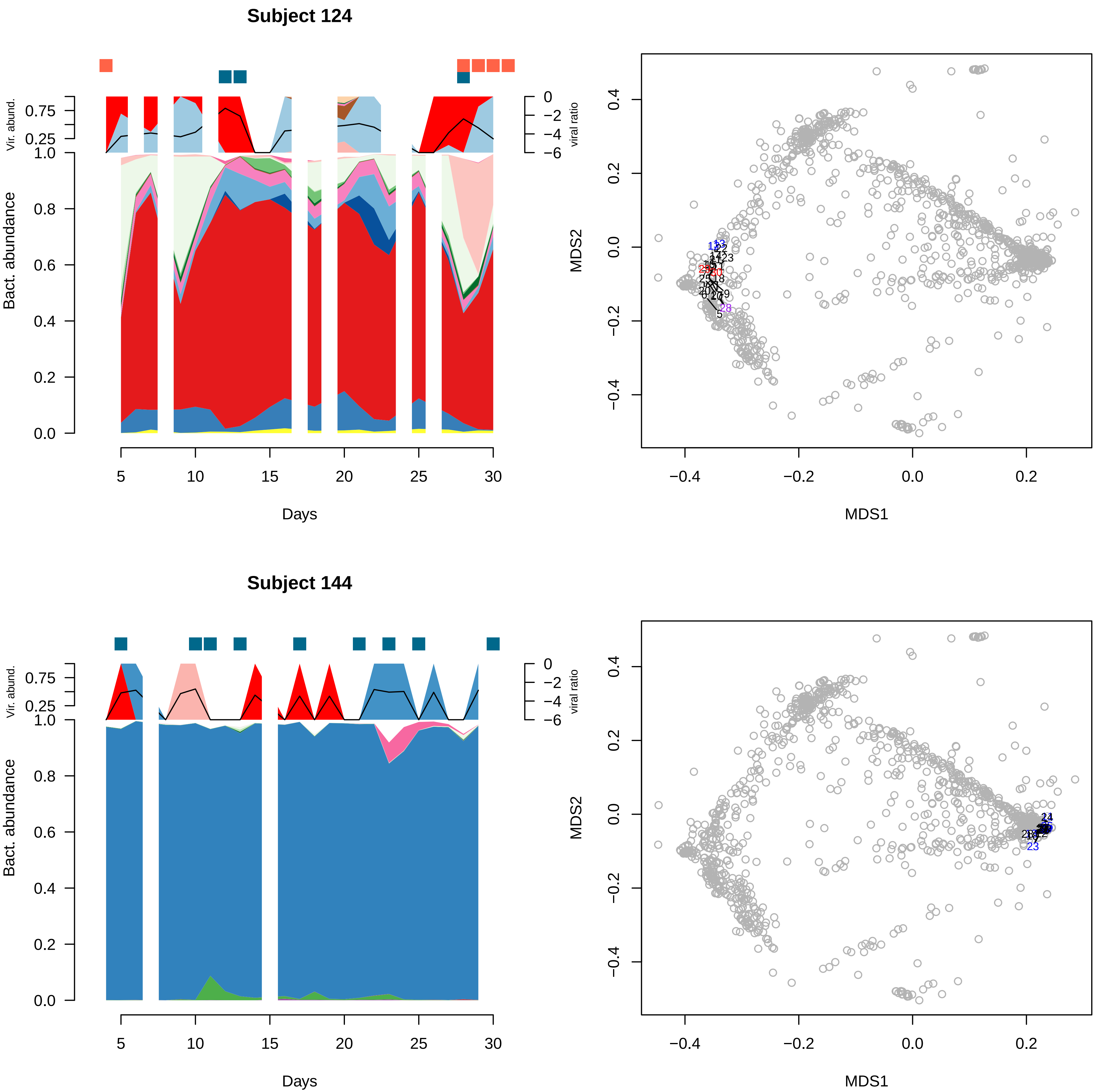

Subject 124
Vir. abund.
Bact. abundance
viral ratio
Days
MDS1
MDS2
Subject 144
Vir. abund.
Bact. abundance
viral ratio
Days
MDS1
MDS2

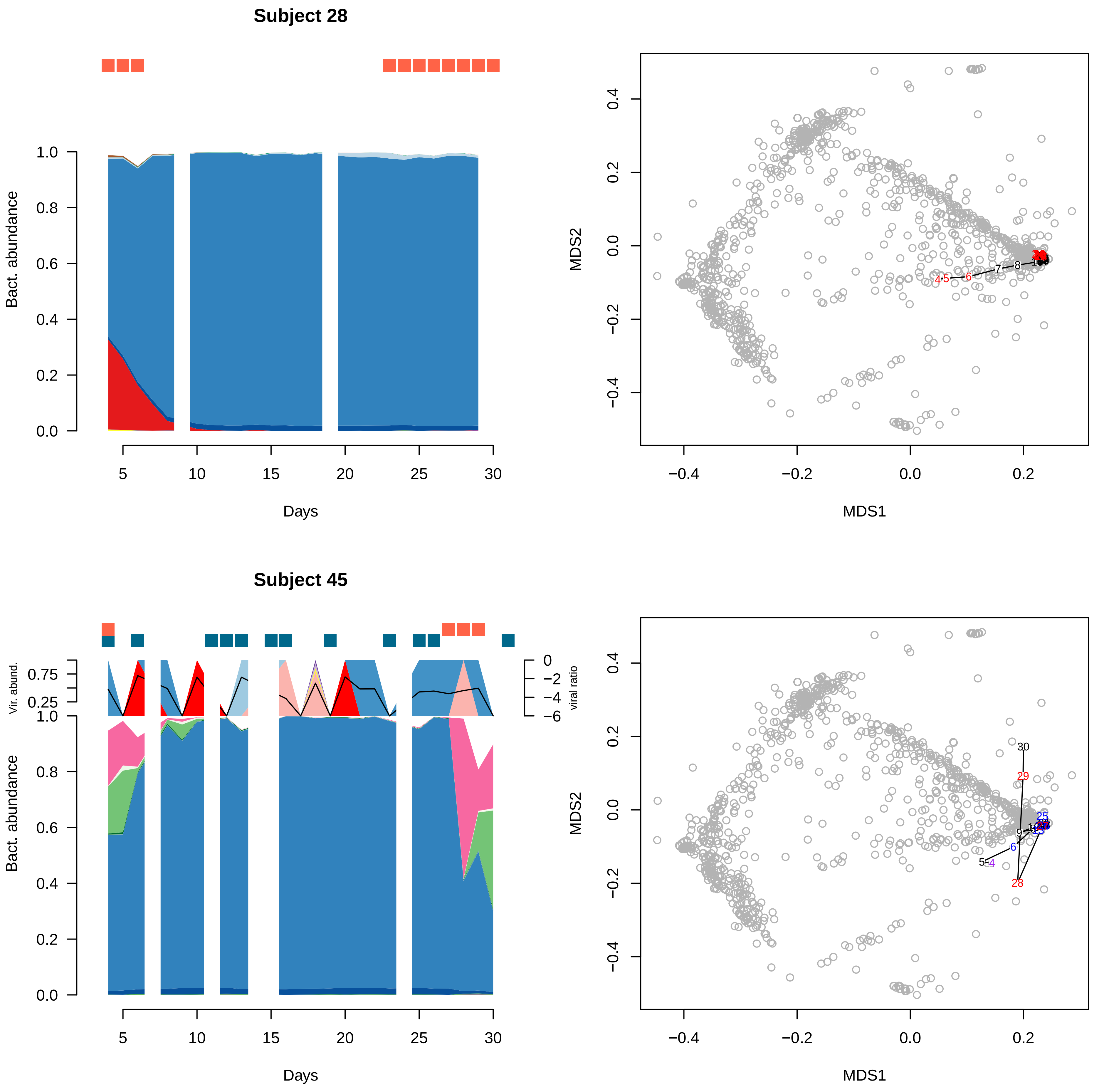

Subject 28
Bact. abundance
Days
MDS1
MDS2
Subject 45
Vir. abund.
viral ratio
Bact. abundance
Days
MDS1
MDS2

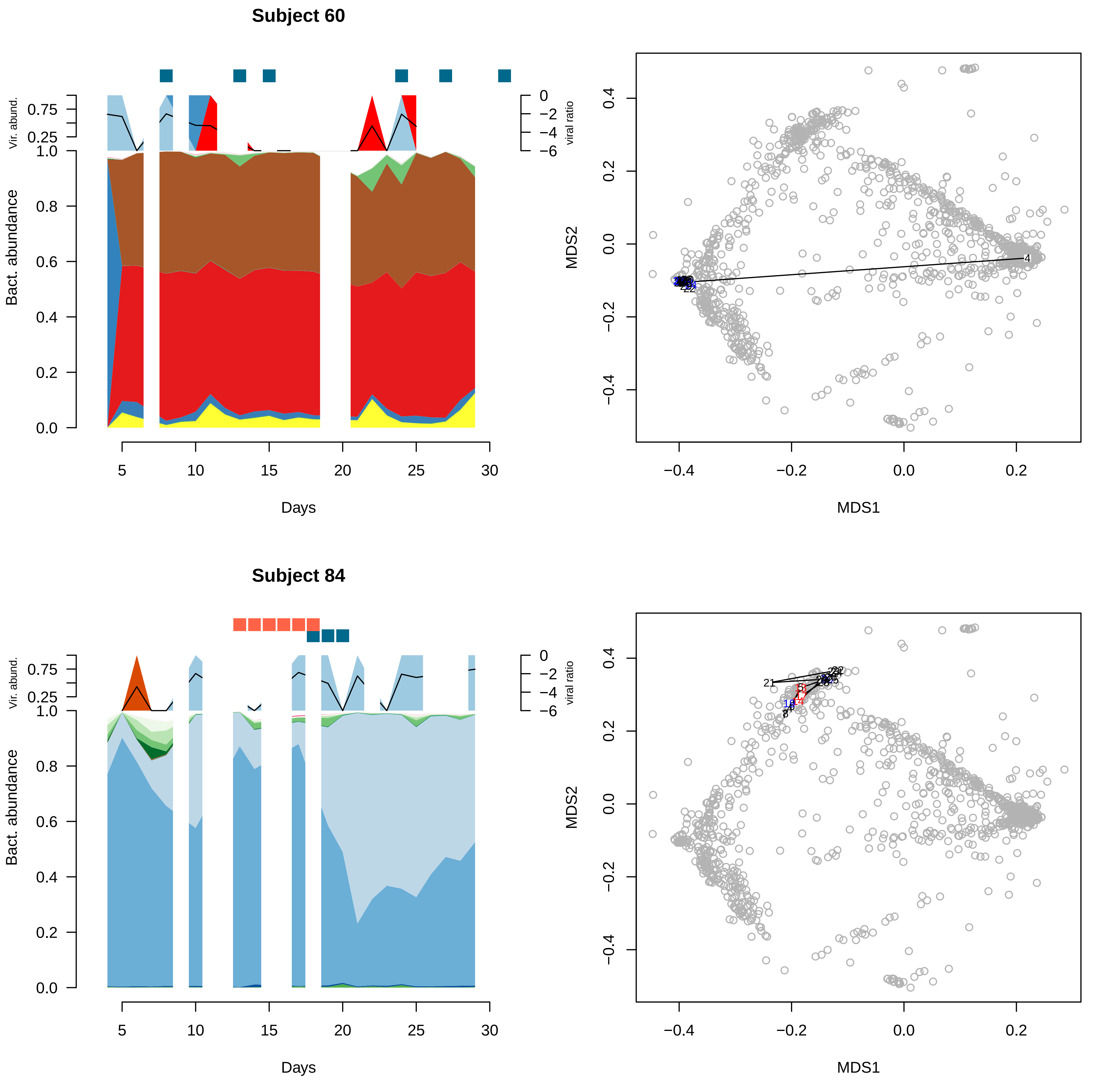
Subject 60
Vir. abund.
0.75
0.25
viral ratio
0
−2
−4
−6
Bact. abundance
1.0
0.8
0.6
0.4
0.2
0.0
5
10
15
20
25
30
Days
MDS2
0.4
0.2
0.0
−0.2
−0.4
MDS1
−0.4
−0.2
0.0
0.2
Subject 84
Vir. abund.
0.75
0.25
viral ratio
0
−2
−4
−6
Bact. abundance
1.0
0.8
0.6
0.4
0.2
0.0
5
10
15
20
25
30
Days
MDS2
0.4
0.2
0.0
−0.2
−0.4
MDS1
−0.4
−0.2
0.0
0.2

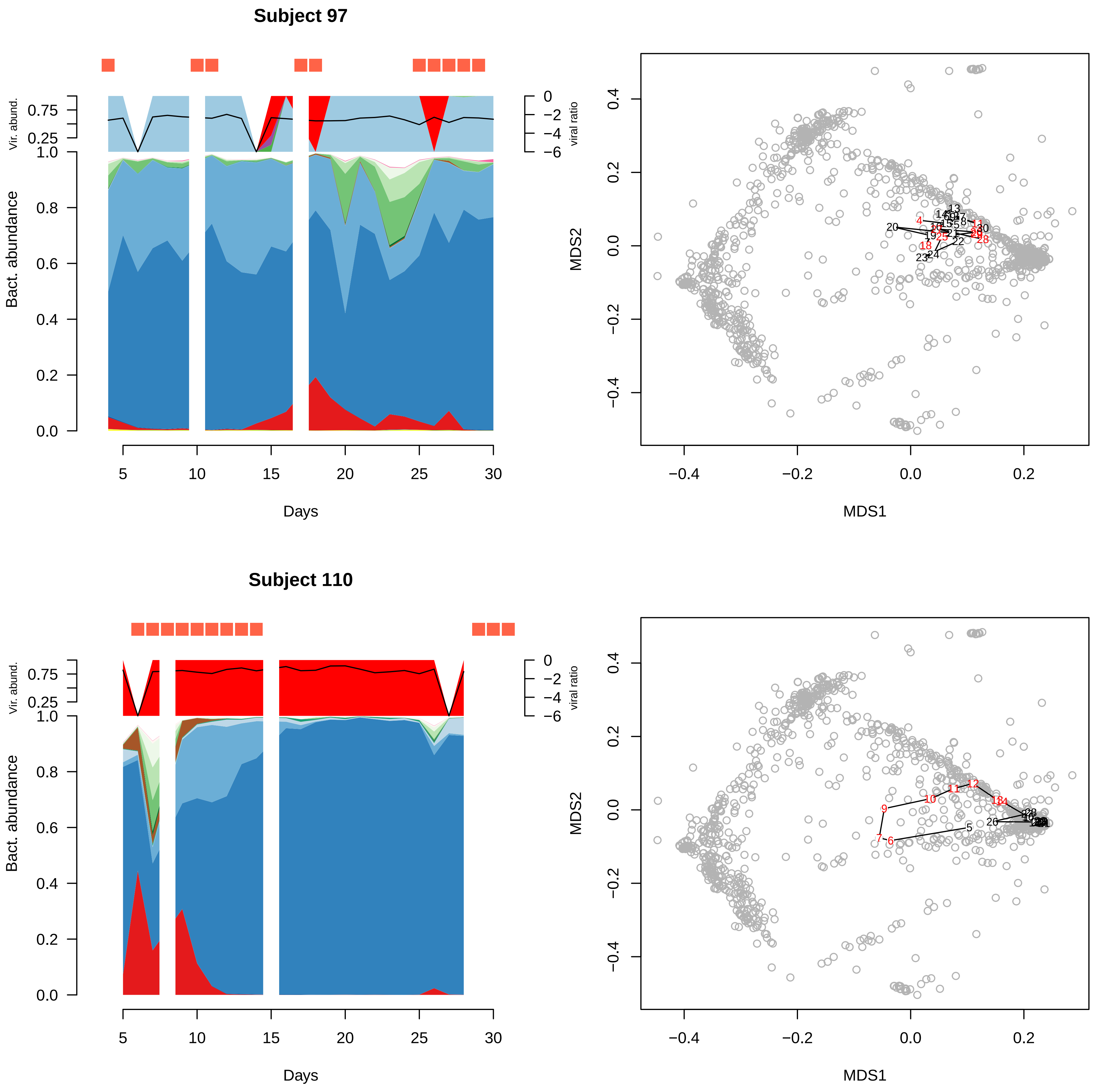
Subject 97
Vir. abund.
viral ratio
Bact. abundance
Days
MDS1
MDS2
Subject 110
Vir. abund.
viral ratio
Bact. abundance
Days
MDS1
MDS2

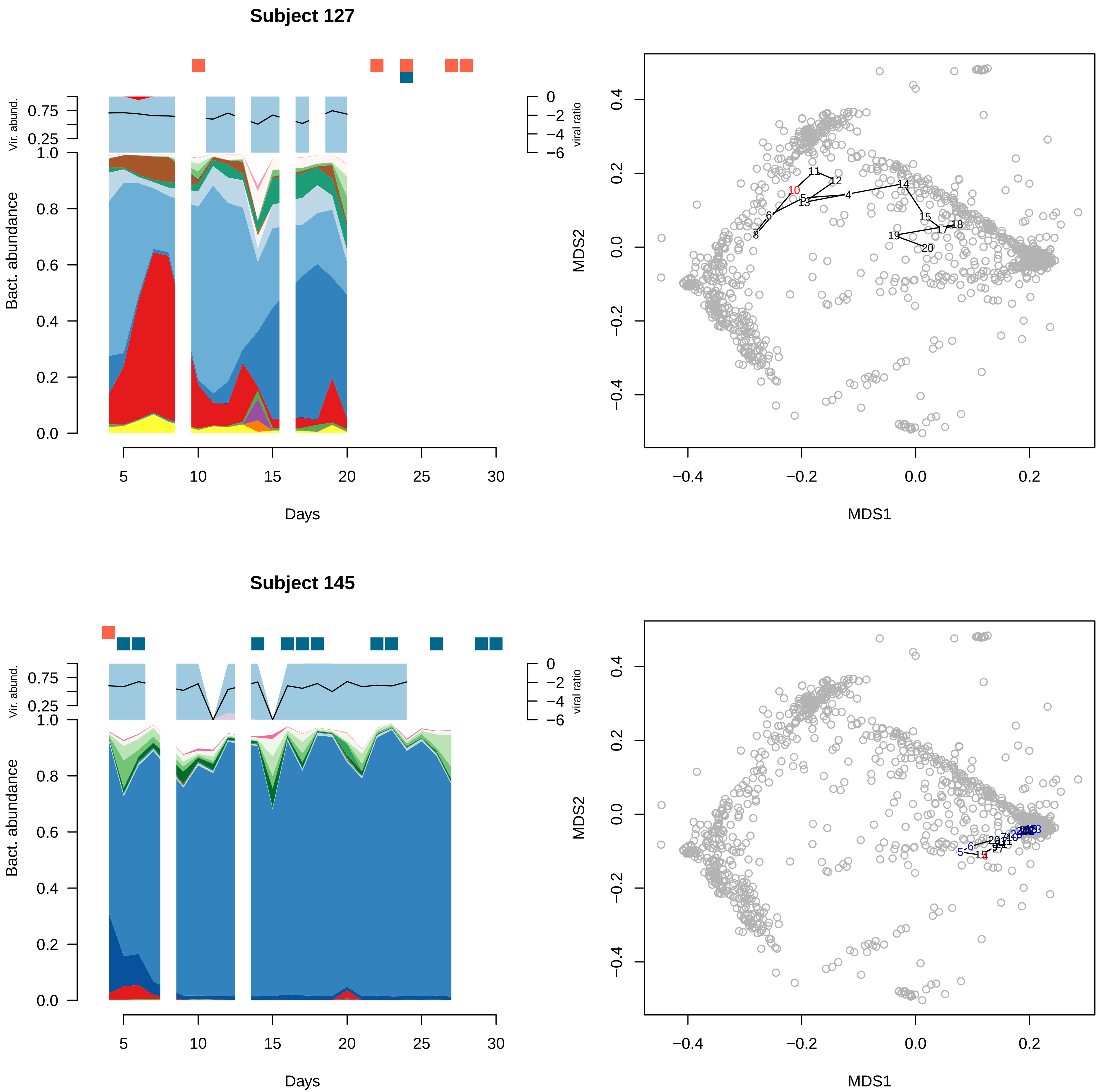

Subject 127
Vir. abund.
Bact. abundance
viral ratio
Days
MDS1
MDS2
Subject 145
Vir. abund.
Bact. abundance
viral ratio
Days
MDS1
MDS2

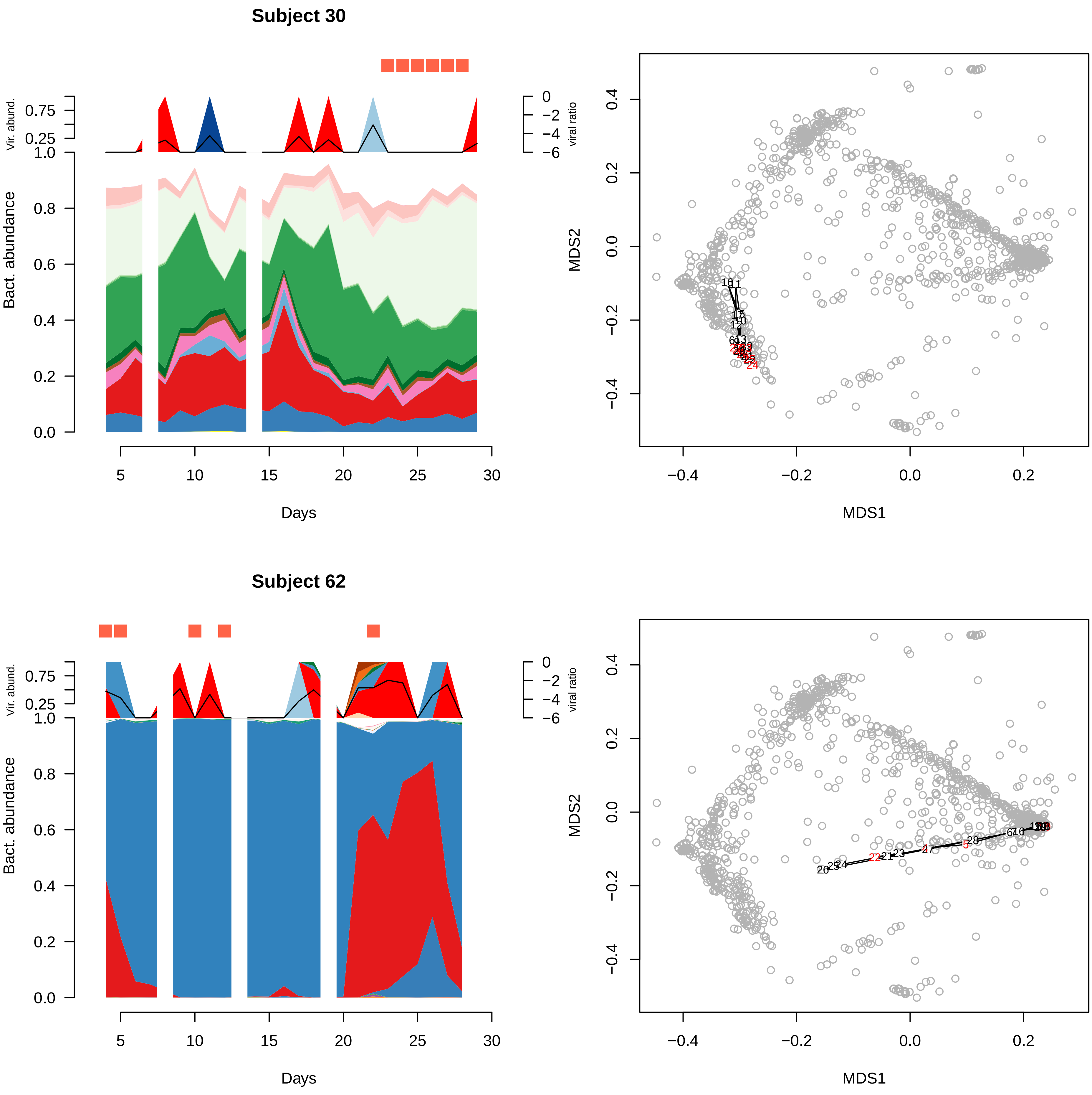

Subject 30
Vir. abund.
0.75
0.25
viral ratio
0
−2
−4
−6
Bact. abundance
1.0
0.8
0.6
0.4
0.2
0.0
5
10
15
20
25
30
Days
MDS2
MDS1
0.4
0.2
0.0
−0.2
−0.4
0.2
Subject 62
Vir. abund.
0.75
0.25
viral ratio
0
−2
−4
−6
Bact. abundance
1.0
0.8
0.6
0.4
0.2
0.0
5
10
15
20
25
30
Days
MDS2
MDS1
0.4
0.2
0.0
−0.2
−0.4
0.2

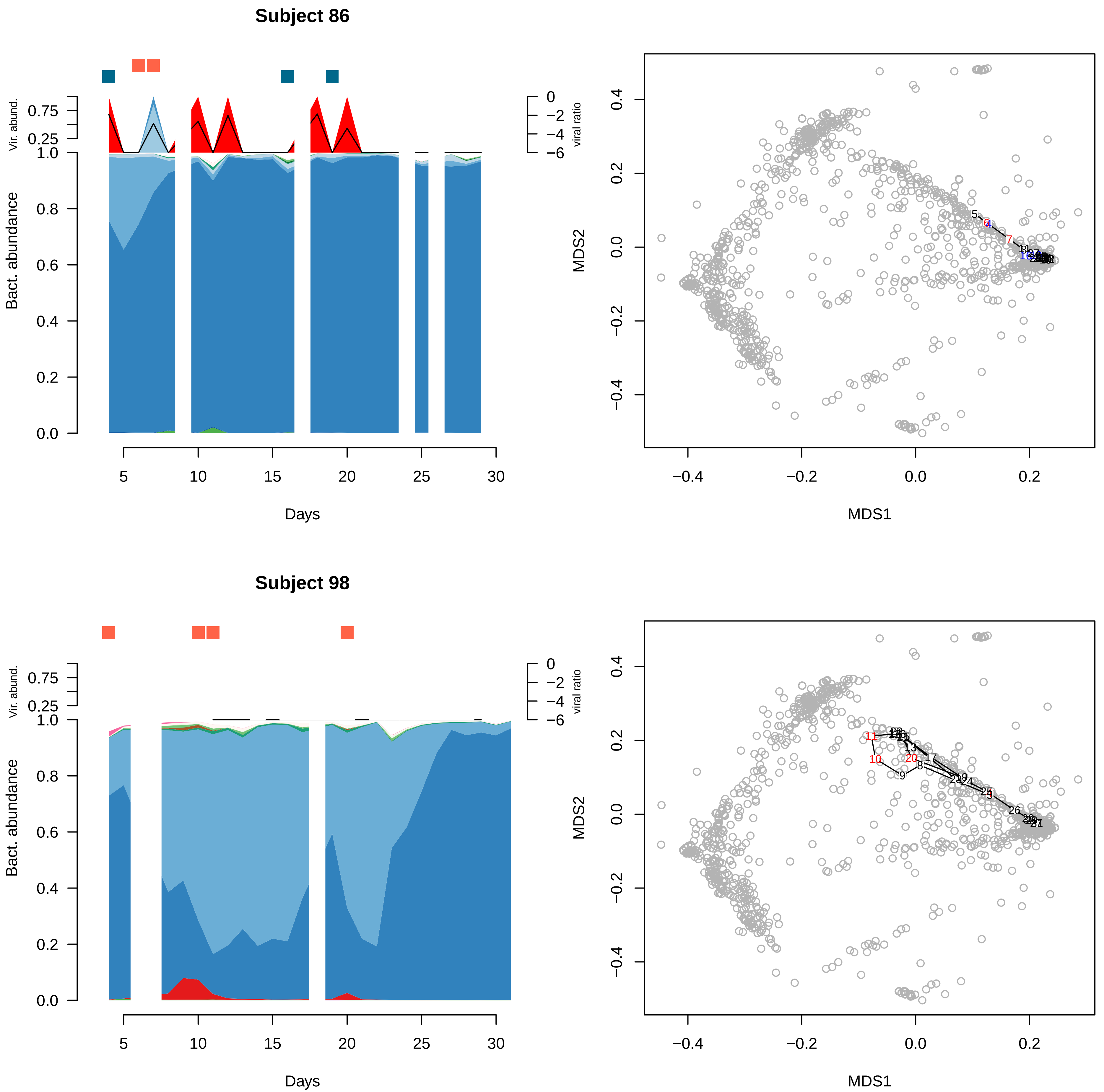

Subject 86
Vir. abund.
0.75
0.25
1.0
0.8
0.6
0.4
0.2
0.0
Bact. abundance
viral ratio
0
−2
−4
−6
5
10
15
20
25
30
Days
MDS1
MDS2
−0.4
−0.2
0.0
0.2
0.4
Subject 98
Vir. abund.
Bact. abundance
viral ratio
Days
MDS1
MDS2

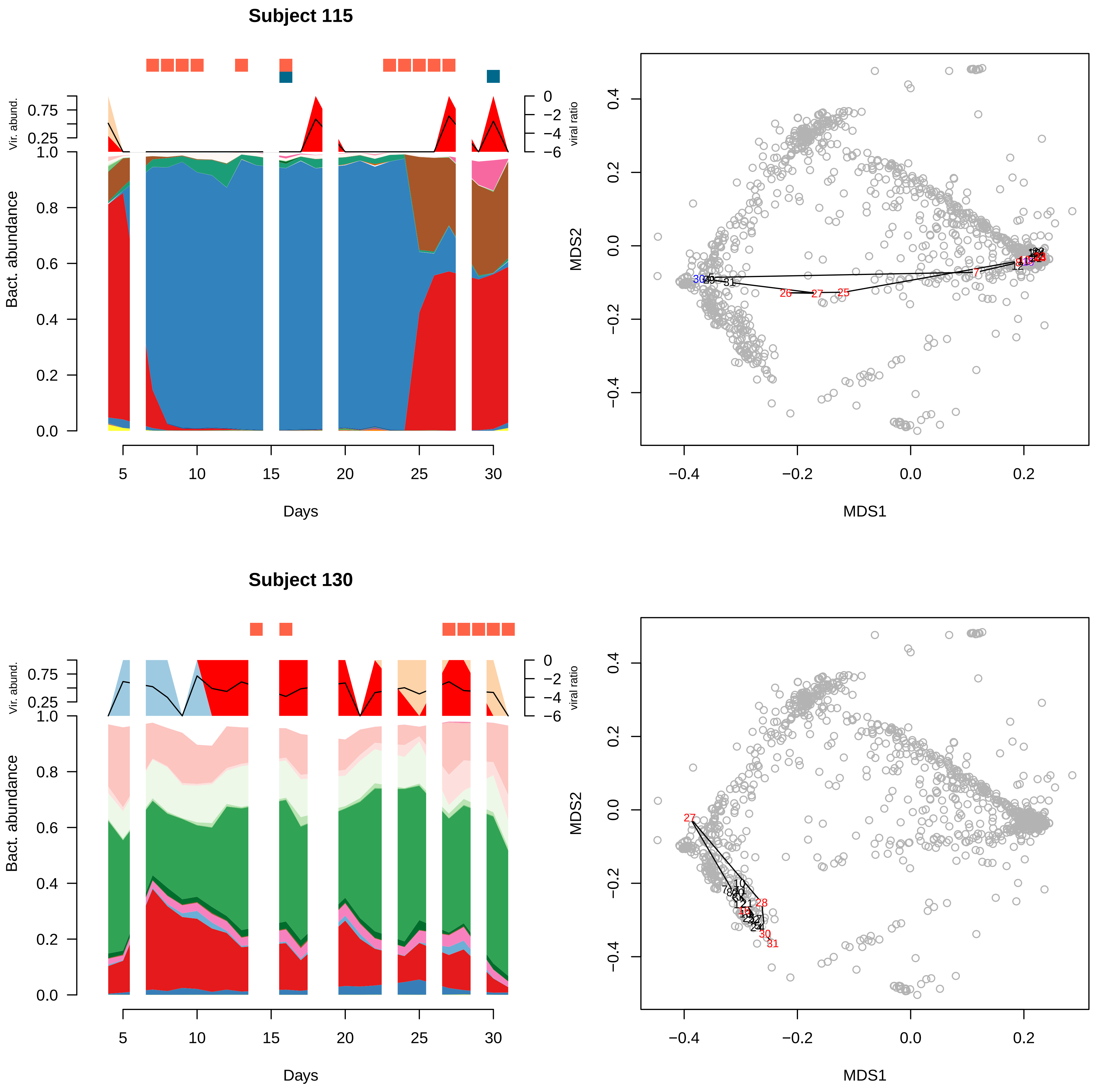

Subject 115
Vir. abund.
0.75
0.25
1.0
Bact. abundance
0.8
0.6
0.4
0.2
0.0
viral ratio
0
−2
−4
−6
5
10
15
20
25
30
Days
MDS2
MDS1
0.4
0.2
0.0
−0.2
−0.4
Subject 130

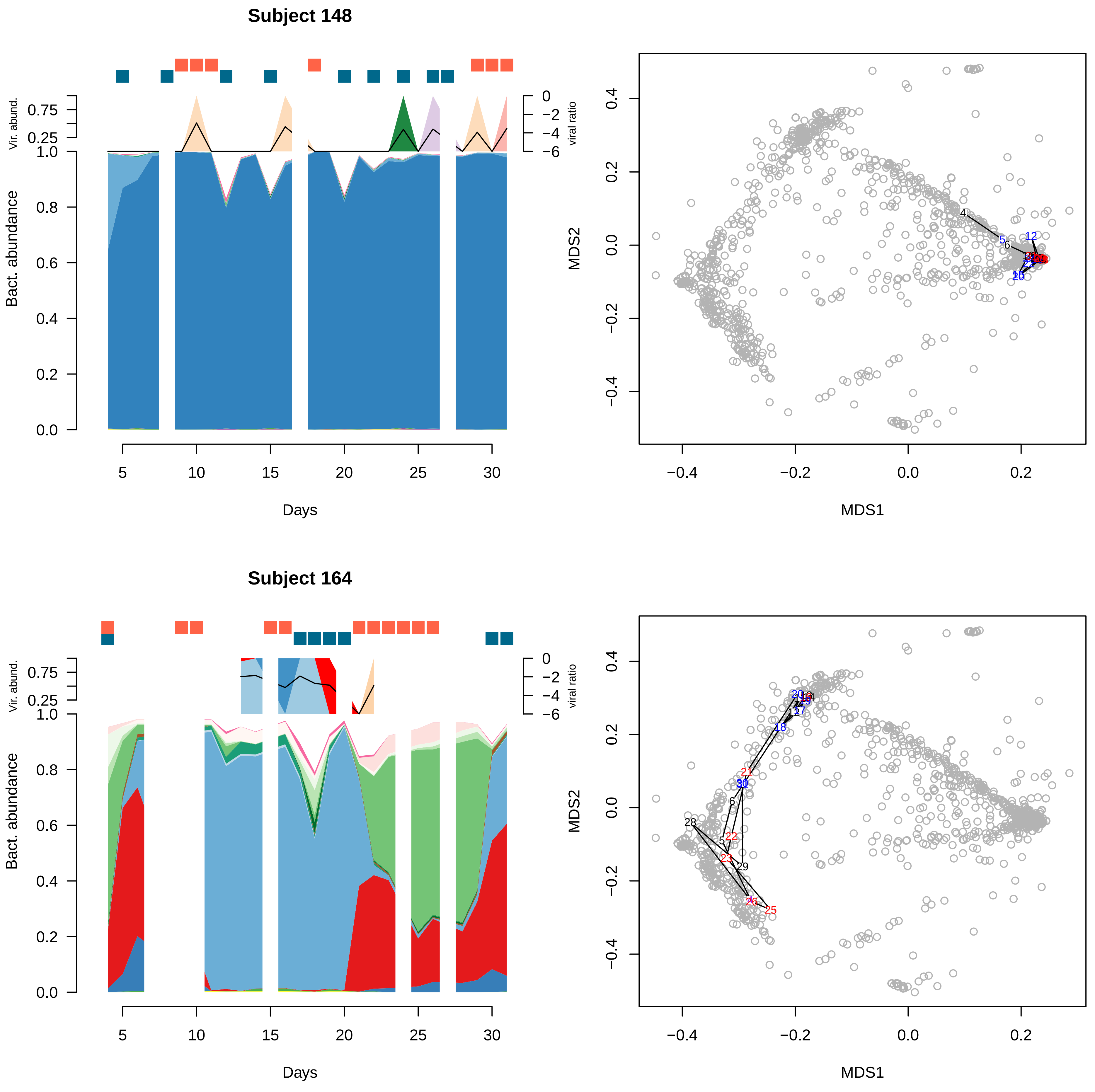
Subject 148
Vir. abund.
viral ratio
Bact. abundance
Days
MDS1
MDS2
Subject 164
Vir. abund.
viral ratio
Bact. abundance
Days
MDS1
MDS2

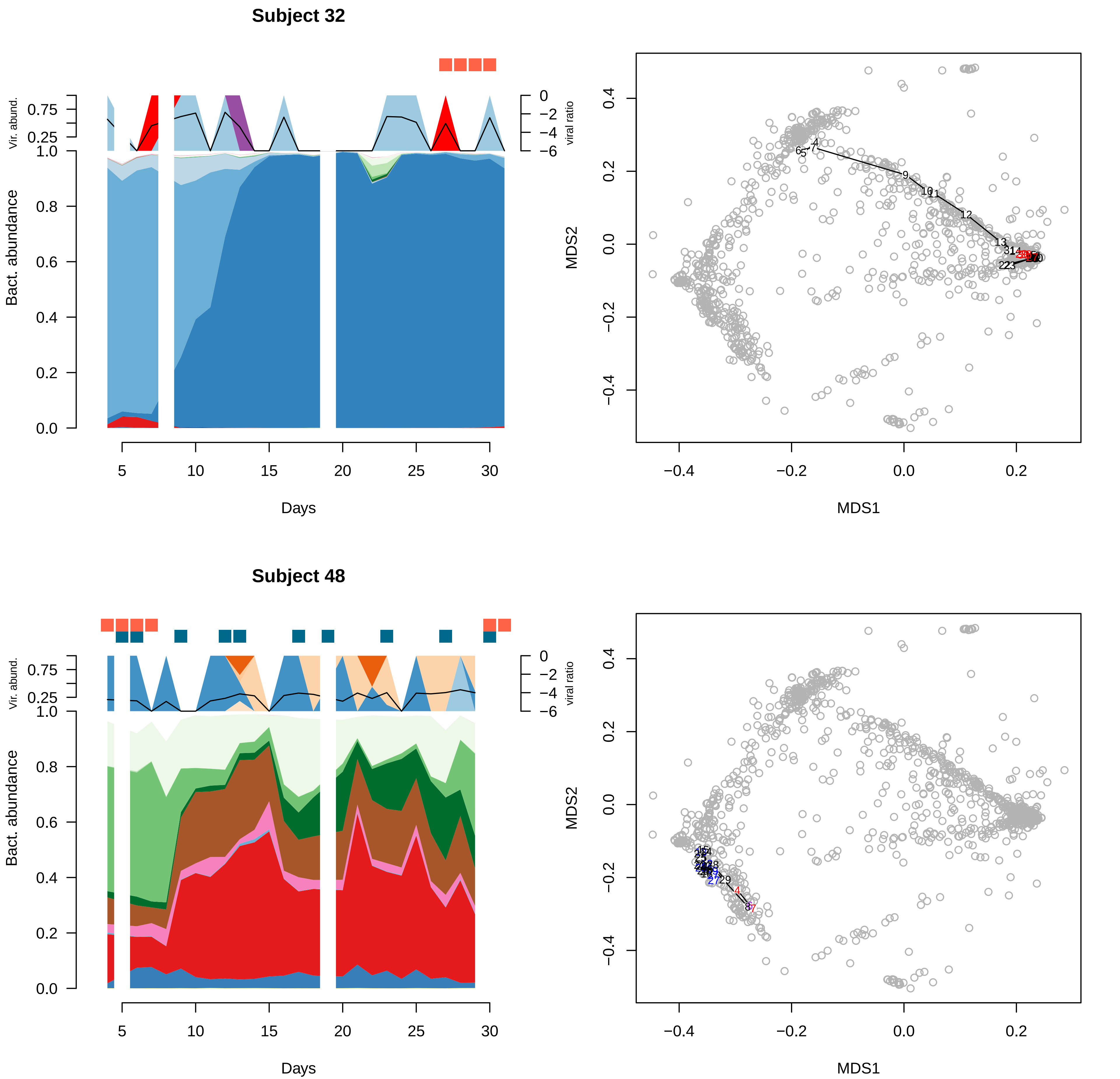

Subject 32
Vir. abund.
0.75
0.25
1.0
viral ratio
0
−2
−4
−6
Bact. abundance
0.8
0.6
0.4
0.2
0.0
5
10
15
20
25
30
Days
MDS2
MDS1
0.4
0.2
0.0
−0.2
−0.4
Subject 48

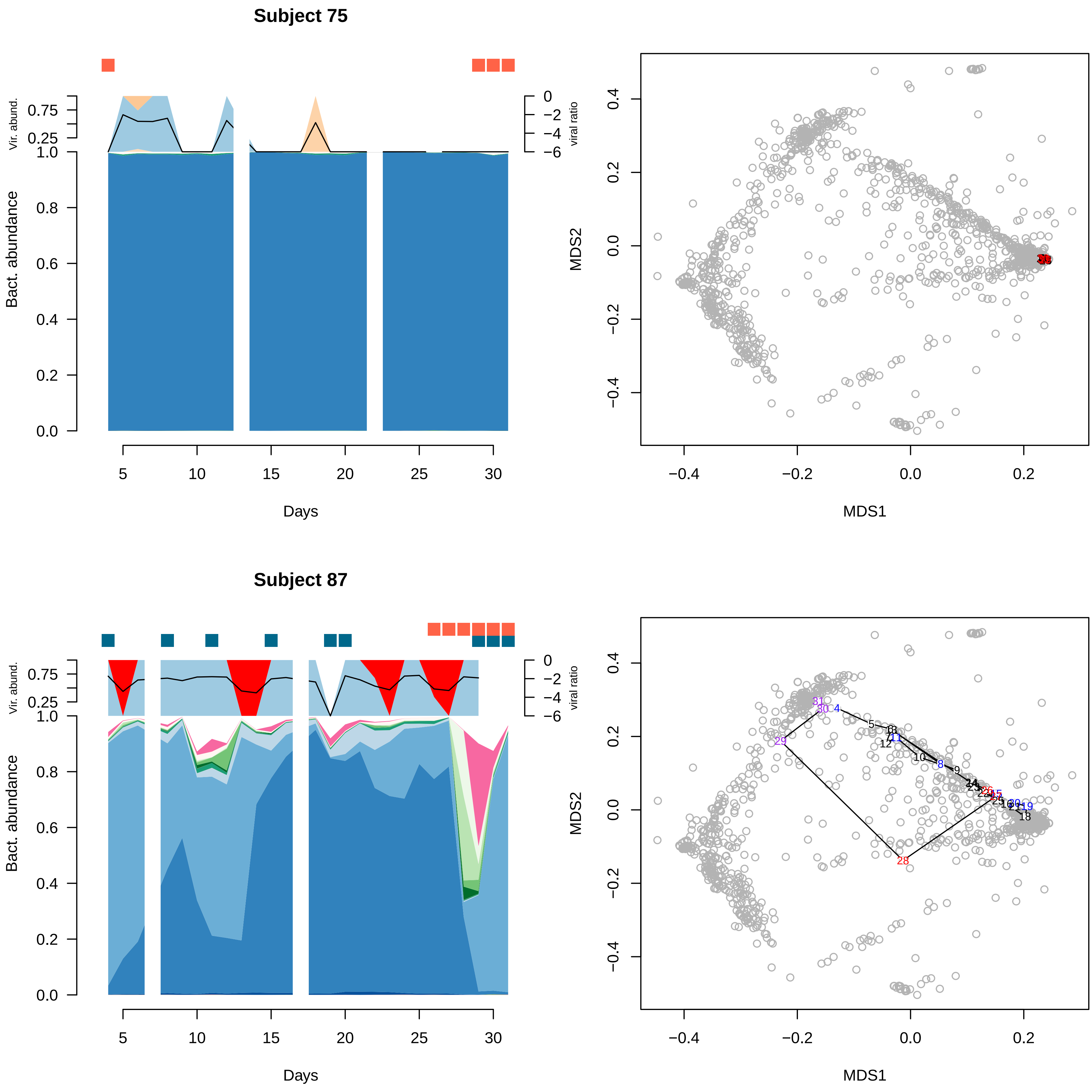

Subject 75
Vir. abund.
0.75
0.25
viral ratio
0
−2
−4
−6
Bact. abundance
1.0
0.8
0.6
0.4
0.2
0.0
5
10
15
20
25
30
Days
MDS2
MDS1
0.4
0.2
0.0
−0.2
−0.4
Subject 87
Vir. abund.
viral ratio
Bact. abundance
Days
MDS2
MDS1

**Subject 117**

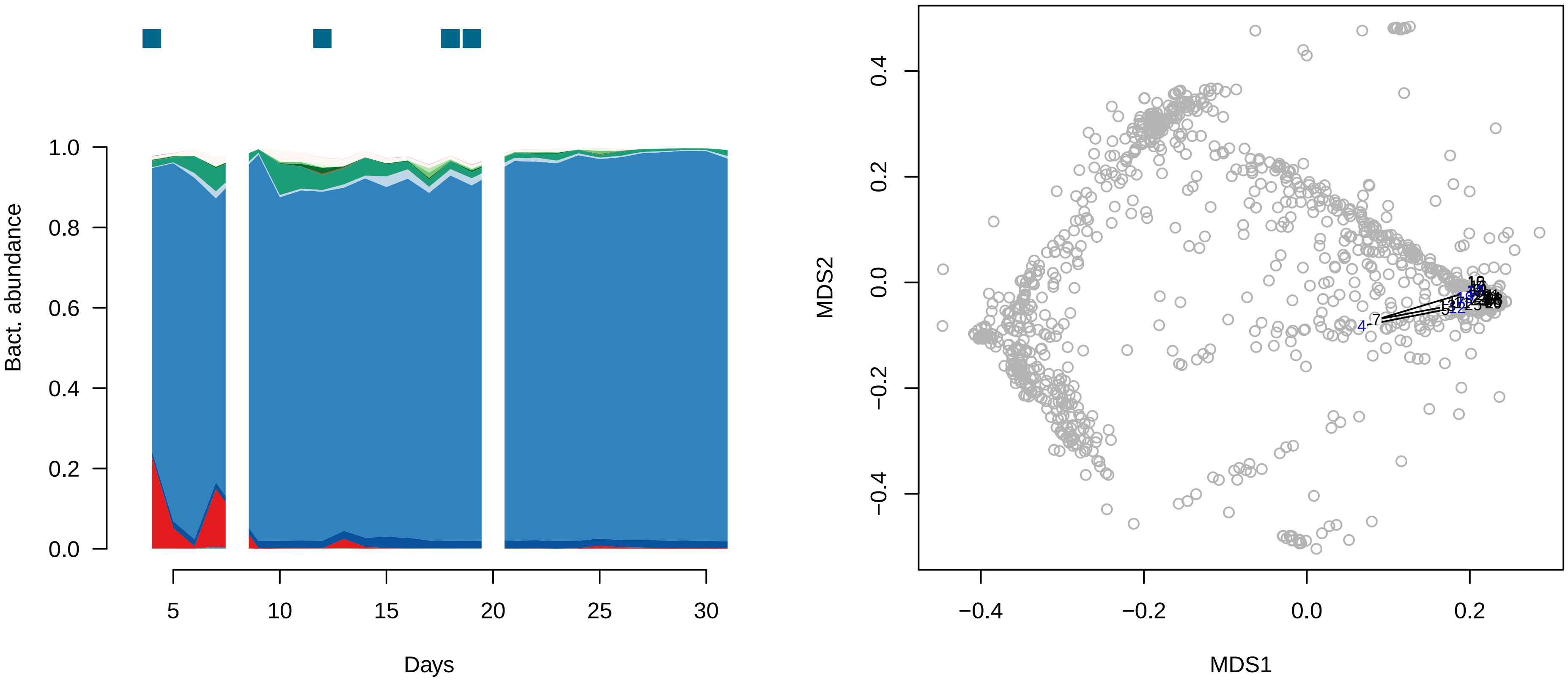


**Subject 137**

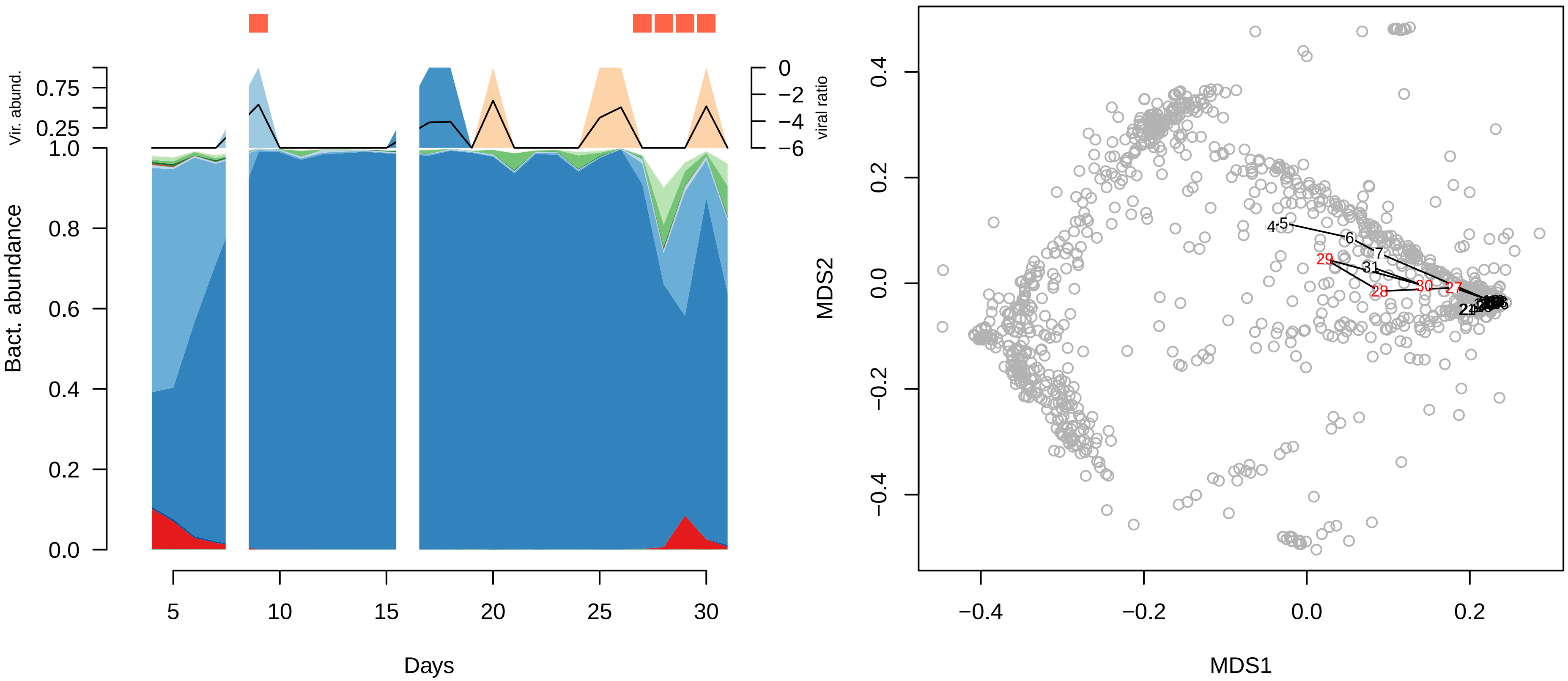

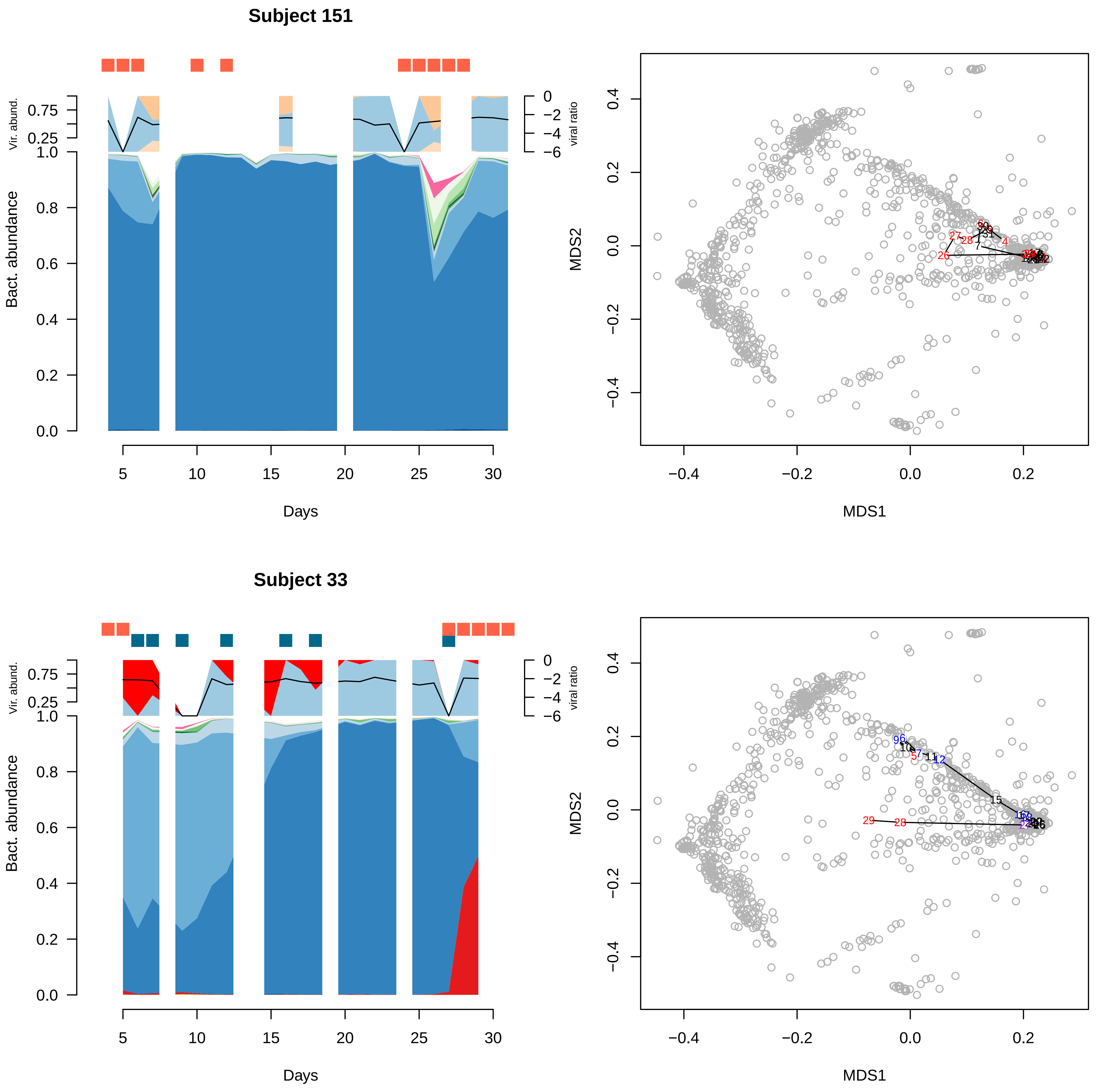
Subject 151
Vir. abund.
viral ratio
Bact. abundance
Days
MDS1
MDS2
Subject 33
Vir. abund.
viral ratio
Bact. abundance
Days
MDS1
MDS2

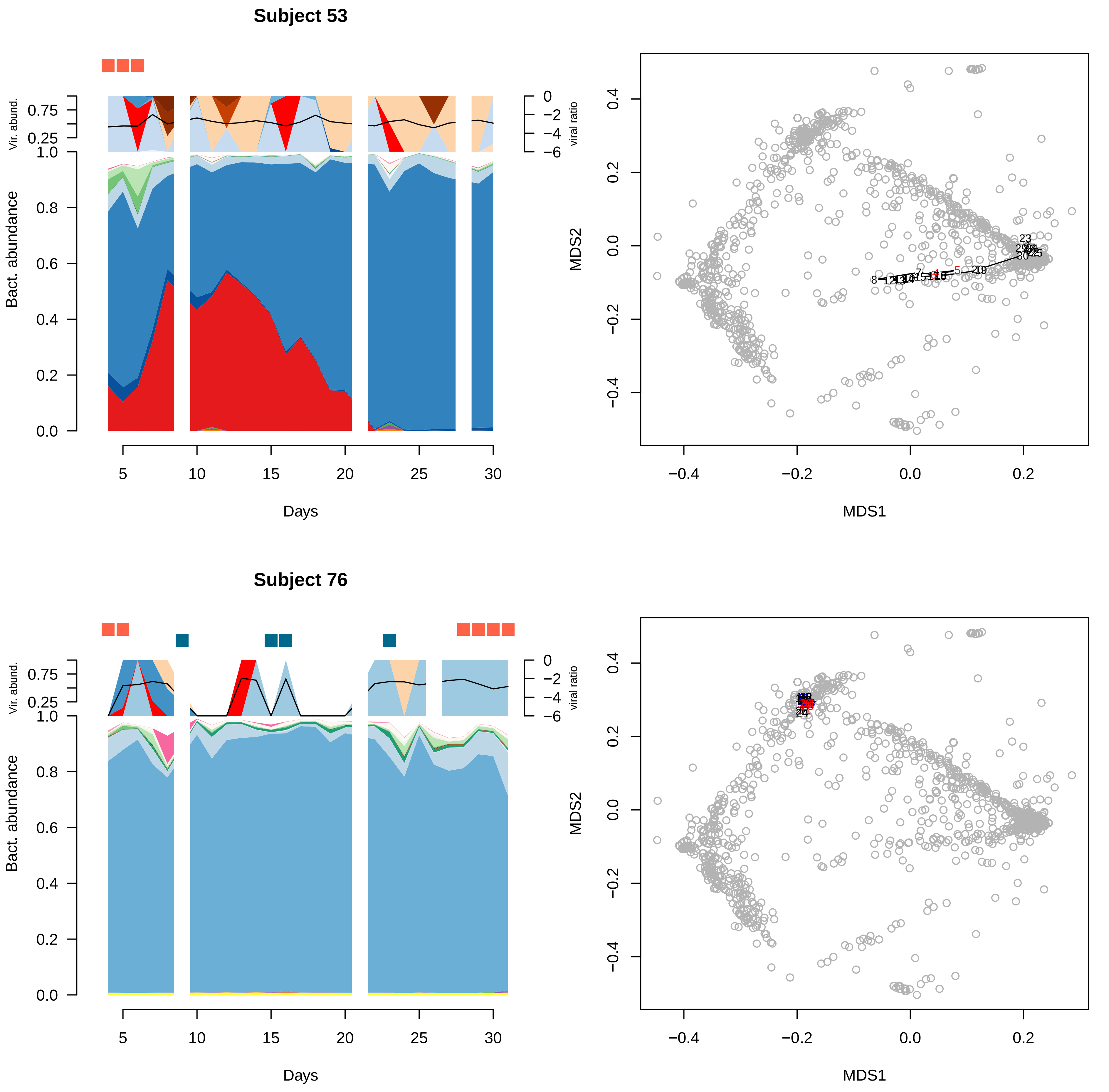

Subject 53
Vir. abund.
0.75
0.25
Bact. abundance
1.0
0.8
0.6
0.4
0.2
0.0
viral ratio
0
−2
−4
−6
5
10
15
20
25
30
Days
MDS2
0.4
0.2
0.0
−0.2
−0.4
MDS1
−0.4
−0.2
0.0
0.2
Subject 76
Vir. abund.
0.75
0.25
Bact. abundance
1.0
0.8
0.6
0.4
0.2
0.0
viral ratio
0
−2
−4
−6
5
10
15
20
25
30
Days
MDS2
0.4
0.2
0.0
−0.2
−0.4
MDS1
−0.4
−0.2
0.0
0.2

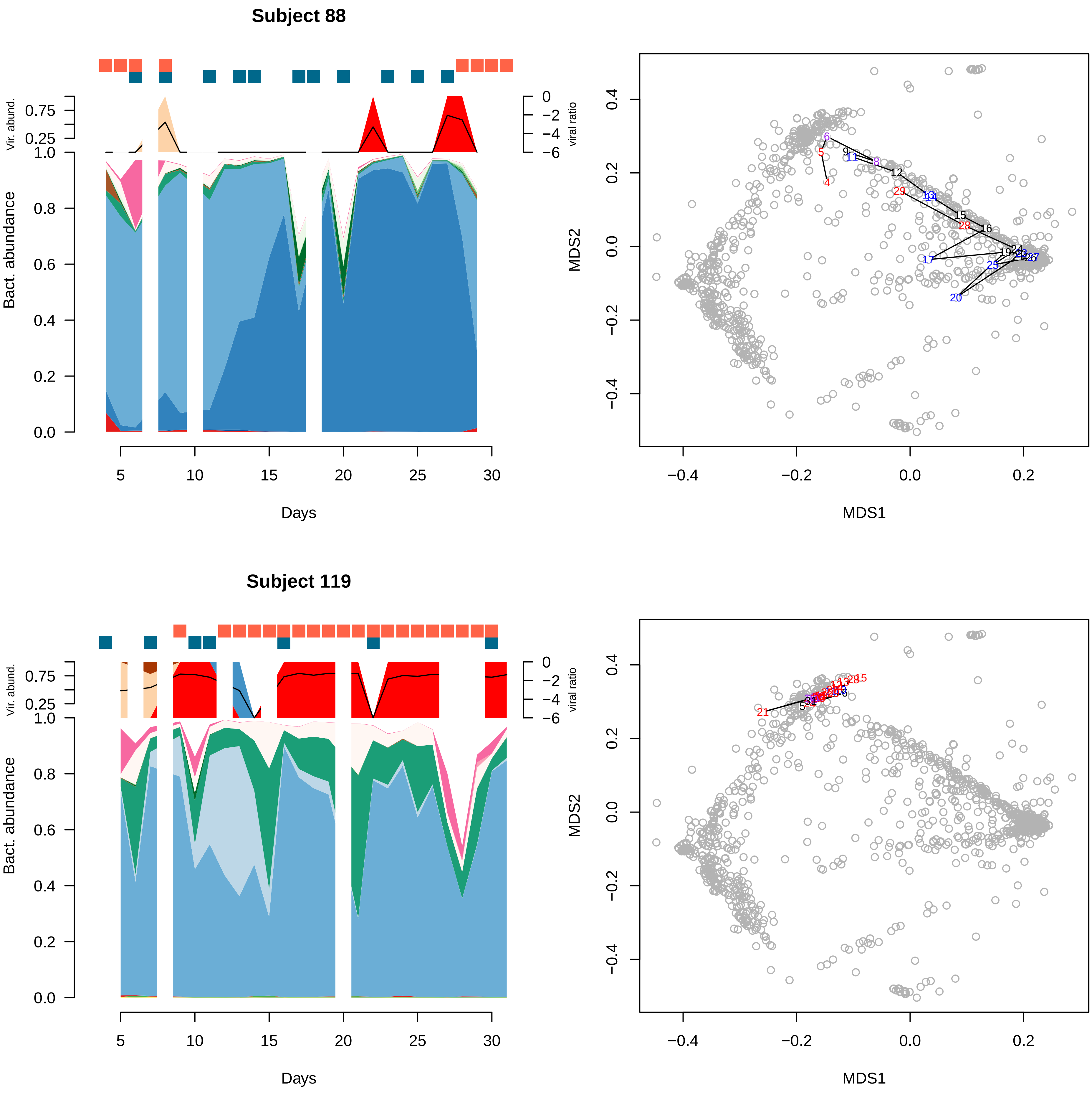

Subject 88
Vir. abund.
viral ratio
Bact. abundance
Days
MDS1
MDS2
Subject 119
Vir. abund.
viral ratio
Bact. abundance
Days
MDS1
MDS2

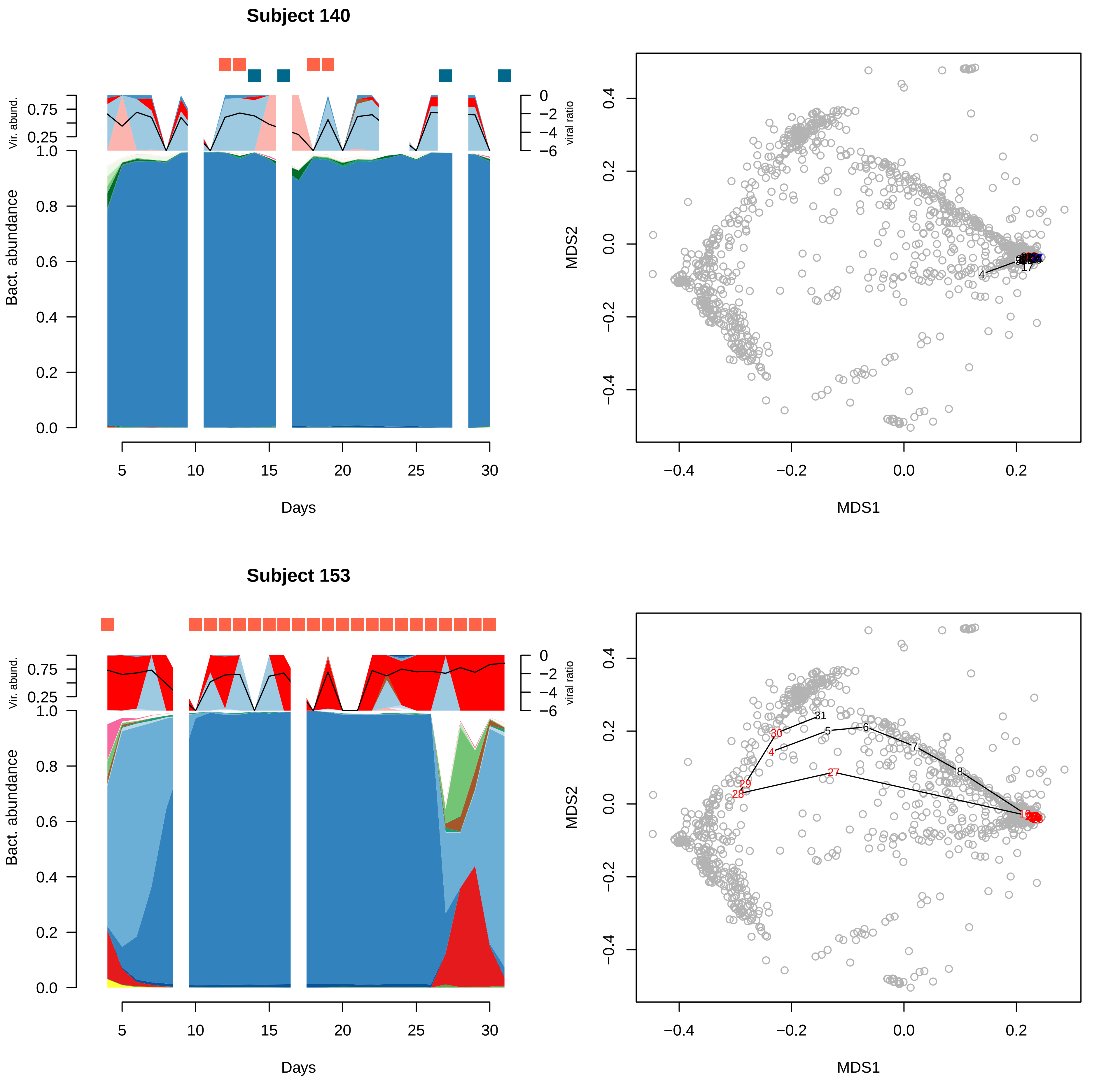

Subject 140
Vir. abund.
viral ratio
Bact. abundance
Days
MDS1
MDS2
Subject 153

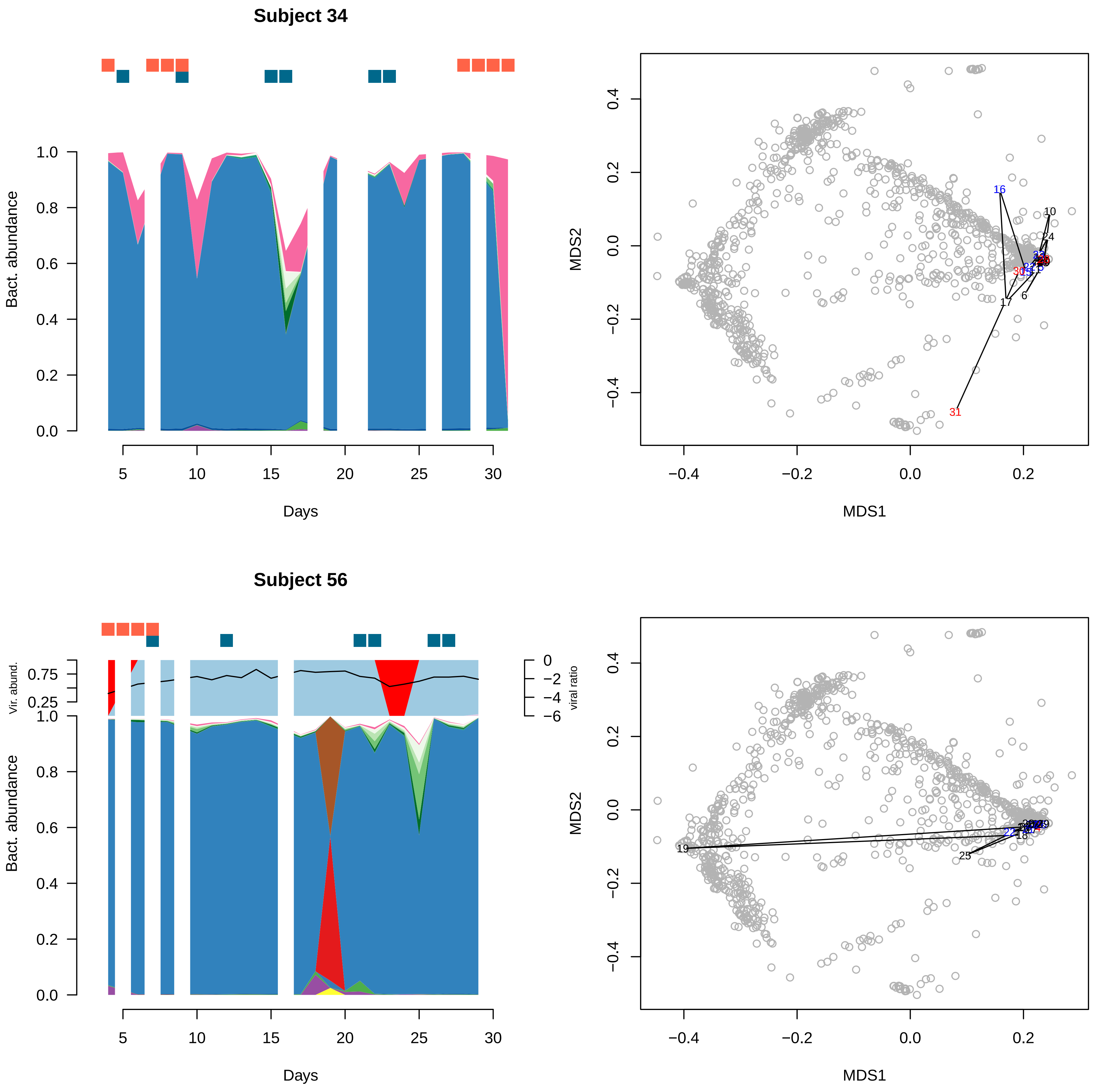
Subject 34
Bact. abundance
Days
MDS1
MDS2
Subject 56
Vir. abund.
viral ratio
Bact. abundance
Days
MDS1
MDS2

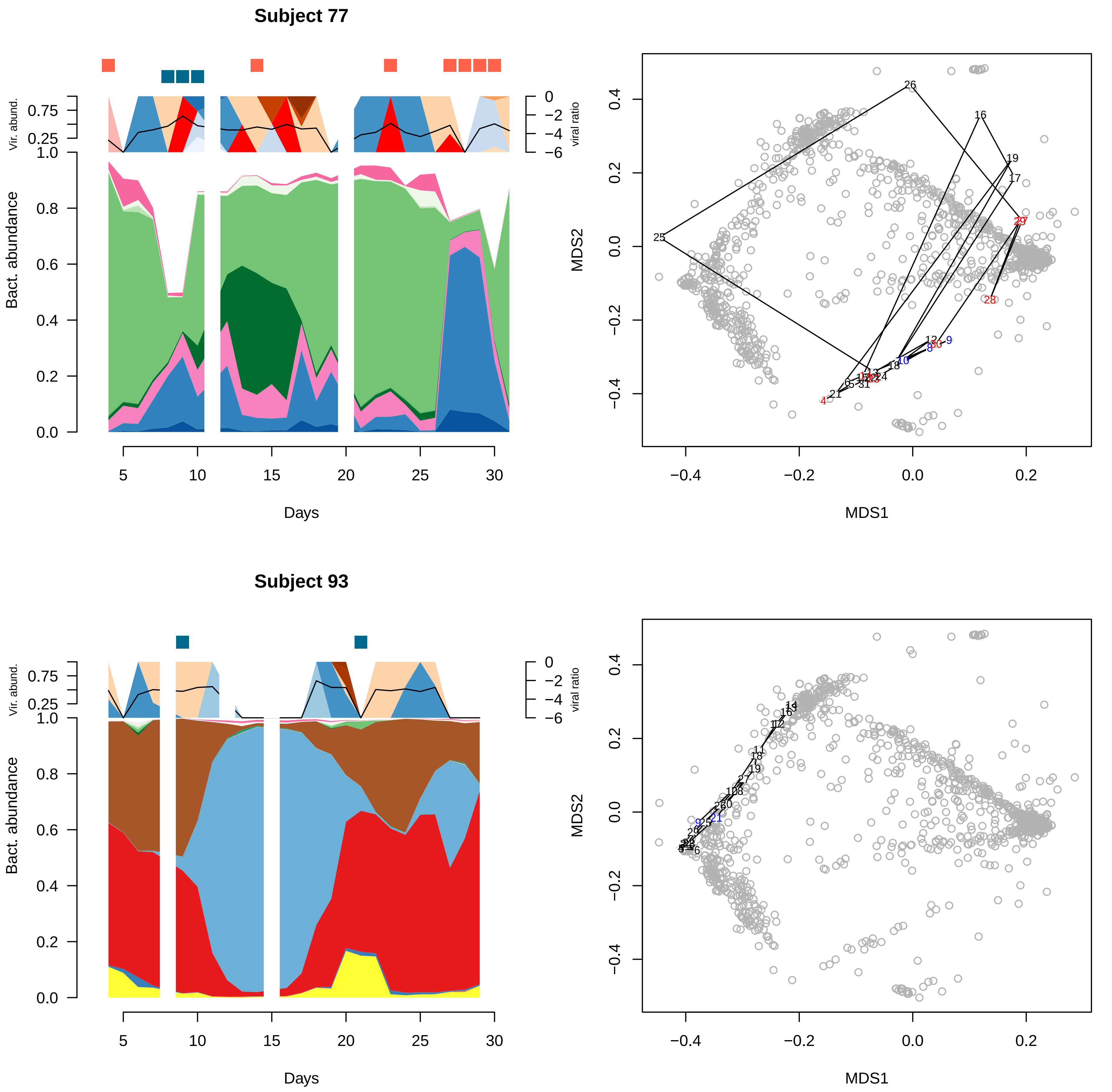
Subject 77
Vir. abund.
viral ratio
Bact. abundance
Days
MDS1
MDS2
Subject 93
Vir. abund.
viral ratio
Bact. abundance
Days
MDS1
MDS2

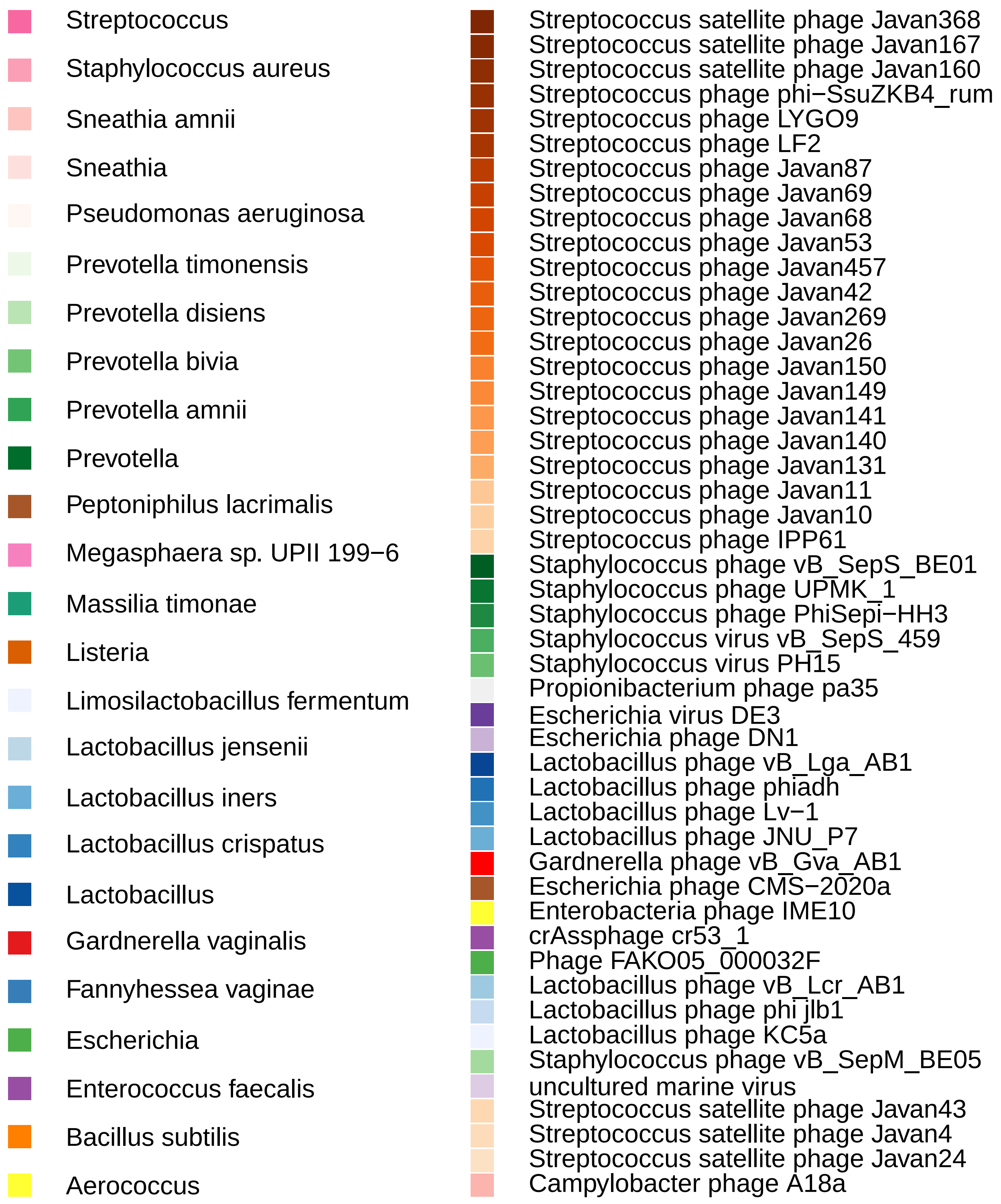
Streptococcus
Staphylococcus aureus
Sneathia amnii
Sneathia
Pseudomonas aeruginosa
Prevotella timonensis
Prevotella disiens
Prevotella bivia
Prevotella amnii
Prevotella
Peptoniphilus lacrimalis
Megasphaera sp. UPII 199−6
Massilia timonae
Listeria
Limosilactobacillus fermentum
Lactobacillus jensenii
Lactobacillus iners
Lactobacillus crispatus
Lactobacillus
Gardnerella vaginalis
Fannyhessea vaginae
Escherichia
Enterococcus faecalis
Bacillus subtilis
Aerococcus
Streptococcus satellite phage Javan368
Streptococcus satellite phage Javan167
Streptococcus satellite phage Javan160
Streptococcus phage phi−SsuZKB4_rum
Streptococcus phage LYGO9
Streptococcus phage LF2
Streptococcus phage Javan87
Streptococcus phage Javan69
Streptococcus phage Javan68
Streptococcus phage Javan53
Streptococcus phage Javan457
Streptococcus phage Javan42
Streptococcus phage Javan269
Streptococcus phage Javan26
Streptococcus phage Javan150
Streptococcus phage Javan149
Streptococcus phage Javan141
Streptococcus phage Javan140
Streptococcus phage Javan131
Streptococcus phage Javan11
Streptococcus phage Javan10
Streptococcus phage IPP61
Staphylococcus phage vB_SepS_BE01
Staphylococcus phage UPMK_1
Staphylococcus phage PhiSepi−HH3
Staphylococcus virus vB_SepS_459
Staphylococcus virus PH15
Propionibacterium phage pa35
Escherichia virus DE3
Escherichia phage DN1
Lactobacillus phage vB_Lga_AB1
Lactobacillus phage phiadh
Lactobacillus phage Lv−1
Lactobacillus phage JNU_P7
Gardnerella phage vB_Gva_AB1
Escherichia phage CMS−2020a
Enterobacteria phage IME10
crAssphage cr53_1
Phage FAKO05_000032F
Lactobacillus phage vB_Lcr_AB1
Lactobacillus phage phi jlb1
Lactobacillus phage KC5a
Staphylococcus phage vB_SepM_BE05
uncultured marine virus
Streptococcus satellite phage Javan43
Streptococcus satellite phage Javan4
Streptococcus satellite phage Javan24
Campylobacter phage A18a

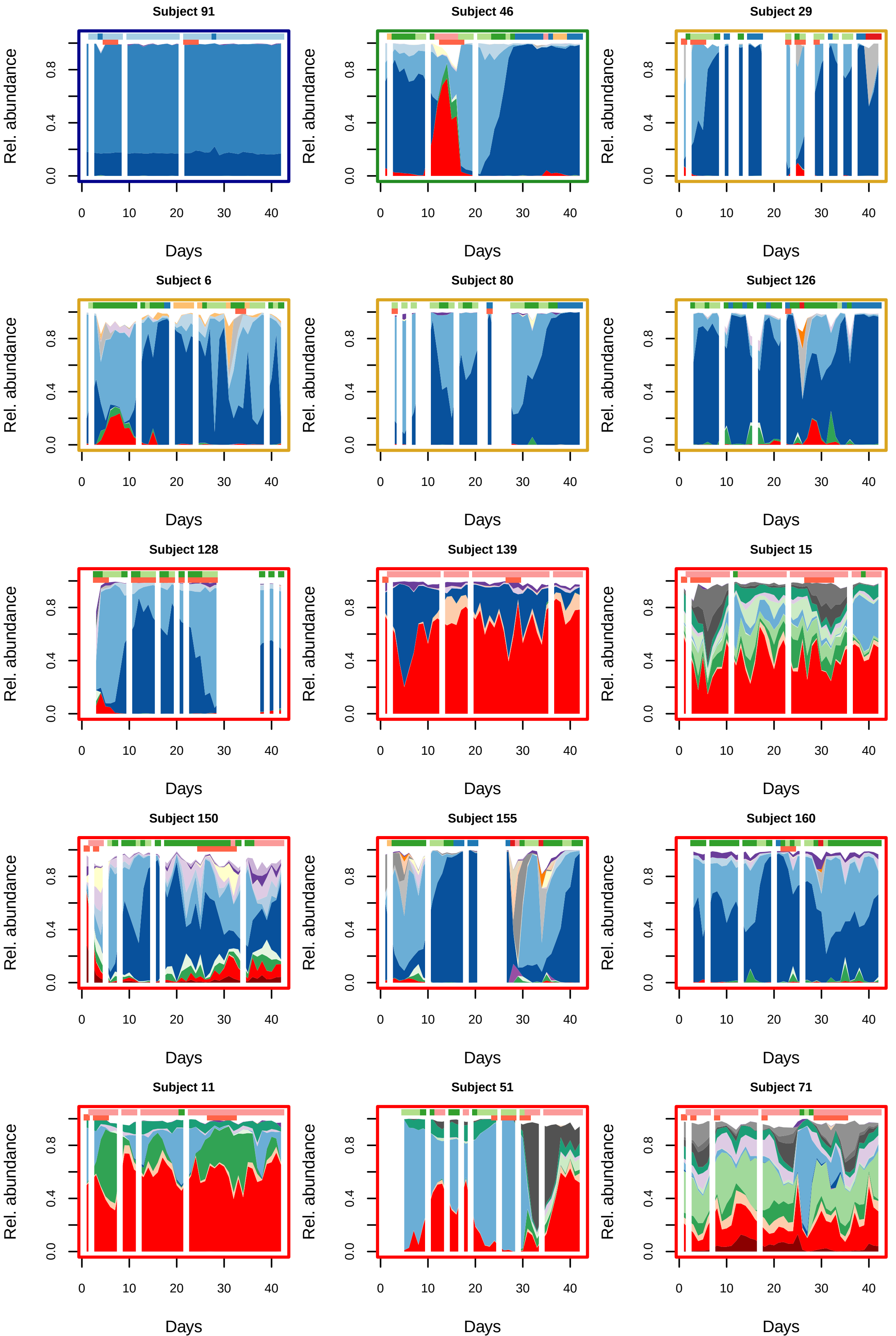
Subject 91
Subject 46
Subject 29
Subject 6
Subject 80
Subject 126
Subject 128
Subject 139
Subject 15
Subject 150
Subject 155
Subject 160
Subject 11
Subject 51
Subject 71
Rel. abundance
0.0
0.4
0.8
0
10
20
30
40
Days

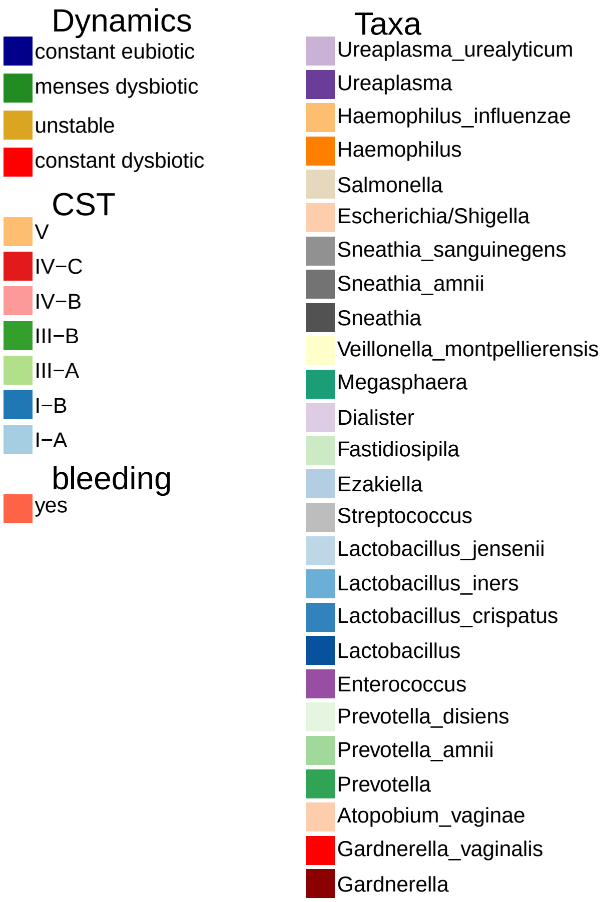
Dynamics
constant eubiotic
menses dysbiotic
unstable
constant dysbiotic
CST
V
IV−C
IV−B
III−B
III−A
I−B
I−A
bleeding
yes
Taxa
Ureaplasma_urealyticum
Ureaplasma
Haemophilus_influenzae
Haemophilus
Salmonella
Escherichia/Shigella
Sneathia_sanguinegens
Sneathia_amnii
Sneathia
Veillonella_montpellierensis
Megasphaera
Dialister
Fastidiosipila
Ezakiella
Streptococcus
Lactobacillus_jensenii
Lactobacillus_iners
Lactobacillus_crispatus
Lactobacillus
Enterococcus
Prevotella_disiens
Prevotella_amnii
Prevotella
Atopobium_vaginae
Gardnerella_vaginalis
Gardnerella

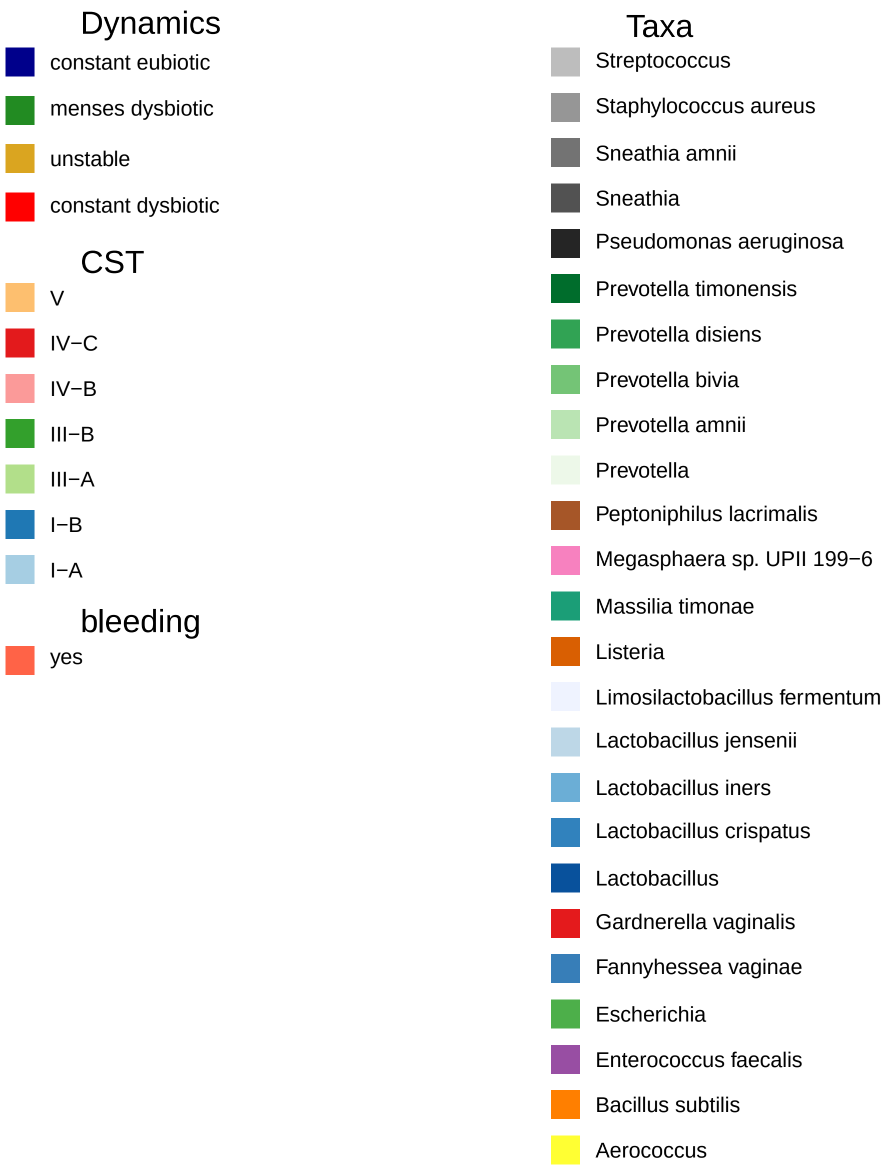
Dynamics
constant eubiotic
menses dysbiotic
unstable
constant dysbiotic
CST
V
IV−C
IV−B
III−B
III−A
I−B
I−A
bleeding
yes
Taxa
Streptococcus
Staphylococcus aureus
Sneathia amnii
Sneathia
Pseudomonas aeruginosa
Prevotella timonensis
Prevotella disiens
Prevotella bivia
Prevotella amnii
Prevotella
Peptoniphilus lacrimalis
Megasphaera sp. UPII 199−6
Massilia timonae
Listeria
Limosilactobacillus fermentum
Lactobacillus jensenii
Lactobacillus iners
Lactobacillus crispatus
Lactobacillus
Gardnerella vaginalis
Fannyhessea vaginae
Escherichia
Enterococcus faecalis
Bacillus subtilis
Aerococcus

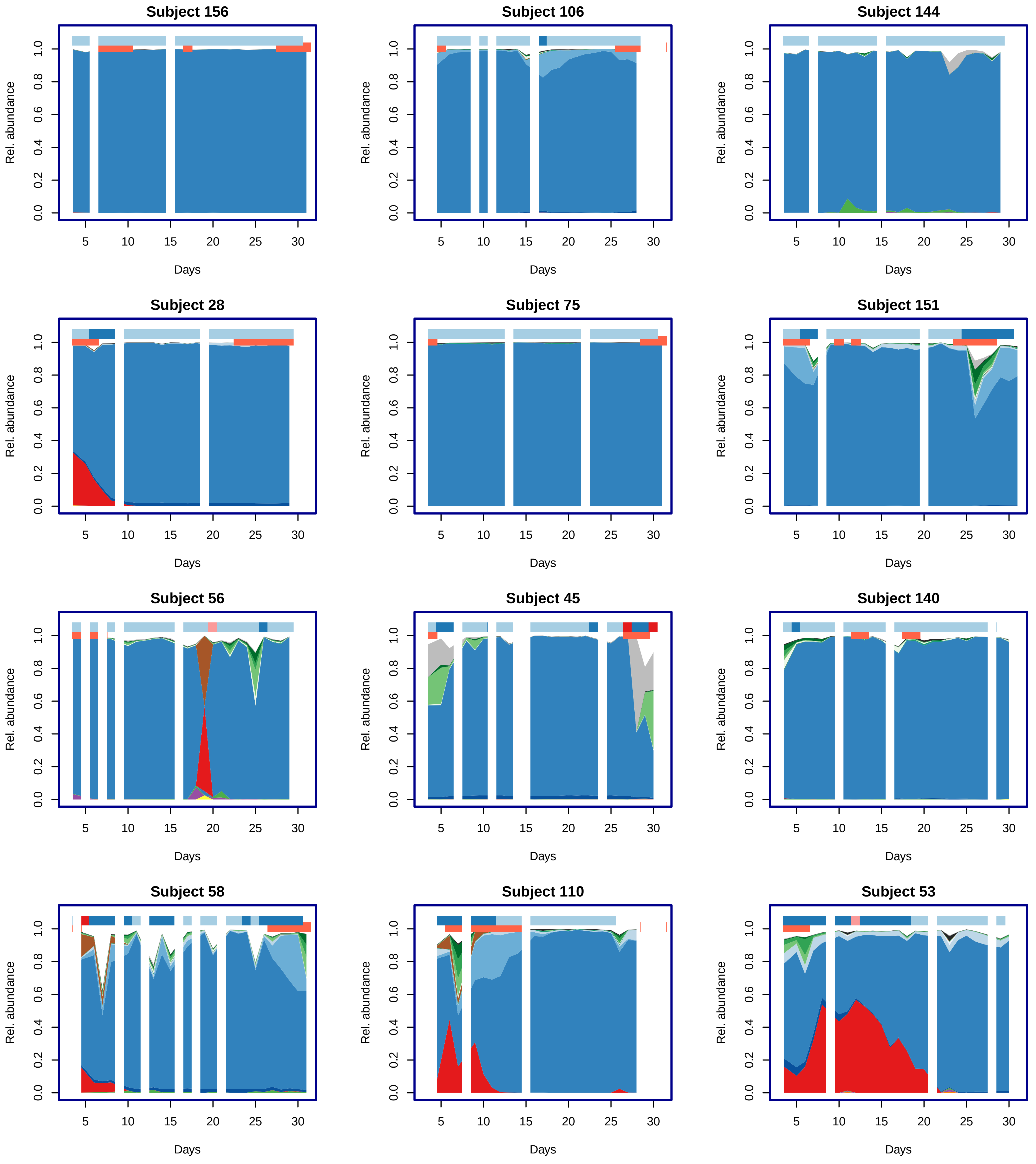
Subject 156
Subject 106
Subject 144
Subject 28
Subject 75
Subject 151
Subject 56
Subject 45
Subject 140
Subject 58
Subject 110
Subject 53
Rel. abundance
0.0
0.2
0.4
0.6
0.8
1.0
Days
5
10
15
20
25
30

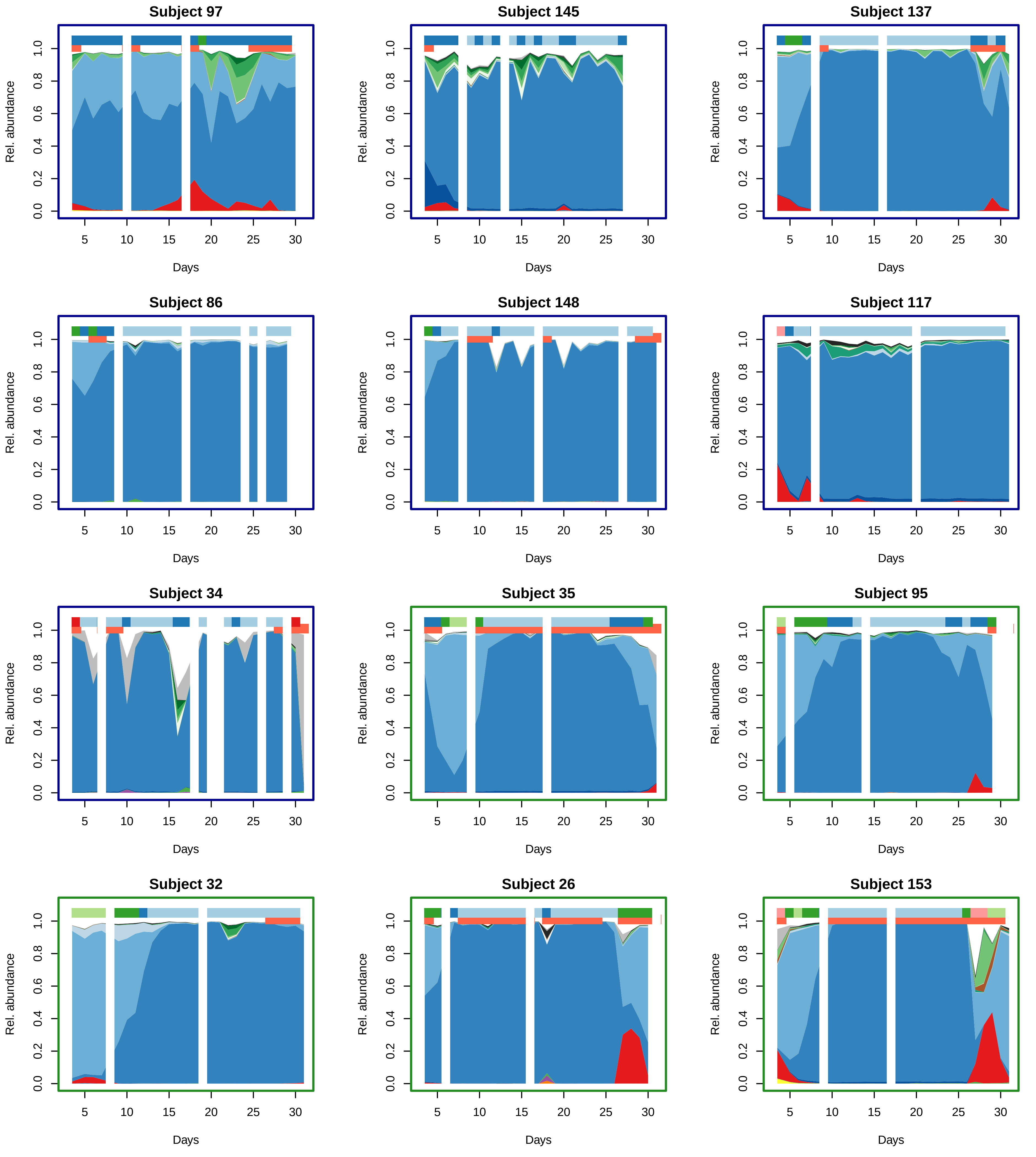

Subject 97
Subject 145
Subject 137
Subject 86
Subject 148
Subject 117
Subject 34
Subject 35
Subject 95
Subject 32
Subject 26
Subject 153
Rel. abundance
Days

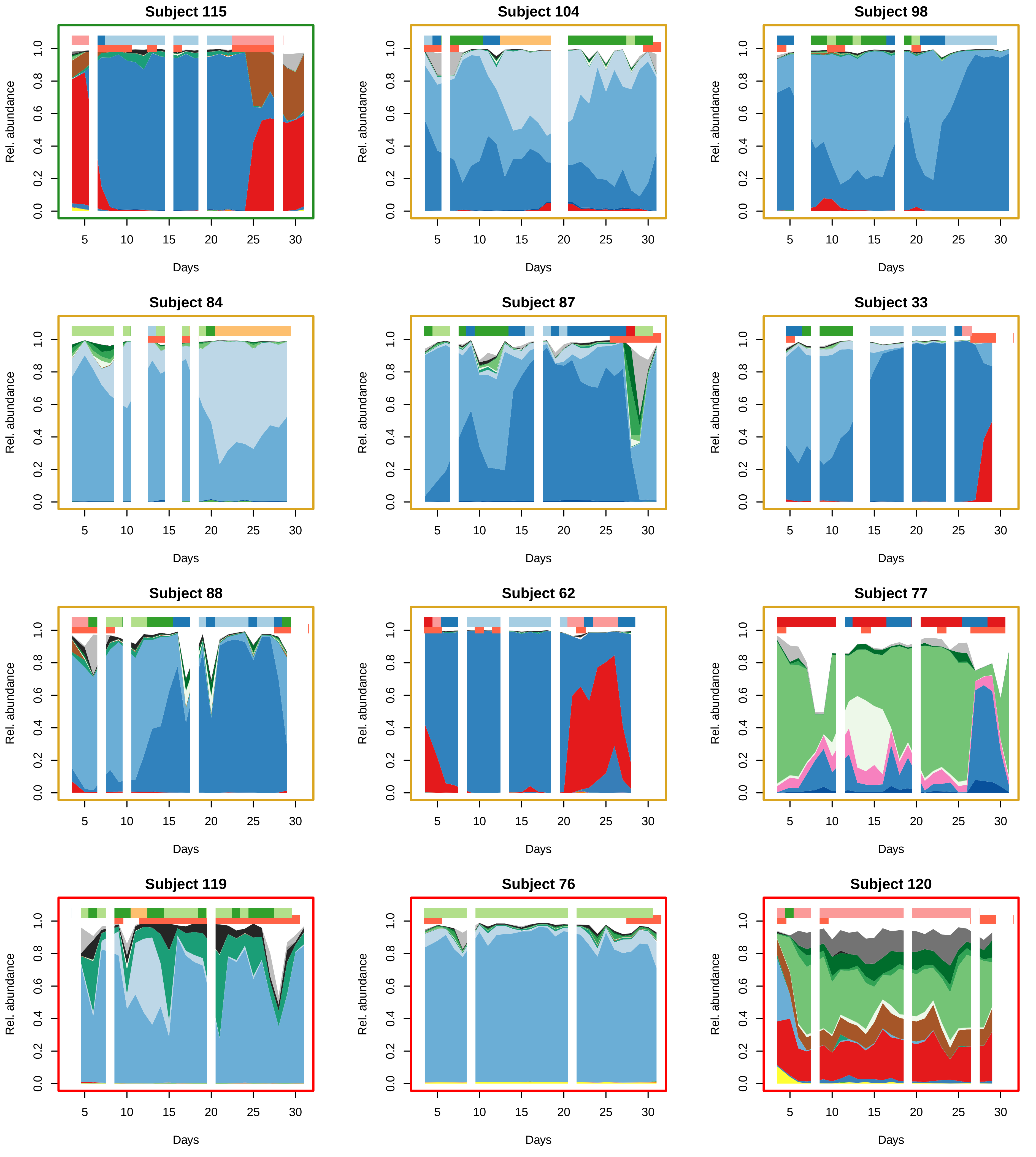

Subject 115
Subject 104
Subject 98
Subject 84
Subject 87
Subject 33
Subject 88
Subject 62
Subject 77
Subject 119
Subject 76
Subject 120
Rel. abundance
0.0
0.2
0.4
0.6
0.8
1.0
5
10
15
20
25
30
Days

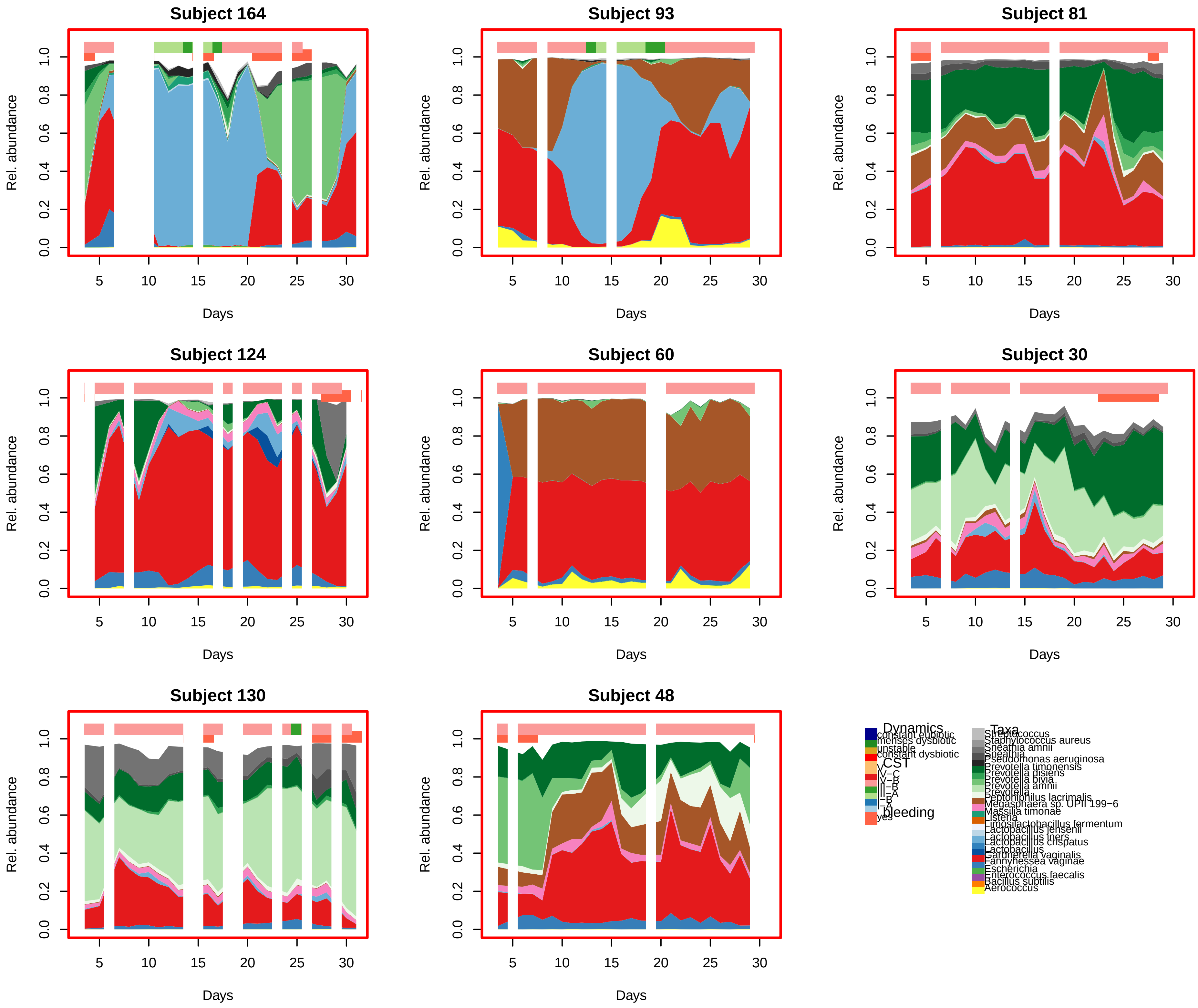
Subject 164
Subject 93
Subject 81
Subject 124
Subject 60
Subject 30
Subject 130
Subject 48
Rel. abundance
0.0
0.2
0.4
0.6
0.8
1.0
5
10
15
20
25
30
Days
Dynamics
constant eubiotic
menses dysbiotic
unstable
constant dysbiotic
CST
V
IV−C
IV−B
III−B
III−A
I−B
I−A
bleeding
yes
Taxa
Streptococcus
Staphylococcus aureus
Sneathia amnii
Sneathia
Pseudomonas aeruginosa
Prevotella timonensis
Prevotella disiens
Prevotella bivia.
Prevotella amnii
Prevotella
Peptoniphilus lacrimalis
Megasphaera sp. UPII 199−6
Massilia timonae
Listeria
Limosilactobacillus fermentum
Lactobacillus jensenii
Lactobacillus iners
Lactobacillus crispatus
Lactobacillus
Gardnerella vaginalis
Fannyhessea vaginae
Escherichia
Enterococcus faecalis
Bacillus subtilis
Aerococcus

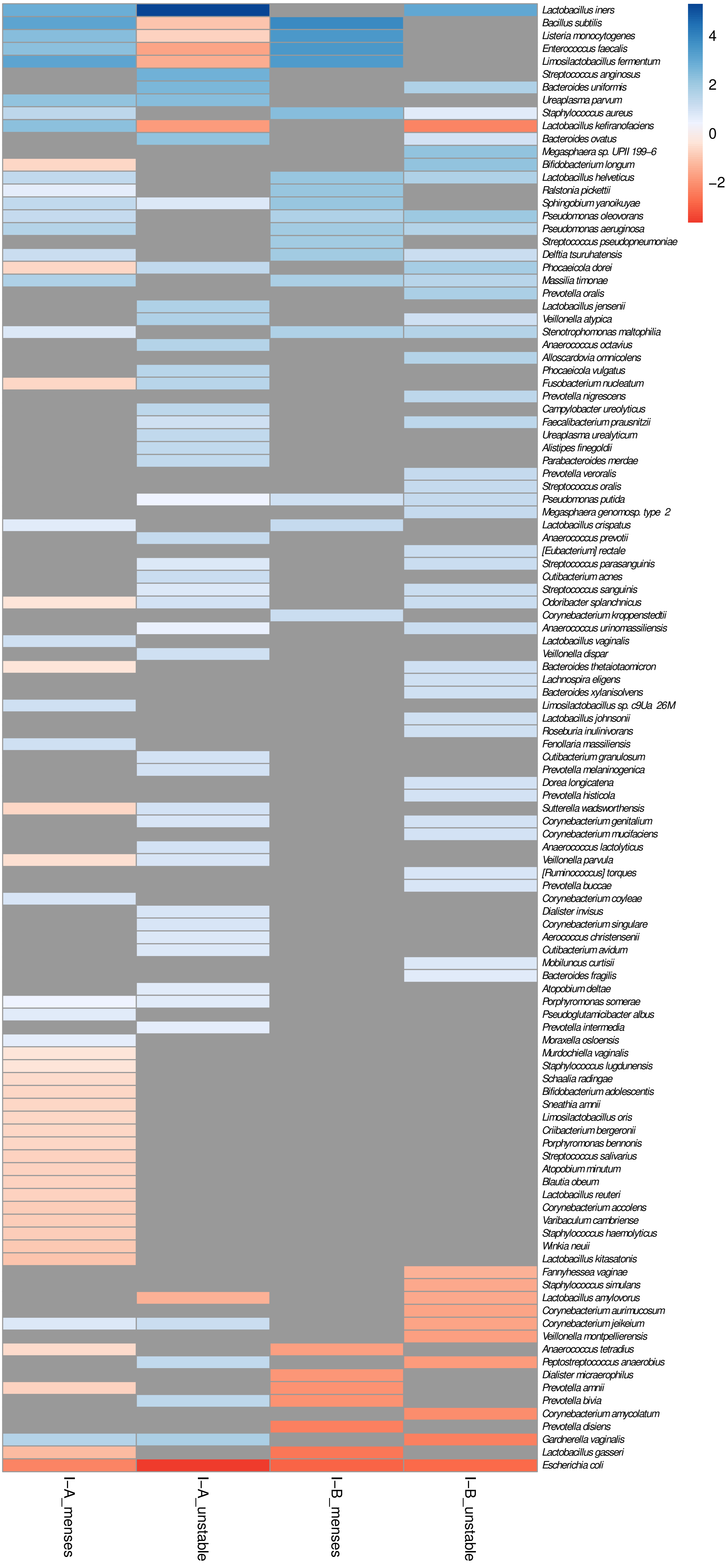
4
2
0
−2
Lactobacillus iners
Bacillus subtilis
Listeria monocytogenes
Enterococcus faecalis
Limosilactobacillus fermentum
Streptococcus anginosus
Bacteroides uniformis
Ureaplasma parvum
Staphylococcus aureus
Lactobacillus kefiranofaciens
Bacteroides ovatus
Megasphaera sp. UPII 199−6
Bifidobacterium longum
Lactobacillus helveticus
Ralstonia pickettii
Sphingobium yanoikuyae
Pseudomonas oleovorans
Pseudomonas aeruginosa
Streptococcus pseudopneumoniae
Delftia tsuruhatensis
Phocaeicola dorei
Massilia timonae
Prevotella oralis
Lactobacillus jensenii
Veillonella atypica
Stenotrophomonas maltophilia
Anaerococcus octavius
Alloscardovia omnicolens
Phocaeicola vulgatus
Fusobacterium nucleatum
Prevotella nigrescens
Campylobacter ureolyticus
Faecalibacterium prausnitzii
Ureaplasma urealyticum
Alistipes finegoldii
Parabacteroides merdae
Prevotella veroralis
Streptococcus oralis
Pseudomonas putida
Megasphaera genomosp. type_2
Lactobacillus crispatus
Anaerococcus prevotii
[Eubacterium] rectale
Streptococcus parasanguinis
Cutibacterium acnes
Streptococcus sanguinis
Odoribacter splanchnicus
Corynebacterium kroppenstedtii
Anaerococcus urinomassiliensis
Lactobacillus vaginalis
Veillonella dispar
Bacteroides thetaiotaomicron
Lachnospira eligens
Bacteroides xylanisolvens
Limosilactobacillus sp. c9Ua_26M
Lactobacillus johnsonii
Roseburia inulinivorans
Fenollaria massiliensis
Cutibacterium granulosum
Prevotella melaninogenica
Dorea longicatena
Prevotella histicola
Sutterella wadsworthensis
Corynebacterium genitalium
Corynebacterium mucifaciens
Anaerococcus lactolyticus
Veillonella parvula
[Ruminococcus] torques
Prevotella buccae
Corynebacterium coyleae
Dialister invisus
Corynebacterium singulare
Aerococcus christensenii
Cutibacterium avidum
Mobiluncus curtisii
Bacteroides fragilis
Atopobium deltae
Porphyromonas somerae
Pseudoglutamicibacter albus
Prevotella intermedia
Moraxella osloensis
Murdochiella vaginalis
Staphylococcus lugdunensis
Schaalia radingae
Bifidobacterium adolescentis
Sneathia amnii
Limosilactobacillus oris
Criibacterium bergeronii
Porphyromonas bennonis
Streptococcus salivarius
Atopobium minutum
Blautia obeum
Lactobacillus reuteri
Corynebacterium accolens
Varibaculum cambriense
Staphylococcus haemolyticus
Winkia neuii
Lactobacillus kitasatonis
Fannyhessea vaginae
Staphylococcus simulans
Lactobacillus amylovorus
Corynebacterium aurimucosum
Corynebacterium jeikeium
Veillonella montpellierensis
Anaerococcus tetradius
Peptostreptococcus anaerobius
Dialister micraerophilus
Prevotella amnii
Prevotella bivia
Corynebacterium amycolatum
Prevotella disiens
Gardnerella vaginalis
Lactobacillus gasseri
Escherichia coli
I−A_menses
I−A_unstable
I−B_menses
I−B_unstable

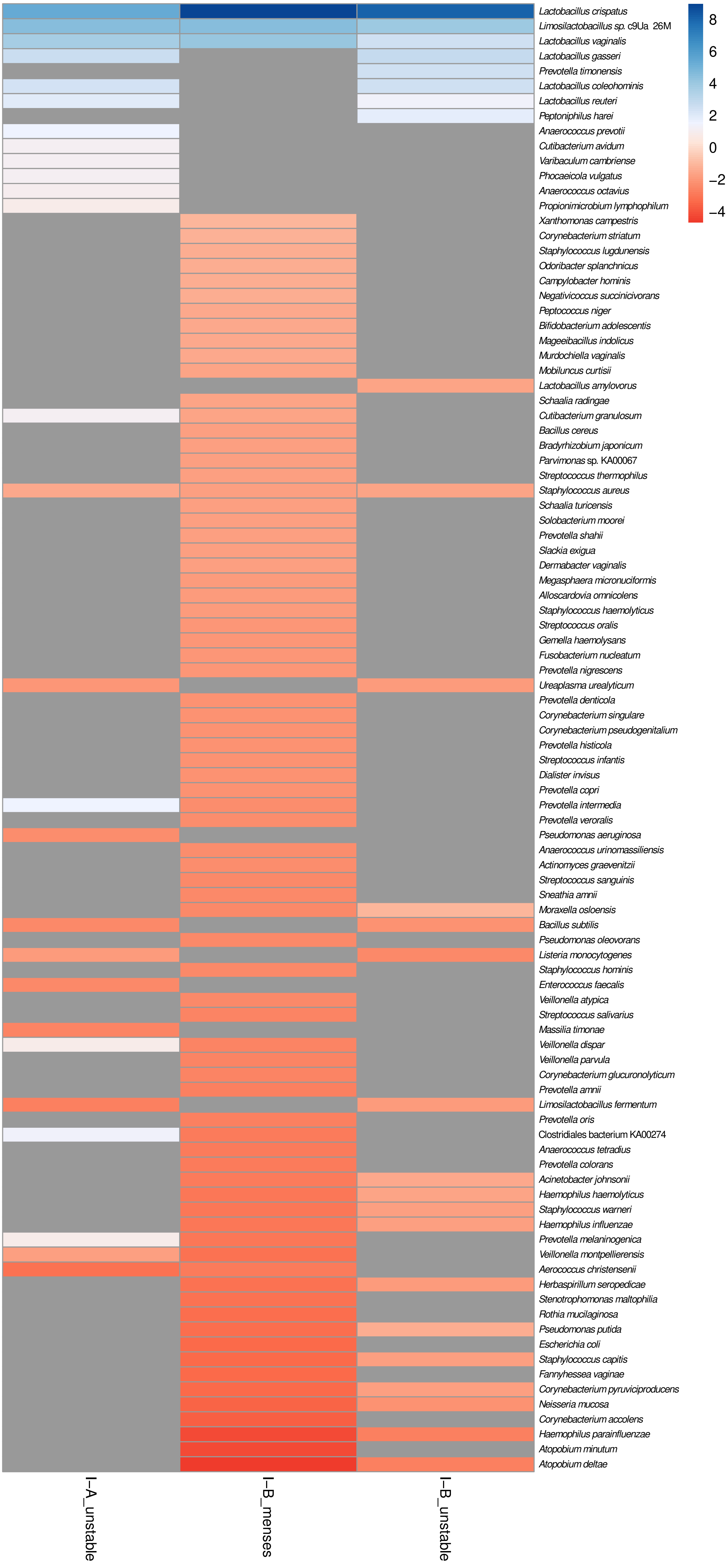
Lactobacillus crispatus
Limosilactobacillus sp. c9Ua 26M
Lactobacillus vaginalis
Lactobacillus gasseri
Prevotella timonensis
Lactobacillus coleohominis
Lactobacillus reuteri
Peptoniphilus harei
Anaerococcus prevotii
Cutibacterium avidum
Varibaculum cambriense
Phocaeicola vulgatus
Anaerococcus octavius
Propionimicrobium lymphophilum
Xanthomonas campestris
Corynebacterium striatum
Staphylococcus lugdunensis
Odoribacter splanchnicus
Campylobacter hominis
Negativicoccus succinicivorans
Peptococcus niger
Bifidobacterium adolescentis
Mageeibacillus indolicus
Murdochiella vaginalis
Mobiluncus curtisii
Lactobacillus amylovorus
Schaalia radingae
Cutibacterium granulosum
Bacillus cereus
Bradyrhizobium japonicum
Parvimonas sp. KA00067
Streptococcus thermophilus
Staphylococcus aureus
Schaalia turicensis
Solobacterium moorei
Prevotella shahii
Slackia exigua
Dermabacter vaginalis
Megasphaera micronuciformis
Alloscardovia omnicolens
Staphylococcus haemolyticus
Streptococcus oralis
Gemella haemolysans
Fusobacterium nucleatum
Prevotella nigrescens
Ureaplasma urealyticum
Prevotella denticola
Corynebacterium singulare
Corynebacterium pseudogenitalium
Prevotella histicola
Streptococcus infantis
Dialister invisus
Prevotella copri
Prevotella intermedia
Prevotella veroralis
Pseudomonas aeruginosa
Anaerococcus urinomassiliensis
Actinomyces graevenitzii
Streptococcus sanguinis
Sneathia amnii
Moraxella osloensis
Bacillus subtilis
Pseudomonas oleovorans
Listeria monocytogenes
Staphylococcus hominis
Enterococcus faecalis
Veillonella atypica
Streptococcus salivarius
Massilia timonae
Veillonella dispar
Veillonella parvula
Corynebacterium glucuronolyticum
Prevotella amnii
Limosilactobacillus fermentum
Prevotella oris
Clostridiales bacterium KA00274
Anaerococcus tetradius
Prevotella colorans
Acinetobacter johnsonii
Haemophilus haemolyticus
Staphylococcus warneri
Haemophilus influenzae
Prevotella melaninogenica
Veillonella montpellierensis
Aerococcus christensenii
Herbaspirillum seropedicae
Stenotrophomonas maltophilia
Rothia mucilaginosa
Pseudomonas putida
Escherichia coli
Staphylococcus capitis
Fannyhessea vaginae
Corynebacterium pyruviciproducens
Neisseria mucosa
Corynebacterium accolens
Haemophilus parainfluenzae
Atopobium minutum
Atopobium deltae
I–A_unstable
I–B_menses
I–B_unstable
8
6
4
2
0
−2
−4

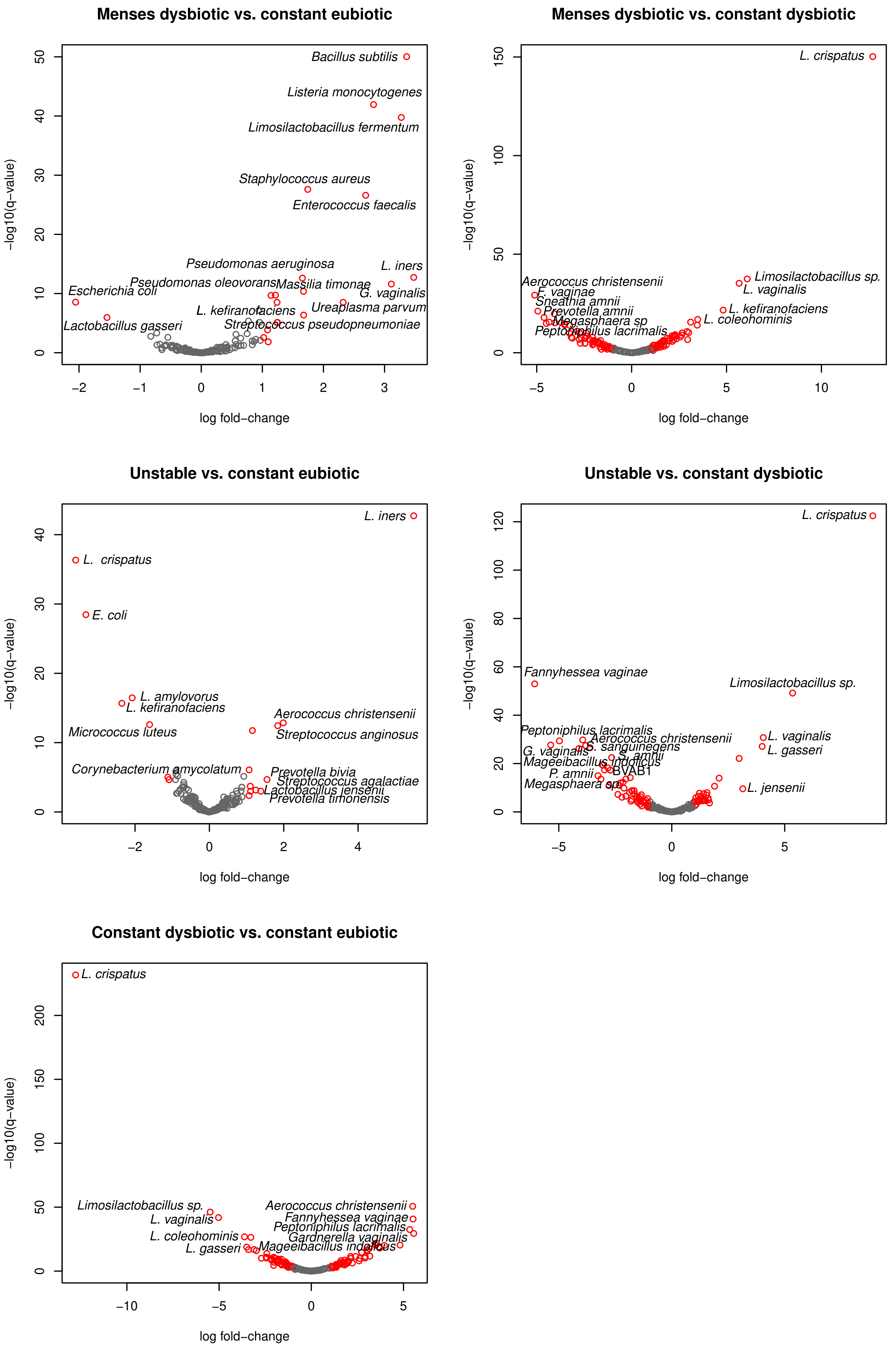
Menses dysbiotic vs. constant eubiotic
Bacillus subtilis
Listeria monocytogenes
Limosilactobacillus fermentum
Staphylococcus aureus
Enterococcus faecalis
Pseudomonas aeruginosa
L. iners
Pseudomonas oleovorans
Massilia timonae
G. vaginalis
Escherichia coli
L. kefiranofaciens
Ureaplasma parvum
Lactobacillus gasseri
Streptococcus pseudopneumoniae
−log10(q−value)
log fold−change
Menses dysbiotic vs. constant dysbiotic
L. crispatus
Aerococcus christensenii
F. vaginae
Sneathia amnii
Prevotella amnii
Megasphaera sp
Peptoniphilus lacrimalis
Limosilactobacillus sp.
L. vaginalis
L. kefiranofaciens
L. coleohominis
−log10(q−value)
log fold−change
Unstable vs. constant eubiotic
L. iners
L. crispatus
E. coli
L. amylovorus
L. kefiranofaciens
Micrococcus luteus
Aerococcus christensenii
Streptococcus anginosus
Corynebacterium amycolatum
Prevotella bivia
Streptococcus agalactiae
Lactobacillus jensenii
Prevotella timonensis
−log10(q−value)
log fold−change
Unstable vs. constant dysbiotic
L. crispatus
Fannyhessea vaginae
Limosilactobacillus sp.
Peptoniphilus lacrimalis
Aerococcus christensenii
L. vaginalis
G. vaginalis
S. sanguinegens
L. gasseri
S. amnii
Mageeibacillus indolicus
P. amnii
BVAB1
Megasphaera sp
L. jensenii
−log10(q−value)
log fold−change
Constant dysbiotic vs. constant eubiotic
L. crispatus
Limosilactobacillus sp.
Aerococcus christensenii
L. vaginalis
Fannyhessea vaginae
Peptoniphilus lacrimalis
L. coleohominis
Gardnerella vaginalis
L. gasseri
Mageeibacillus indolicus
−log10(q−value)
log fold−change

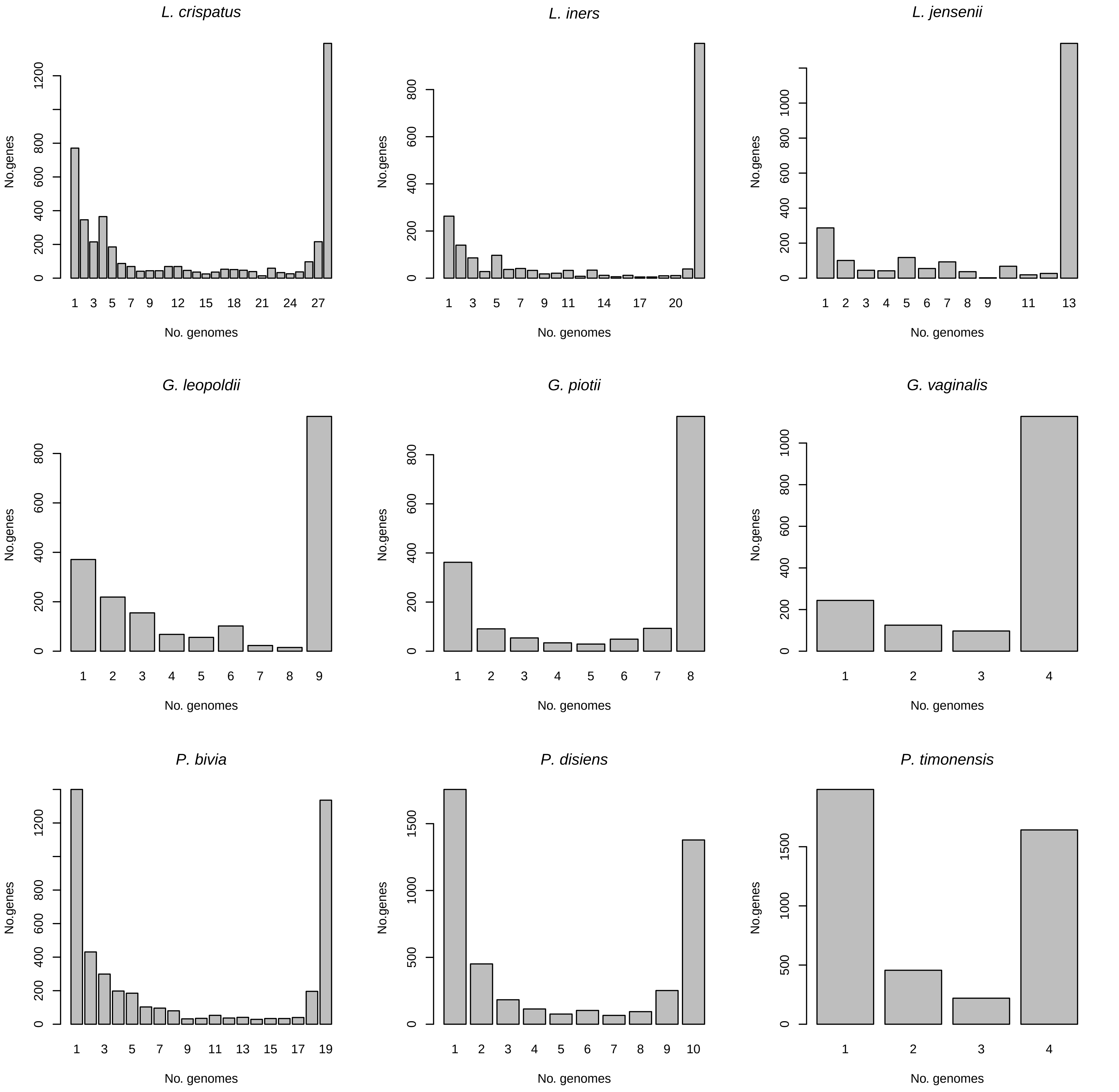

L. crispatus
No.genes
0
200
400
600
800
1200
No. genomes
1
3
5
7
9
12
15
18
21
24
27
L. iners
No.genes
0
200
400
600
800
No. genomes
1
3
5
7
9
11
14
17
20
L. jensenii
No.genes
0
200
400
600
800
1000
No. genomes
1
2
3
4
5
6
7
8
9
11
13
G. leopoldii
No.genes
0
200
400
600
800
No. genomes
1
2
3
4
5
6
7
8
9
G. piotii
No.genes
0
200
400
600
800
No. genomes
1
2
3
4
5
6
7
8
G. vaginalis
No.genes
0
200
400
600
800
1000
No. genomes
1
2
3
4
P. bivia
No.genes
0
200
400
600
800
1200
No. genomes
1
3
5
7
9
11
13
15
17
19
P. disiens
No.genes
0
500
1000
1500
No. genomes
1
2
3
4
5
6
7
8
9
10
P. timonensis
No.genes
0
500
1000
1500
No. genomes
1
2
3
4

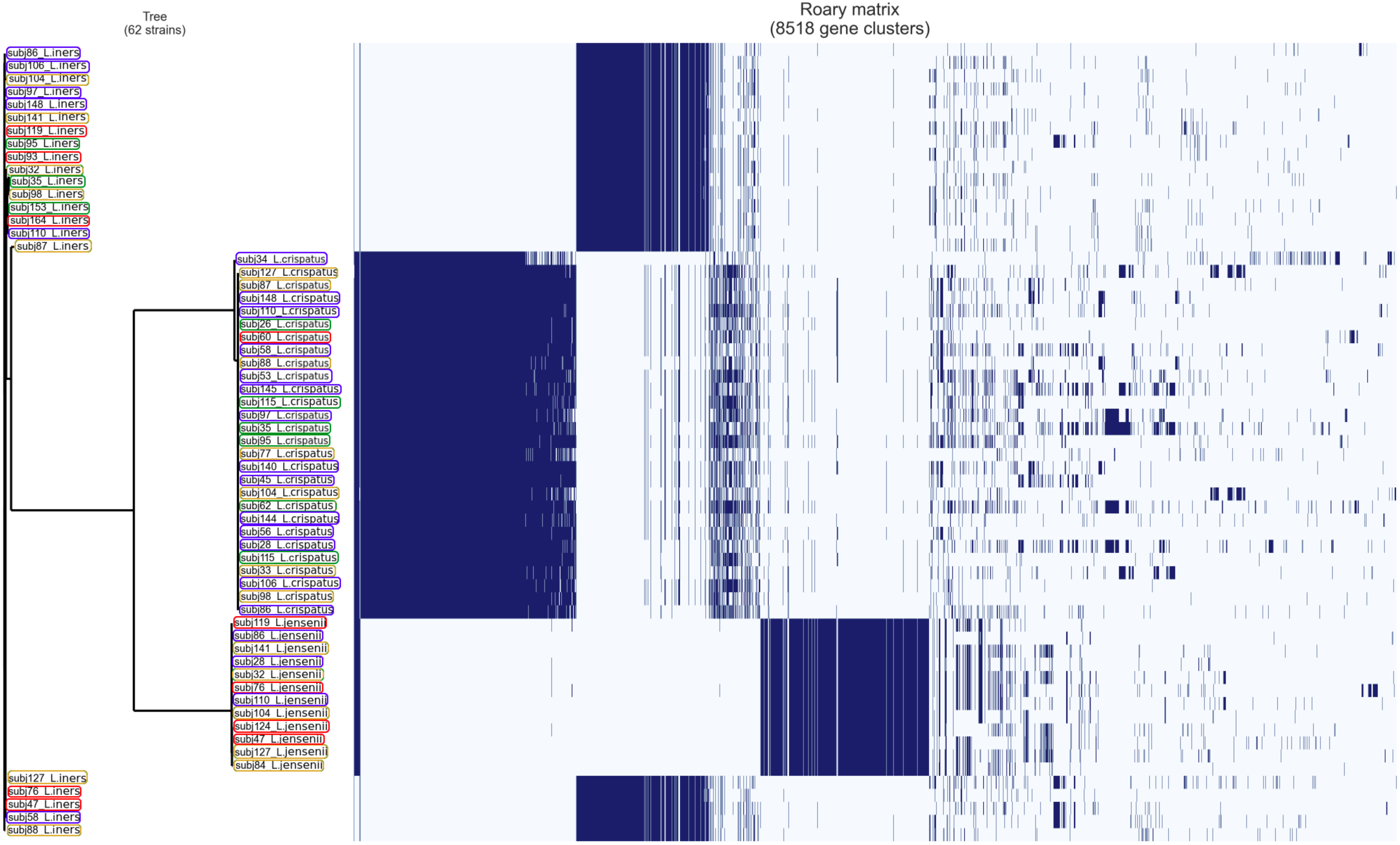

Tree
(62 strains)
Roary matrix
(8518 gene clusters)
subj86_L.iners
subj106_L.iners
subj104_L.iners
subj97_L.iners
subj148_L.iners
subj141_L.iners
subj119_L.iners
subj95_L.iners
subj93_L.iners
subj32_L.iners
subj35_L.iners
subj98_L.iners
subj153_L.iners
subj164_L.iners
subj110_L.iners
subj87_L.iners
subj34_L.crispatus
subj127_L.crispatus
subj87_L.crispatus
subj148_L.crispatus
subj110_L.crispatus
subj26_L.crispatus
subj60_L.crispatus
subj58_L.crispatus
subj88_L.crispatus
subj53_L.crispatus
subj145_L.crispatus
subj115_L.crispatus
subj97_L.crispatus
subj35_L.crispatus
subj95_L.crispatus
subj77_L.crispatus
subj140_L.crispatus
subj45_L.crispatus
subj104_L.crispatus
subj62_L.crispatus
subj144_L.crispatus
subj56_L.crispatus
subj28_L.crispatus
subj115_L.crispatus
subj33_L.crispatus
subj106_L.crispatus
subj98_L.crispatus
subj86_L.crispatus
subj119_L.jensenii
subj86_L.jensenii
subj141_L.jensenii
subj28_L.jensenii
subj32_L.jensenii
subj76_L.jensenii
subj110_L.jensenii
subj104_L.jensenii
subj124_L.jensenii
subj47_L.jensenii
subj127_L.jensenii
subj84_L.jensenii
subj127_L.iners
subj76_L.iners
subj47_L.iners
subj58_L.iners
subj88_L.iners

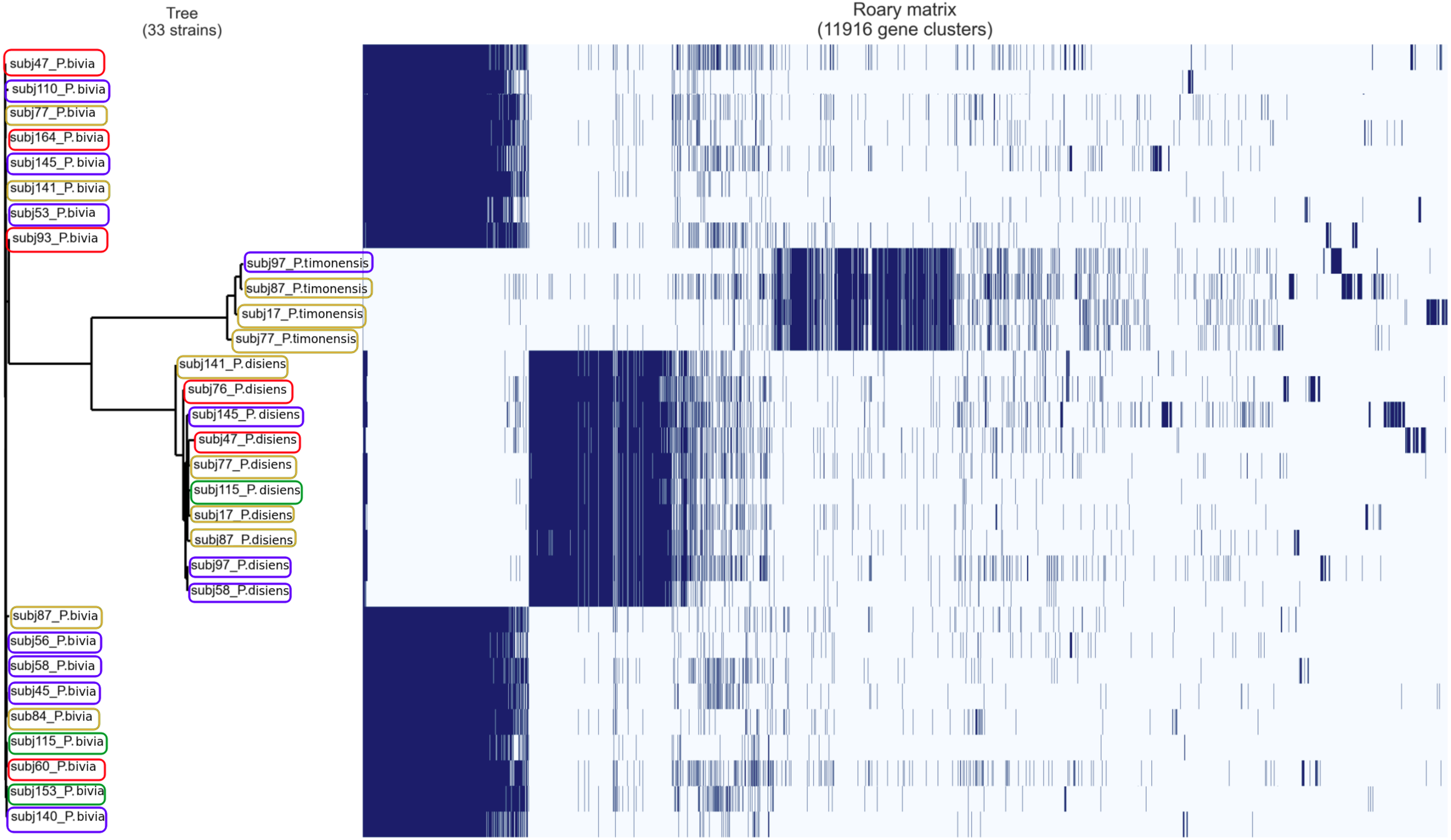

Tree
(33 strains)
Roary matrix
(11916 gene clusters)
subj47_P.bivia
subj110_P. bivia
subj77_P.bivia
subj164_P. bivia
subj145_P. bivia
subj141_P. bivia
subj53_P.bivia
subj93_P.bivia
subj97_P.timonensis
subj87_P.timonensis
subj17_P.timonensis
subj77_P.timonensis
subj141_P.disiens
subj76_P.disiens
subj145_P. disiens
subj47_P.disiens
subj77_P.disiens
subj115_P. disiens
subj17_P.disiens
subj87_P.disiens
subj97_P.disiens
subj58_P.disiens
subj87_P.bivia
subj56_P.bivia
subj58_P.bivia
subj45_P.bivia
sub84_P.bivia
subj115_P. bivia
subj60_P. bivia
subj153_P. bivia
subj140_P. bivia
